# A pathogenic subpopulation of human glioma associated macrophages linked to glioma progression

**DOI:** 10.1101/2025.02.12.637857

**Authors:** Kenny Kwok Hei Yu, Zaki Abou-Mrad, Kristof Törkenczy, Isabell Schulze, Jennifer Gantchev, Gerard Baquer, Kelsey Hopland, Evan D. Bander, Umberto Tosi, Cameron Brennan, Nelson S. Moss, Pierre-Jacques Hamard, Richard Koche, Caleb Lareau, Nathalie Y.R. Agar, Taha Merghoub, Viviane Tabar

## Abstract

Malignant gliomas follow two distinct natural histories: *de novo* high grade tumors such as glioblastoma, or lower grade tumors with a propensity to transform into high grade disease. Despite differences in tumor genotype, both entities converge on a common histologically aggressive phenotype, and the basis for this progression is unknown. Glioma associated macrophages (GAM) have been implicated in this process, however GAMs are ontologically and transcriptionally diverse, rendering isolation of pathogenic subpopulations challenging. Since macrophage contextual gene programs are orchestrated by transcription factors acting on *cis*-acting promoters and enhancers in gene regulatory networks (GRN), we hypothesized that functional populations of GAMs can be resolved through GRN inference. Here we show via parallel single cell RNA and ATAC sequencing that a subpopulation of human GAMs can be defined by a GRN centered around the Activator Protein-1 transcription factor FOSL2 preferentially enriched in high grade tumors. Using this GRN we nominate ANXA1 and HMOX1 as surrogate cell surface markers for activation, thus permitting prospective isolation and functional validation in human GAMs. These cells, termed malignancy associated GAMs (mGAMs) are pro-invasive, pro-angiogenic, pro-proliferative, possess intact antigen presentation but skew T-cells towards a CD4+FOXP3+ phenotype under hypoxia. Ontologically, mGAMs share somatic mitochondrial mutations with peripheral blood monocytes, and their presence correlates with high grade disease irrespective of underlying tumor mutation status. Furthermore, spatio-temporally mGAMs occupy distinct metabolic niches; mGAMs directly induce proliferation and mesenchymal transition of low grade glioma cells and accelerate tumor growth in vivo upon co-culture. Finally mGAMs are preferentially enriched in patients with newly transformed regions in human gliomas, supporting the view that mGAMs play a pivotal role in glioma progression and may represent a plausible therapeutic target in human high-grade glioma.

## Introduction

Malignant gliomas are primary brain cancers characterized by a progressive disease process with two distinct natural histories ^1^: *de novo* glioblastoma (GBM), which follows a rapid and invasive growth pattern, and a low-grade, more indolent tumor that eventually transforms into high-grade disease akin to GBM ^2^. Current glioma classification is based on integrated histological and molecular features ^3–5^, and isocitrate dehydrogenase (IDH) has emerged a key determinant for molecular stratification: IDH wildtype (wtIDH) gliomas correspond to GBM, and the majority of low-grade gliomas (LGG) harbor a mutation in the IDH1 gene. LGG can be further subdivided based on the presence or absence of chromosomal 1p-19q co-deletion ^6^; with 1p19q co-deleted tumors, often with TERT promoter mutations corresponding to oligodendrogliomas, and IDH mutant (mIDH) non-co-deleted tumors astrocytomas, which are associated with mutations in TP53 with ATRX loss.

Human mIDH LGGs have a propensity to progress and transform into higher grades such as Grade 3 and Grade 4. This transformation was initially described as a multi-step process mediated through genomic instability and acquisition of increasing mutation burden ^7–9^, however to date no consistent genomic marker has been shown to be necessary for transformation, and the majority of mutations detected are evolutionary neutral ^10^. Indeed in LGG primary-recurrent sample pairs, transformation occurred frequently despite minimal changes in mutational profile ^11^, suggesting that non-genomic factors likely play a critical role in malignant progression. In this regard, the immune microenvironment has been strongly implicated in glioma progression ^12–14^. Most prominent are glioma associated macrophages (GAM), which consist of resident microglia and bone marrow derived macrophages, constituting up to 50% of the tumor mass, with their presence associated with increasing tumor grade ^12,15^. GAMs are thought to orchestrate a host of pro-tumorigenic functions including angiogenesis, glioma cell migration, invasion and immunosuppression ^12^, however direct ablation or re-programming of GAMs have shown conflicting results in the preclinical setting and global targeting of glioma myeloid cells has failed in clinical trials ^16–18^. One potential explanation is the lack of targeting specificity given the transcriptional diversity found within GAM subpopulations ^19–24^, however identification of robust functional GAM subsets in humans has been challenging, and to date, markers that define pathological GAM are lacking ^25^. Macrophages are highly plastic myeloid cells sensitive to environmental context. Functional specification of macrophages follow a gene regulatory logic based on combinatorial expression and binding of lineage determining and signal dependent transcription factors ^26–28^. We hypothesized that recurrent functional glioma macrophage signatures may be detected through derivation of their underlying gene regulatory network (GRN). To test this, we performed integrated analysis on parallel single-cell RNA and ATAC sequencing data to identify GRNs in a cohort of human surgical samples. Analysis highlighted a subpopulation of GAMs driven by a transcription factor network centered on the transcription factor FOSL2. We demonstrate that this FOSL2 driven subpopulation of GAMs is most abundant in high-grade tumors, irrespective of IDH mutation status, and term these cells malignancy associated GAMs (mGAMs). Through network sensitivity analysis we identify and validate the cell surface marker pairing of Annexin A1 (ANXA1) and Hemoxygenase 1 (HMOX1) as robust markers of the mGAM phenotype that permit prospective isolation of mGAMs for functional analysis, and find that mGAMs are associated with invasion, angiogenesis, extracellular matrix (ECM) protein production and CD4+FOXP3+ T-cell induction. We show that mGAMs share common mitochondrial mutations with bone marrow derived monocytes, and determine the spatial and temporal profile of mGAMs to demonstrate that mGAMs reside within tumor hypoxic niches and induce proliferative mesenchymal transition in IDH mutant glioma cell lines with concomintant expression of CD44 and CHI3L1, which when implanted in vivo accelerate tumor progression. Finally we also demonstrate specific enrichment of mGAMs within transforming regions of low grade glioma.

## Results

### Human GAMs from different conditions share overlapping gene expression and accessibility profiles with inconsistent subcluster membership

GAMs were isolated from surgical samples of IDH wildtype (wtIDH, n=7), IDH mutant gliomas (mIDH, n=8), and normal brain (NB) control samples from the institutional rapid autopsy program (n=2). Tumor mutation profiling ^29^ showed characteristic IDH mutations in the mIDH group, as well as typical GBM mutations such as EGFR, PTEN loss and CDKN2A/B mutants in wtIDH patients (**Figure 1A**). Patient demographic details are listed in **Table 1**. Freshly resected surgical specimens were dissociated into single cell suspensions followed by CD45^+^ magnetic activated cell sorting and profiled using parallel single-cell RNA (scRNA) and single-cell ATAC (scATAC) sequencing. GAM identification and subclustering was based on single-cell gene marker expression ^19^ (**Figure 1B-C**). Cell type proportions were comparable between the two sequencing modalities (**Figure S1A**). The majority of CD45^+^ cells were microglia/macrophages (80.2% in scRNA, 84.64% in scATAC), with a smaller proportion of T-cells (8.43%in scRNA, 6.75% in scATAC) (**Figure 1D**), across all tumor samples. Differential gene expression profiles and gene activity determined by scRNAseq and scATACseq showed that the GAM profiles in mIDH and wtIDH samples show significant overlap (**Figure S1B**), however Gene Ontology (GO) analysis of highly variable genes at the pseudobulk level showed an increase in hypoxia related genes in wtIDH GAMs and an increase in leukocyte migration genes in mIDH GAMs (**Figure 1E**). NB microglia genes clustered separately from the tumor samples (**Figure 1F**).

**Figure 1:**
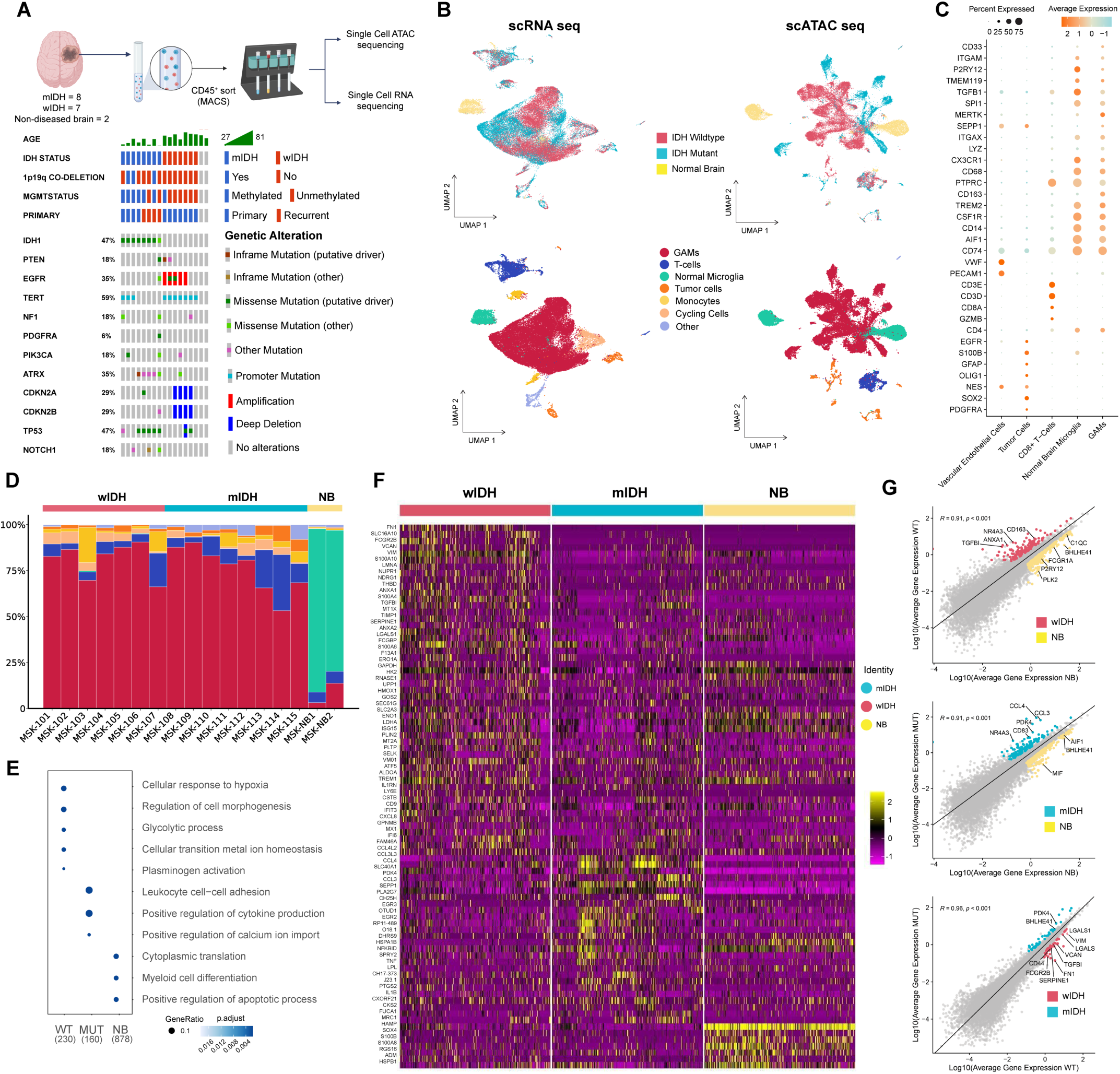
Single cell RNA and ATAC sequencing of GAMs. A) Tissue processing workflow and oncopanel data from a curated set of human specimens. B) UMAPs of scRNA (left panels) and scATAC sequencing (right panels) showing IDH mutation status and cell type distributions. C) Cell type identification using canonical markers for myeloid cells, T-cells, endothelial cells, tumor cells and normal microglia. D) Cell type proportions in different samples analyzed. E) Gene ontology analysis of differentially expressed genes from GAMs in each condition. F) Gene Expression cluster analysis between different conditions. G) Correlation analysis of gene expression between different conditions, note the high degree of correlation between the comparison groups and significant overlap in genes between all three comparison pairs.

**Table 1.**
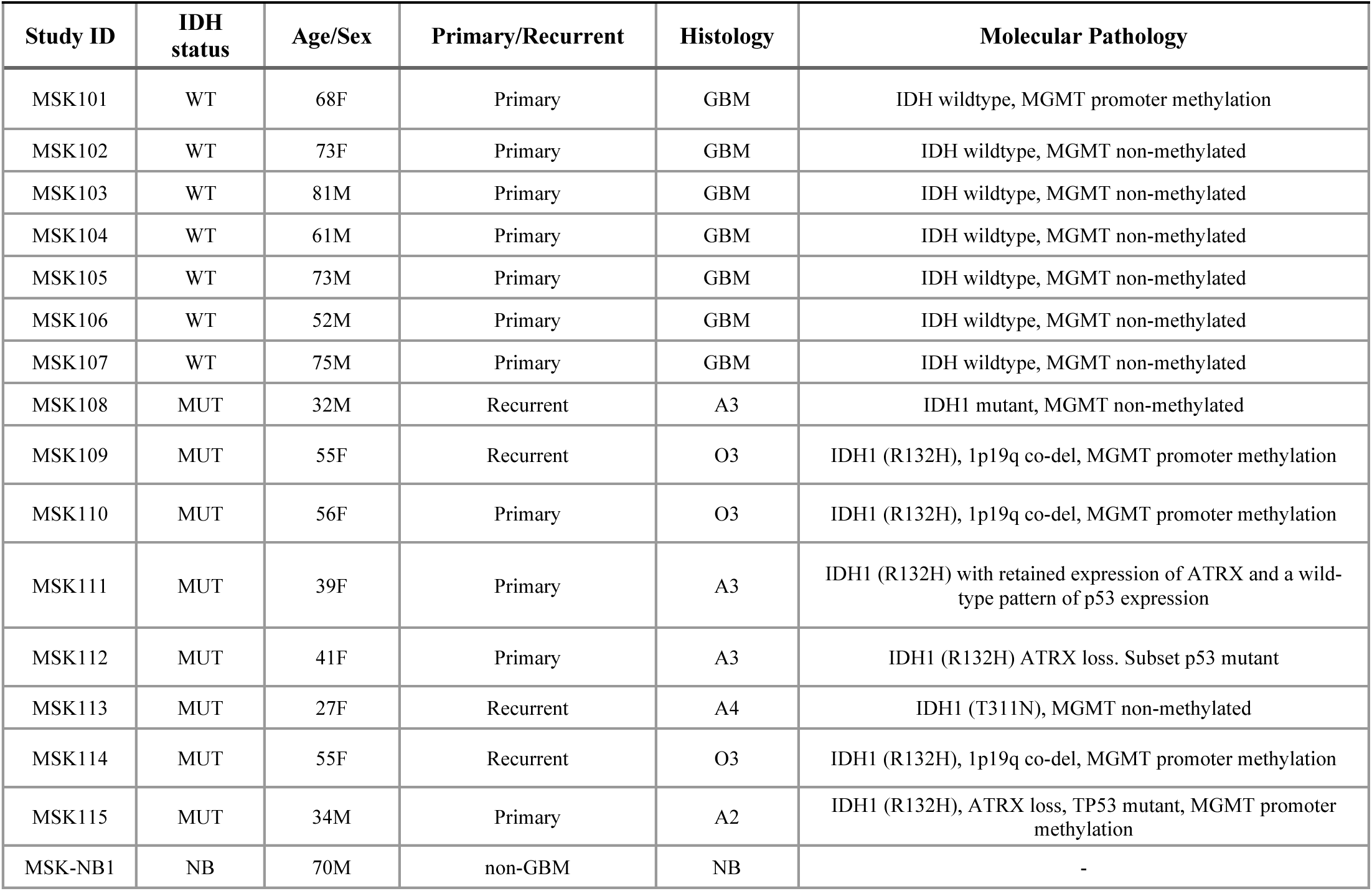
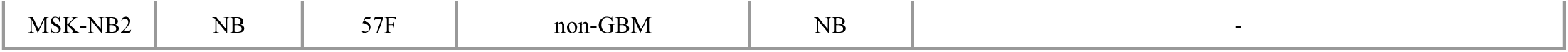
Patient characteristics for samples used for scRNA and scATAC analyses.

**Table 2.** Patient characteristics for samples used for the flow cytometry analyses and BT142 co-culture.

| Study ID | IDH status | Age/Sex | Primary/Recurrent | Histology | Molecular Pathology |
| --- | --- | --- | --- | --- | --- |
| MSK118 | WT | 65F | Primary | GBM | IDH wildtype, MGMT non-methylated |
| MSK119 | MUT | 45M | Primary | A4 | IDH (R132H), ATRX loss, TP53 mutant, MSH6 loss |
| MSK120 | MUT | 26F | Primary | A3 | IDH (R132H), ATRX loss, TP53 mutant |
| MSK121 | MUT | 41M | Recurrent | A3 | IDH (R132C), ATRX loss, TP53 mutant |
| MSK122 | WT | 73M | Primary | GBM | IDH wildtype, MGMT methylated |
| MSK123 | WT | 50M | Primary | GBM | IDH wildtype, MGMT methylated |
| MSK124 | WT | 48M | Primary | GBM | IDH wildtype |
| MSK125 | WT | 74F | Recurrent | GBM | IDH wildtype, MGMT non-methylated |
| MSK126 | WT | 73M | Primary | GBM | IDH wildtype, MGMT methylated |
| MSK127 | MUT | 33F | Primary | A2 | IDH (R132H), 1p19q co-del, TERT, MGMT methylated |
| MSK128 | WT | 79F | Primary | GBM | IDH wildtype, MGMT methylated |
| MSK129 | WT | 56M | Recurrent | GBM | IDH wildtype, MGMT methylated |
| MSK130 | WT | 64M | Recurrent | GBM | IDH wildtype, MGMT non-methylated |
| MSK131 | WT | 53F | Primary | GBM | IDH wildtype, MGMT non-methylated |
| MSK132 | WT | 72F | Recurrent | GBM | IDH wildtype |
| MSK133 | MUT | 45F | Primary | A3 | IDH1 (R132H), ATRX loss, TP53 mutant, MGMT non-methylated |
| MSK134 | WT | 57F | Primary | GBM | IDH wildtype, MGMT methylated |
| MSK135 | WT | 81F | Primary | GBM | IDH wildtype, MGMT methylated |
| MSK136 | WT | 86F | Primary | GBM | IDH wildtype, MGMT methylated |
| MSK137 | WT | 53M | Recurrent | GBM | IDH wildtype, MGMT non-methylated |
| MSK138 | WT | 62F | Primary | GBM | IDH wildtype, MGMT non-methylated |
| MSK139 | WT | 73F | Recurrent | GBM | IDH wildtype, MGMT non-methylated |
| MSK140 | WT | 81M | Primary | GBM | IDH wildtype, MGMT non-methylated |
| MSK141 | WT | 55M | Primary | GBM | IDH wildtype, MGMT non-methylated |
| MSK142 | WT | 61F | Primary | GBM | IDH wildtype, MGMT non-methylated |
| MSK143 | WT | 66M | Recurrent | GBM | IDH wildtype, MGMT non-methylated |
| MSK144 | WT | 78F | Primary | GBM | IDH wildtype, MGMT methylated |
| MSK145 | MUT | 25M | Recurrent | A2 | IDH2 (R132S), ATRX loss, TP53 mutant, NF1 |
| MSK146 | MUT | 28M | Primary | A4 | IDH2 (R132G), ATRX loss, TP53 mutant |
| MSK147 | MUT | 40M | Recurrent | A3 | IDH2 (R172S), ATRX loss, TP53 mutant |
| MSK148 | MUT | 44M | Recurrent | A4 | IDH1 (R132H), ATRX loss, TP53 mutant, MGMT promoter methylation |
| MSK149 | WT | 75F | Primary | GBM | IDH wildtype, MGMT non-methylated |
| MSK150 | WT | 64M | Primary | GBM | IDH wildtype, MGMT methylated |
| MSK151 | WT | 79M | Primary | GBM | IDH wildtype, MGMT methylated |
| MSK152 | WT | 60M | Recurrent | GBM | IDH wildtype, MGMT non-methylated |
| MSK153 | WT | 49M | Recurrent | GBM | IDH wildtype, MGMT non-methylated |
| MSK154 | WT | 74M | Recurrent | GBM | IDH wildtype, MGMT methylated |
| MSK155 | WT | 69F | Recurrent | GBM | IDH wildtype, MGMT methylated |
| MSK156 | MUT | 42M | Recurrent | A3 | IDH1 (R132H), ATRX loss, TP53 mutant, MGMT promoter methylation |
| MSK157 | WT | 60F | Recurrent | O3 | IDH (R132H), 1p19q co-deleted |

Furthermore, mIDH and wtIDH GAM genes showed a high degree of correlation in expression, with only a minority of genes differentially expressed in either condition (**Figure 1G**): These genes included PDK4 and BHLHE41 in mutant samples, and VIM, CD44, LGALS1, VCAN, and TGFB1 in the wtIDH GAMs. There was increased expression of chemokine genes such as CCL3 and CCL4 in mIDH GAMs compared to normal brain microglia, and an increased expression of CD163, ANXA1, NR4A3 and TGFB1 in wtIDH GAMs compared to NB microglia. Analysis of inferred gene activity based on scATAC ^30^ corroborated scRNA results and again did not show significant differences between mIDH and wtIDH GAMs (**Figure S1B**). Overall, mIDH and wtIDH GAMs share substantial similarities in both gene expression and gene accessibility, highlighting the need to better define GAM subpopulations within distinct glioma subtypes.

#### Analysis of subclusters in GAMs reveals inconsistent cluster membership

Unsupervised Louvain clustering of the combined populations of mIDH, wtIDH and NB immune cells resulted in 9 clusters (**Figure 2A**). To determine if global differences in gene expression between mIDH and wtIDH GAMs were driven by cluster-specific cells, we determined the expression of genes differentially upregulated in wildtype GAMs such as VIM, CD44 and LGALS1 in each cluster. We found that expression of these genes was frequently distributed across multiple clusters, across both wtIDH and mIDH GAMs (**Figure S2A**). Prior studies have proposed that these subclusters correspond to GAM subtypes ^24^, however bioinformatically assigned cluster membership may not reflect true biological activity ^31^, and may be susceptible to batch effect and biases in data generation. We therefore compared the consistency of cluster assignments by performing label transfer from two published datasets: one study by Abdelfattah et al. ^24^ which contained a combination of mIDH and wtIDH GAM samples and a second larger aggregated GBM only dataset (GBMap) ^32^. These two datasets allowed us to evaluate the consistency of cell type labelling and the robustness of cluster membership by examining cell type assignments. In most cases, label transfer from these two reference datasets showed highly variable labelling of cells from the same initial cluster assignment, depending on which reference dataset the labels were taken from (**Figure 2B**). In contrast, certain transcriptionally distinct cell types such as monocytes were consistently labelled as a distinct group in both reference datasets (**Figure 2B**). Overall, cell type assignments based on consensus clustering of larger aggregated datasets such as GBMap generated more biologically plausible results. For example, nearly all NB microglial cells mapped to degenerative microglia in GBMap ^32^. Similarly, the TAM-Mg subgroup was more enriched in mIDH GAMs, and the TAM-BDM/Hypoxia MES subgroup was more enriched in wtIDH GAMs (**Figure 2C**), consistent with what has previously been reported ^21,33,34^. Together, these findings favor the use of larger aggregated datasets, but highlight inconsistencies in subcluster assignment using standard clustering approaches.

**Figure 2:**
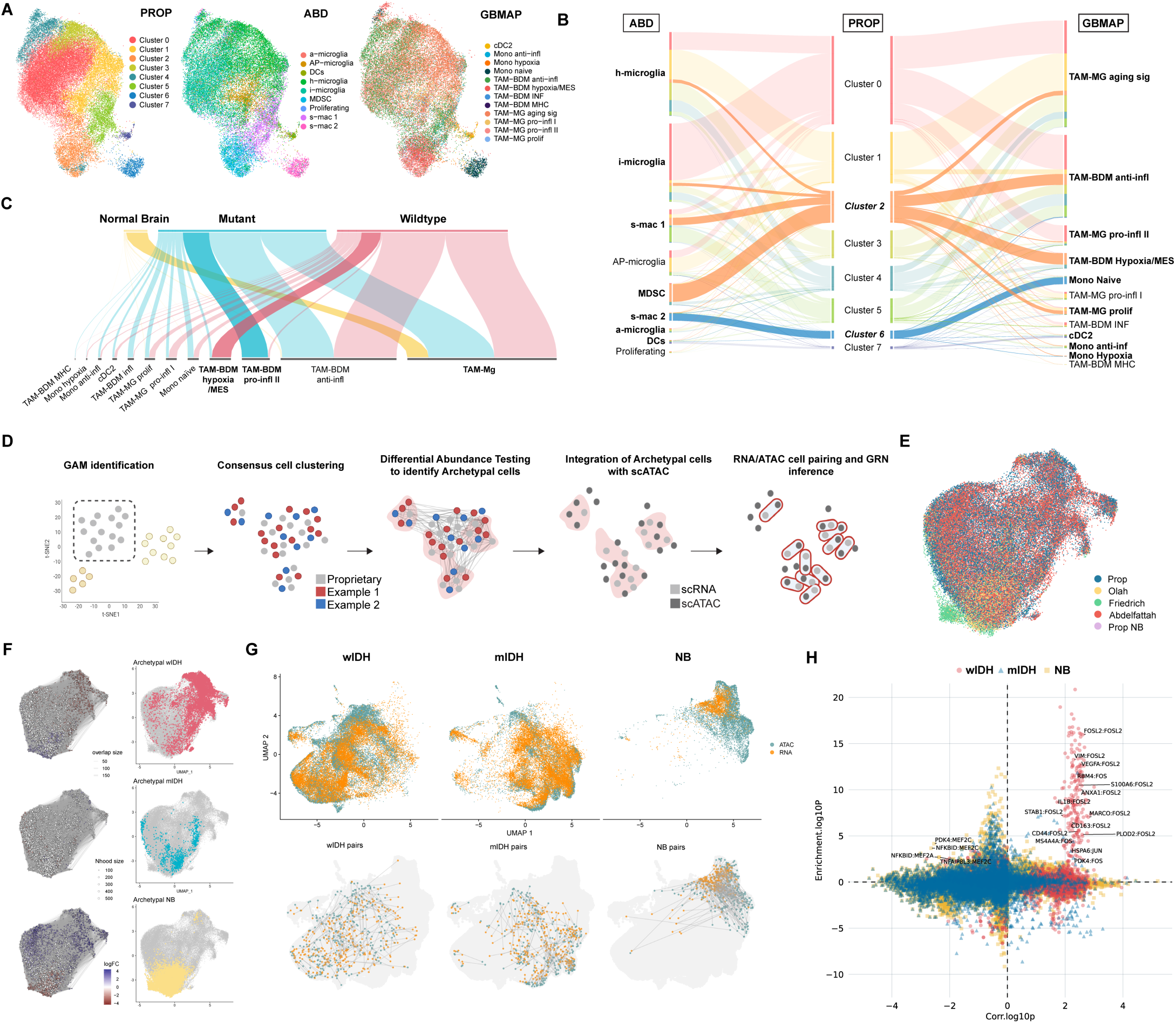
Gene Regulatory Network Inference in GAMs. A) Louvain clustering of proprietary dataset and label transfer of subcluster labels from Abdelfattah 2022 ^24^ (middle), and GBMap ^32^ (right). B-C) Cluster label identification for the same cells using different reference datasets. Cells labelled as Cluster 2 using Louvain clustering (middle column) are assigned different cell types depending on the reference dataset. Some cell types (e.g. Cluster 4/monocytes) are consistently labelled. D) Schematic representation of computational pipeline. GAMs are first identified in scRNA data then combined with additional public datasets (Examples 1&2) before performing differential abundance testing to identify archetypal cells. Archetypal cells from the original dataset are then matched to their scATAC counterpart and GRN inference is performed on these “pseudo-pairs”. E) UMAP projection of combined proprietary and public datasets. F) Differential abundance testing to identify archetypal cells. G) scRNA-scATAC pairing (Optmatch) ^38^ H) GRN inference on archetypal cells (FigR) ^38^.

### Gene Regulatory Network inference highlight FOSL2 as significant transcription factor network in high-grade gliomas

#### Consensus clustering of GAMs and differential abundance testing identifies archetypal myeloid cell states

Given these findings, we integrated our scRNA datasets with multiple public datasets that contained a mixture of high-grade and low-grade, mIDH and wtIDH gliomas ^24,33^ and NB microglia ^35^ using the *Harmony* implementation within *Seurat* ^36^ (**Figure 2D, E-G**). We then applied label-free differential abundance testing using *MiloR* ^37^, placing all cells within a K-nearest neighbor graph modelling cellular states as overlapping neighborhoods, before performing differential abundance testing. This resulted in an enrichment of modelled cell states within each specified condition, which we term “Archetypal cells”. Archetypal cells made up 19.0% percent of wtIDH GAM cells, 4.5% of mIDH GAM cells and 17.3% of normal brain microglia within our consensus dataset (**Figure S2B**). Repeat differential gene expression analysis using *Deseq2* now demonstrated much clearer differences and separation between archetypal cell types (**Figure S2C**).

#### scRNA-scATAC integration and cell pairing and GRN construction identify an archetypal GAM signature driven by FOSL2

We next integrated our parallel scRNA and scATAC sequencing dataset through dimensionality reduction and co-embedding, then applied the optimal full matching algorithm *Optmatch* ^38^ to computationally pair cells from our scRNA and scATAC sequencing datasets to form “pseudopairs”. Cell pairings were filtered by pairs containing archetypal scRNA cells and secondary gene regulatory network inference was performed using *FigR* ^38^ (**Figure 2D**). *FigR* defines Domains of Regulatory Chromatin (DORC) as regions of accessibility and describes transcription factor (TF)–target gene relationships. DORCs significantly enriched in wtIDH samples included [FOSL2:CCL3], [FOSL2:VIM], [FOSL2:MARCO] and [FOSL2:VEGFA]. Whereas in mIDH samples enriched DORCS included [TAL1:SLC1A3], [TAL1:SALL1] and [KLF2:CD74] (**Figure 2H**). Defining individual DORCs allowed the construction of TF-networks and generation of GRNs specific for each archetype (**Figure 3A**). Wildtype archetypal cells consist of four main TF networks including FOSL2, FOSB, NR4A3 and CXXC5. MUT archetypal cells consist of TAL1, REL, and KLF2 TF networks and Archetypal NB microglial cells had ETS2 and FOXP2. As a separate validation we applied *ChromVAR* ^30^ to our scATAC data to confirm accessible gene-TF binding sites for each condition (**Figure S2D**). This method corroborated the differences in archetypal accessibility and confirmed that the accessibility differences observed were not secondary to dissociation artifact. Analysis of DORCs identified FOSL2 as a transcription factor consistently enriched in the wildtype GAM archetype. We then inferred FOSL2’s target genes, also known as its *regulon*, through DORC enrichment. We collated target genes which had a regulation score above 0.5 (“FOSL2 large regulon”, 248 genes) and further filtered genes based on their expression level and regulation score to derive a smaller, more focused targeted gene set (referred to as “FOSL2 regulon” from now, 54 genes). Constituent members of the FOSL2 regulon included genes such as VIM, S100A6, ANXA1, ANXA2, CD44, TGFB1 sharing some overlap with a mesenchymal signature previously described in both glioma cells and GAMs ^34^, associated with epithelial-mesenchymal transition (EMT) and extracellular matrix (ECM) remodeling. GO analysis showed that the top GO biological processes were cellular response to hypoxia (GO:0036294), and regulation of angiogenesis (GO:0045765) (**Figure 3B**). We determined the cell type specificity of the FOSL2 regulon and evaluated cell populations within GBMap ^32^. This analysis confirmed that the FOSL2 regulon was predominantly expressed in myeloid cells and not tumor cells, even though the transcription factor FOSL2 itself was also expressed in tumor cells (**Figure S2E**). Since FOSL2 is a member of the Activator Protein 1 (AP-1), we asked if the FOSL2 regulon was part of a generic acute stress response and identified genes from the Integrated Stress Response (ISR) and immediate early genes (IEG) sets and found minimal overlap (**Figure S3A**). We also asked if hypoxia alone was sufficient to induce this signature. Since the control microglial cells were obtained from autopsy samples and would have been exposed to some degree of hypoxia, we analyzed NB microglial cells with high degree of HIF1A expression as a cross validation and found minimal overlap with the FOSL2 regulon (**Figure S3B**). Taken together these data suggest that archetypal GAMs in IDH wildtype gliomas have high FOSL2 regulon expression, express genes associated with hypoxic response, ECM remodeling and angiogenesis, but are distinct from a generic AP-1 stress response and have a transcriptome distinct from normal microglial cells exposed to hypoxia.

**Figure 3:**
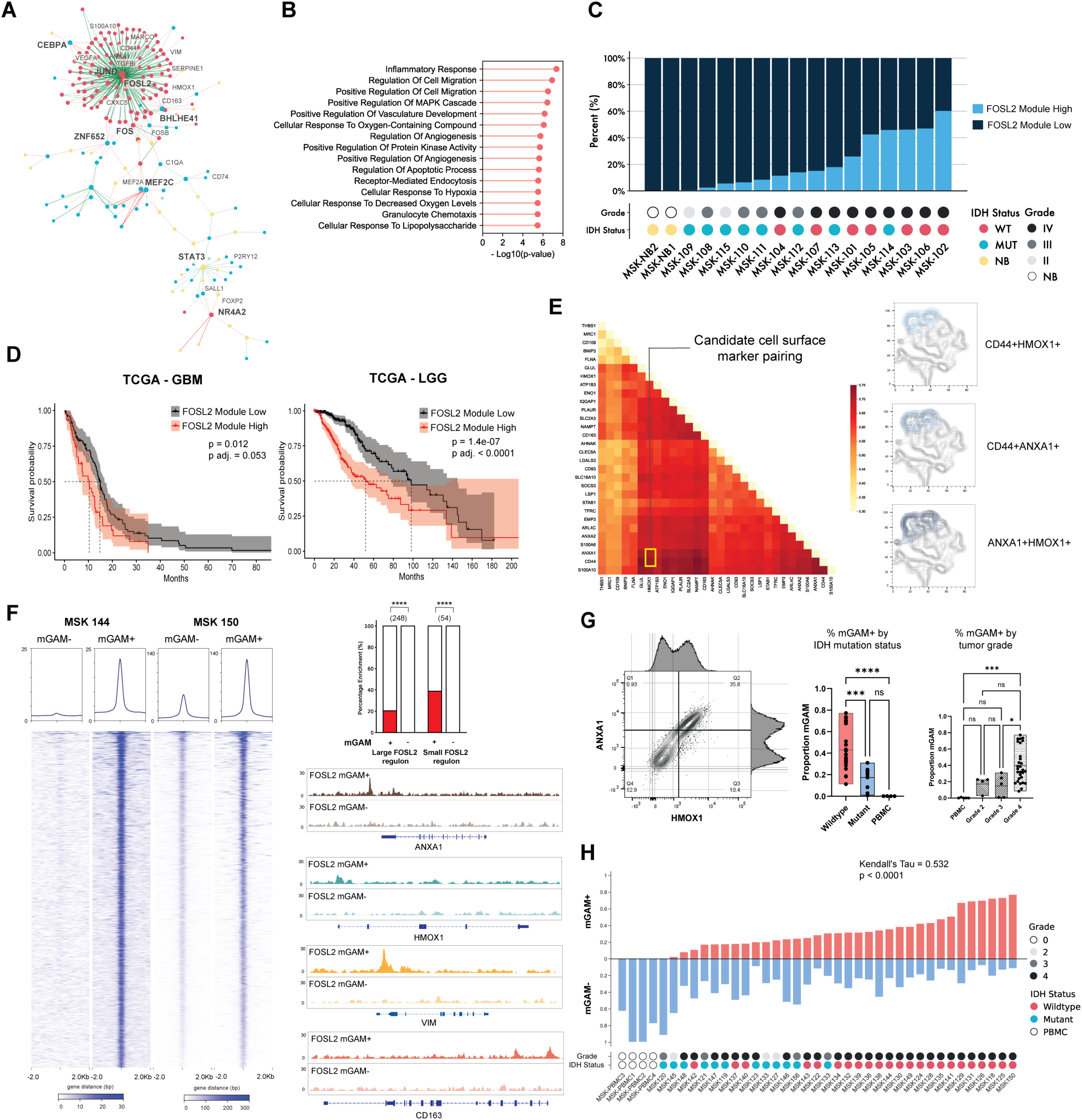
GRN analysis nominates HMOX1 and ANXA1 as cell surface markers of mGAMs. A) GRN inference for mIDH and wtIDH GAMs and NB microglia. B) Gene Ontology terms within the FOSL2 regulon. c) GAMs from all single-cell analyzed samples binarized according to FOSL2 module score “High” versus “Low”, plotted alongside corresponding grade and IDH mutation status. Each patient sample is assigned an MSK number (MSK#). D) Survival analysis using TCGA data for GBM dataset (left), and LGG dataset (right). E) Network Sensitivity Analysis showing top marker pairs for cell surface marker genes within the FOSL2 regulon (left). tSNE plot of flow cytometric data confirming top marker pairs ANXA1:CD44 and HMOX1:CD44 and ANXA1:HMOX1 label the same population of cells. F) ChIP-seq data of mGAMs versus non-mGAM populations showing enrichment of FOSL2 protein binding at predicted sites in the mGAM population (left and top right). Proportion of genes with FOSL2 binding in mGAM positive versus negative populations. Left barchart shows large FOSL2 regulon, right barchart shows small FOSL2 regulon, number of genes within regulon shown in brackets. Representative tracks showing FOSL2 ChIPseq peaks at ANXA1 and HMOX1, as well as VIM and CD163 (bottom right). G) Flow cytometry plot showing presence of two populations and increased proportion of mGAMs in high-grade tumors and IDH wildtype tumors. H) Waterfall plot of patients analyzed/flow sorted using the ANXA1/HMOX1 mGAM markers (top), showing strength of correlation between mGAM % and grade (Kendall’s tau statistic = 0.532, p<0.0001).

### Archetypal GAMs are present in high-grade gliomas irrespective of IDH mutation status

To assess FOSL2 regulon expression, we assigned cells a FOSL2 regulon module score and binarized cells into FOSL2 regulon score High versus Low (**Figure S2F-G**). Interestingly, certain mIDH samples had distinctly higher proportions of FOSL2 regulon High cells, which corresponded to histologically aggressive grade 4 mIDH astrocytomas. We therefore reasoned that the FOSL2 archetypal cell enrichment may in fact be correlated with grade rather than IDH mutation status (**Figure 3C**); remarkably we also found that expression of the FOSL2 regulon was more closely aligned with grade than with IDH mutation status, suggesting an association with disease progression. Since GAM infiltration and the mesenchymal subtype of GBM have been associated with a worse prognosis ^15,39,40^, we also examined the GBM subtypes in our samples. The degree of enrichment of the FOSL2 regulon did not correlate with a specific GBM mutational subtype ^4^ (**Figure S3C**), suggesting a more general phenomenon. We also applied the FOSL2 regulon analysis to TCGA data and charted survival differences in the GBM ^3,4^ and LGG dataset and found a significant survival difference in the LGG group, with an increased level of FOSL2 regulon expression associated with a poorer prognosis (p adj. < 0.01), and a trend towards significance within the GBM group (p adj. = 0.053) (**Figure 3D**). Given the association with higher grade tumors, we termed this subgroup of GAMs driven by the FOSL2 regulon malignancy associated GAMs (mGAM).

### Prospective identification of ANXA1 and HMOX1 as markers of the mGAM phenotype

We next searched for cell surface markers which could allow for prospective isolation of mGAMs from patient samples for functional evaluation. Using a combinatorial classification algorithm (see Methods) based on scRNA data, we identified two-marker cell surface protein combinations with high levels of specificity and sensitivity for isolating mGAMs. Top gene pairs were ones containing either CD44, ANXA1, or S100A10, specifically when combined with any of the following nine genes: GLUL, HMOX1, ATP1B3, ENO1, IQGAP1, PLAUR, SLC2A3, NAMPT, and CD163 (**Figure 3E**). Although CD44 was noted to be frequently represented in the top pairings, the ubiquity of CD44 expression in different cell types in glioma ^41^ led us to select ANXA1 and HMOX1 as our candidate pairing with confirmed CD44 co-expression (**Figure 3E right**). Using these cell surface markers for flow sorting, we next validated FOSL2 DNA binding sites in mGAM and non-mGAM cells using low input Chromatin Immunoprecipitation sequencing (ChIPseq). We found that FOSL2 binding was globally decreased in the non-mGAM samples as compared to the mGAM population at the predicted sites in both cases (**Figure 3F**), and there was a high degree of overlap between the predicted small FOSL2 regulon (21 out of 54 genes, 38.9% overlap, and the larger regulon (46 out of 248 genes, 18.5% overlap). In contrast none of the genes in either the large or small regulon were noted to have peak enrichment in the non-mGAM samples (p = < 2.2e-16, odds ratio = 6.63) (**Figure 3F right**). These data therefore provide molecular support for differential FOSL2 binding in mGAM and non-mGAM populations, validating our computationally derived GRN predictions.

### ANXA1 and HMOX1 surface expression correlates with clinical grade in a spectrum of glioma patient samples

We next applied mGAM markers to CD45^+^ cells isolated from a separate cohort of mIDH and wtIDH surgical samples as well as peripheral blood samples from non-glioma patients. The proportion of mGAMs, defined as ANXA1^+^HMOX1^+^ immune cells, was higher in high-grade tumors in both the wtIDH and mIDH samples, with negligible proportions found in PBMCs (**Figure 3G-H**). Corroborating the findings from the prior single-cell sequenced cohort, the proportion of mGAMs within the GAM population was not associated with a specific mutational profile (**Figure S3D**). We conclude that the surface marker pair of ANXA1 and HMOX1 identifies cells differentially enriched for FOSL2 binding at gene promoter and enhancer sites associated with the FOSL2 regulon, and the percentage of mGAMs, as defined by cell surface protein markers, is more strongly correlated with grade than IDH status (Kendall tau statistic = 0.532 vs 0.493, p < 0.0001 and p < 0.001 respectively). Since CD45^+^ sort may also include non-myeloid cells, flow sorted mGAMs were also re-analyzed for ANXA1, HMOX1 and myeloid marker expression on a separate flow cytometer to confirm myeloid-exclusive expression of mGAM markers (**Figure S3E**).

### Primary human mGAMs are multi-functionally pro-tumorigenic

To dissect the function of mGAMs, we performed a series of assays focused on presumed function of the FOSL2 regulon genes, including secretion of extracellular matrix proteins such as vimentin, invasive capacity and angiogenesis. mGAMs were placed in short term culture (**Figure 4A**), with supernatant then analyzed for ECM proteins such as vimentin and versican. These proteins, predicted to be upregulated within the FOSL2 regulon, showed a differential increase in the mGAM population (p = 0.0055 and p = 0.0321 respectively) (**Figure 4A**). Angiogenesis, measured using the tube forming assay in human umbilical vein endothelial cells (HUVEC), was also significantly increased in the mGAM population (p < 0.0001) compared to non mGAMs (**Figure 4C**).

**Figure 4:**
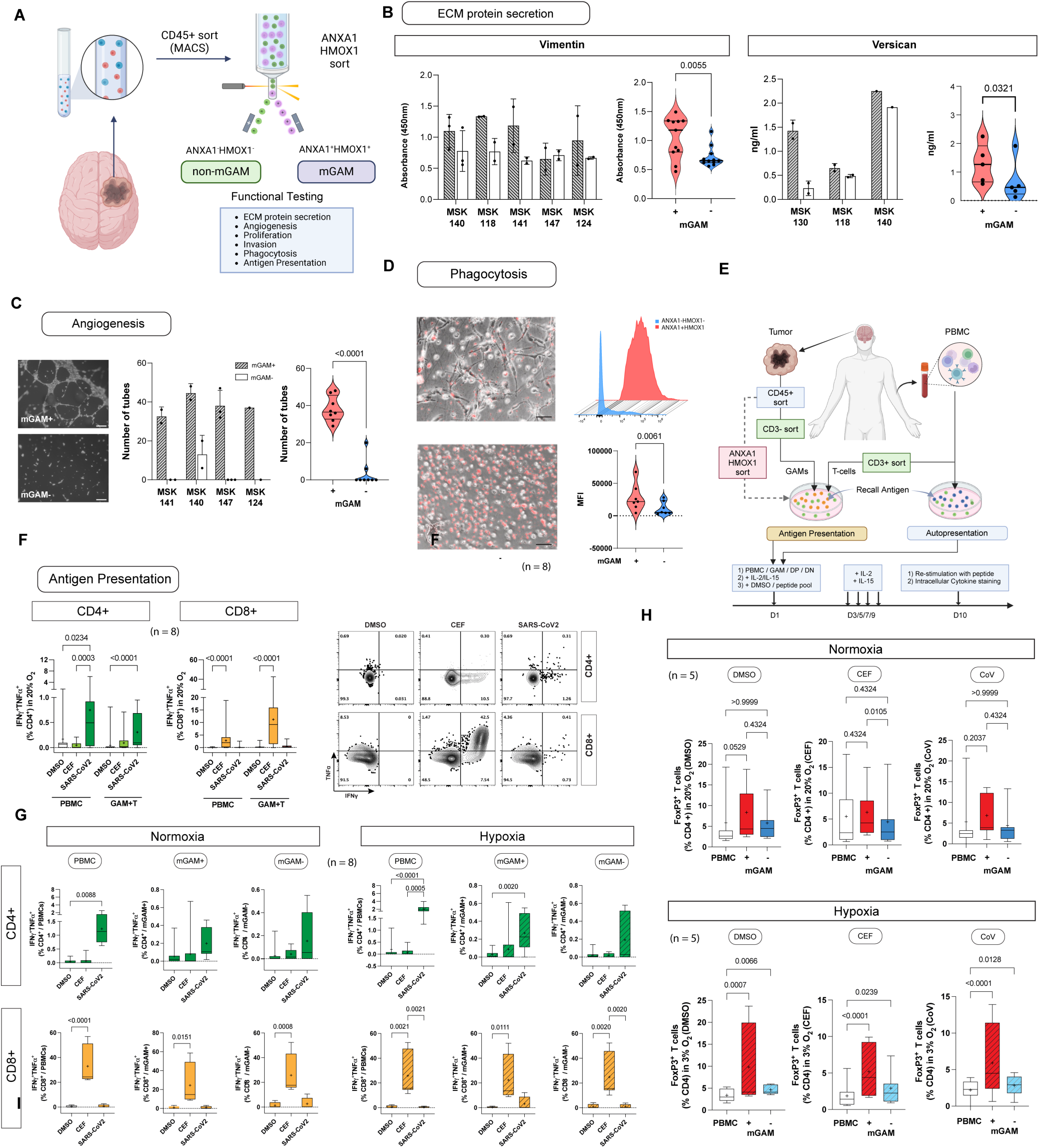
mGAMs display pro-tumorigenic functions. A) Flow sorting strategy to isolate mGAMs and non-mGAMs for functional testing. B) Extracellular matrix protein secretion (Vimentin and Versican) measurements in mGAM and non-mGAM supernatant using ELISA. C) Angiogenesis tube forming assay using HUVEC (scalebar = 100µm). Left) representative calcein AM stained images for HUVECs cultured with mGAM versus non-mGAM supernatant at 24 hours. D) Phagocytosis assay using pH-rhodo E.coli particles showing increased phagocytosis capacity in mGAMs (scalebar = 50µm). E) Autologous Antigen Recall Assay Experimental Design. F) T cell responses in a recall antigen assay targeting MHC Class I and Class II receptors in PBMC controls versus unsorted GAMs. G) Antigen recall assay in primary GAMS sorted for mGAM versus non-mGAMs cultured under normoxic and hypoxic conditions. H) mGAM versus non-mGAM T-cell co-culture followed by T-cell phenotyping demonstrating significant increase in CD4+FOXP3+ T cells when T-cells are co-cultured with mGAMs under hypoxia.

### mGAMs are capable of presenting antigen and induce FOXP3^+^ T-cell expansion under hypoxia

Given GAMs are phagocytes we sought to determine their antigen presentation properties. We first measured phagocytosis capacity using the pH-rhodo assay. mGAMs exhibited consistent increased phagocytic capacity compared to their non-mGAM counterparts (n=8) as measured by pH-rhodo E.coli particle ingestion (**Figure 4D**). We then explored whether this difference in phagocytic function was associated with differences in antigen processing and presentation. GAMs are innate immune cells that should theoretically be capable of antigen presentation; however, functional demonstration of effective antigen presentation has not been previously reported in primary human GAMs. To probe the ability of both GAMs and mGAM to present antigen, we took advantage of prior vaccination for SARS-CoV2 in our patient population to collect autologous patient tumor and blood. We designed an antigen recall assay to assess the ability for GAMs and mGAMs to elicit polyfunctional (TNF/IFNγ) T-cell responses via known MHC Class I and II recall antigens (**Methods**, **Figure 4E**). Briefly, patient PBMCs were exposed to a known recall antigen – in this case the SARS-CoV2 15-mer spike protein, as well as CMV, EBV and Influenza proteins (CEF), to assess CD4^+^ and CD8^+^ T-cell polyfunctional response to antigen stimulation respectively. An auto-presentation assay using the mixed lymphocyte culture was used as an autologous control to confirm responses to MHC class I and class II antigens. In the first cohort of patients, the antigen recall assay was performed directly on patient GAMs (n=8). GAMs were capable of antigen presentation to CD4 and CD8 T-cells to produce polyfunctional T-cell responses, with some patient variability (**Figure 4F**). We next performed the antigen recall assay in freshly sorted mGAMs versus non-mGAMs under normoxic (20%) and hypoxic (3%) conditions to mimic in vivo tumor environment (**Figure 4G**). Here again both mGAMs and non-mGAMs exhibited robust MHC Class II responses to SARS-CoV2 protein in CD4 cells and MHC Class I responses to CEF cocktail in CD8 cells, with no significant differences in T-cell responses between mGAMs and non-mGAMs with or without hypoxia (**Figure 4G**). Immunophenotyping of T-cells which were co-cultured with mGAMs and non-mGAMs however showed a significant increase in FOXP3^+^ proportions in CD4^+^ T-cells co-cultured with mGAMs, with the effect particularly marked under hypoxia (**Figure 4H-I**). This effect was independent of the type of antigen used.

Taken together these data suggest that mGAMs are capable of antigen presentation but also induce expansion of CD4^+^FOXP3^+^ T cells, thought to represent immature T-regulatory cells, through an antigen independent pathway under hypoxic conditions. Collectively these data demonstrate a coherent set of pro-tumorigenic functional changes differentially found in mGAMs.

### mGAMs share somatic mitochondrial mutations with peripheral blood monocytes

We next asked if mGAMs could be bone marrow derived sharing common ontogeny to blood borne monocytes. mGAMs share transcriptional similarity to hypoxic monocytes (**Figure S6A**), and co-culture of primary human monocytes with hypoxia and glioma cells partially recapitulated the mGAM phenotype, whereas hypoxia alone did not (**Figure S6B-E**). Proof however that mGAMs are of blood-borne origin remained lacking. We therefore applied a mitochondrial DNA single-cell ATAC-seq (mtscATAC-seq) protocol ^42^ in mGAMs and non-mGAMs isolated from clinical samples as well as matched PBMCs in a separate set of patient samples (n=2; **Figure 5A**). We first validated the mGAM chromatin accessibility profile in these samples against that of our computationally defined FOSL2 GRN and confirmed that the isolated mGAMs had a higher concordance with the FOSL2 regulon accessibility profile than non-mGAMs (**Figure 5B-C**). This analysis further showed that monocytes had the highest concordance with the FOSL2 chromatin accessibility profile (**Figure 5C**). As naïve monocytes do not express ANXA1 and HMOX1 at baseline and have a gene expression profile that is distinct from mGAMs, we interpret this epigenetic signature to indicate that monocytes may be “primed” to differentiate into mGAMs depending on the environmental context. Using the paired mitochondrial DNA profiles from the mtscATAC-seq libraries ^43^, we sought to determine if mGAMs could be traced back to existing peripheral blood populations from two patient samples: a transformed low-grade astrocytoma and a wtIDH GBM (see Methods). We hypothesized that if mGAMs were derived from peripheral blood or bone marrow, then we should see a significant proportion of blood-borne cells within the mGAM population. Indeed, heteroplasmy analyses of 650 high-confidence mutations showed a strong concordance of allele frequencies between mGAMs and monocytes, suggesting considerable clonal overlap between the populations, including highly heteroplasmic variants like m.8780T>C and 9253G>A (**Figure 5D-E**). Further, the proportion of cells carrying a high-confidence mtDNA variant from monocytes was higher in mGAMs (37.3%) than non-mGAMs (34.5%; **Figure 5F**). Collectively, these analyses suggest an enrichment of blood-borne myeloid cells in the mGAM population though we cannot exclude the possibility that resident microglia could also adopt an mGAM phenotype. Given the possibility that mGAMs might be bone marrow derived and the fact that ANXA1 can be induced by dexamethasone, a drug commonly administered to patients with gliomas, we also analyzed mGAM markers in peripheral blood from patients receiving dexamethasone peri-operatively and confirmed the exclusive expression of mGAM markers within the GAM population (**Figure S6F**).

**Figure 5:**
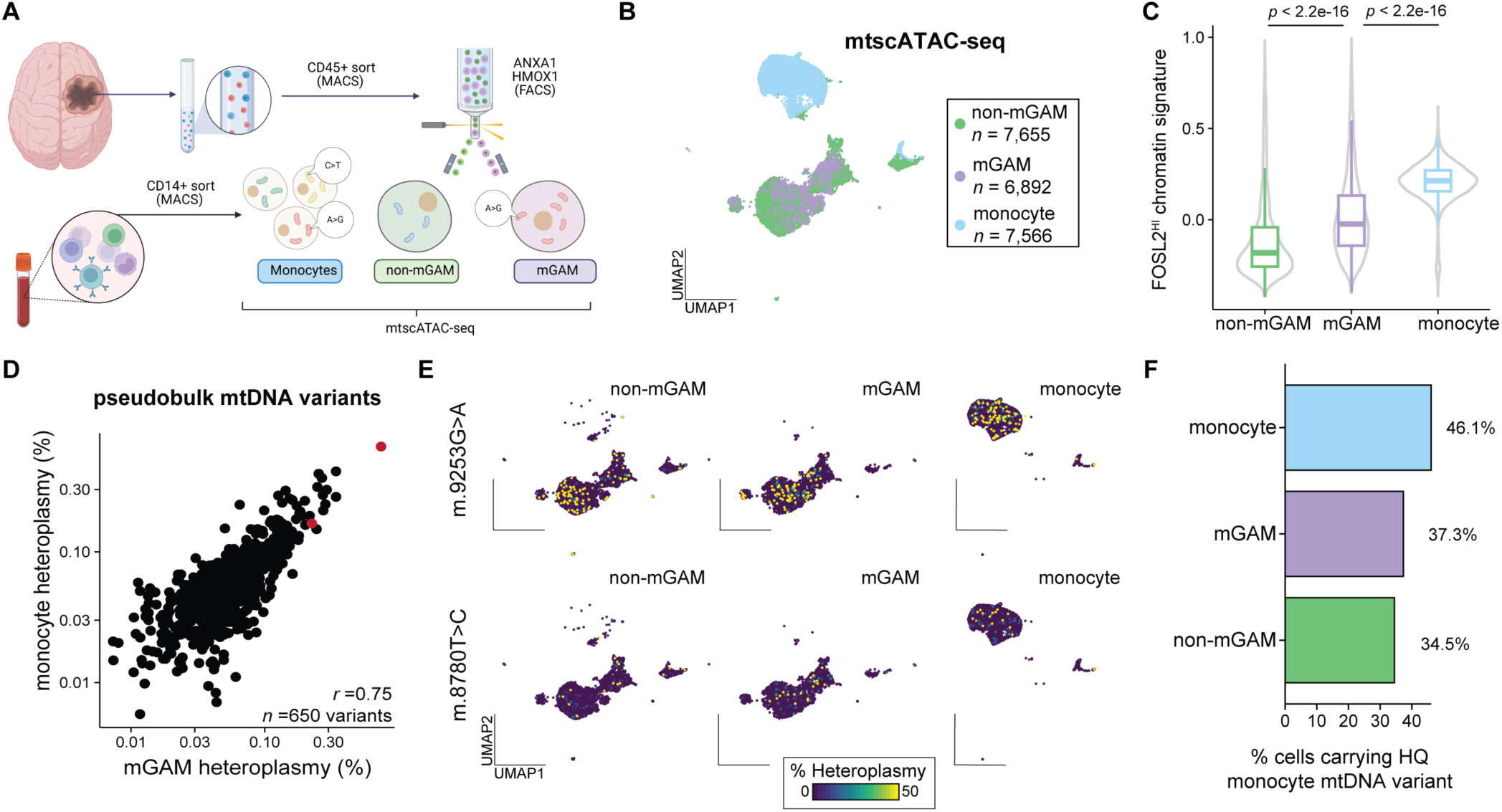
mGAMs share somatic mutations with peripheral monocytes. A) Schematic diagram showing mitochondrial lineage tracing workflow using mtscATACseq. B) UMAP embedding of mtscATAC-seq data in 3 different populations analyzed. C) FOSL2 regulon accessibility profile segregated by the 3 populations analyzed. D) monocyte and mGAM heteroplasmy correlation plot (red dots demonstrate highly heteroplasmic variants shown in next panel. E) Distribution of highly correlated variants in the 3 different populations. F) Percentage of cells with high-confidence monocyte-derived variants within each population.

### mGAM^+^ co-localize to hypoxic niches and dysregulated metabolism in grade dependent manner

We next investigated the distribution of mGAMs within the glioma microenvironment. Cyclic immunofluorescence (CyCIF) staining was performed on frozen tumor tissue from surgical glioma samples which had previously undergone single-cell analysis. mGAM surface markers along with a panel of additional markers such as IBA1, FOSL2, CD44 and SOX2, and hypoxia markers such as HIF1α, were applied to IDH wildtype high-grade gliomas (n=9), IDH mutant gliomas (n=9) and normal brain samples (n=1). Buttressing our findings from the single-cell data, mGAMs, defined as ANXA1^+^HMOX1^+^IBA1^+^ cells, were found throughout high-grade tumors, frequently aggregating in clusters in GBM and high-grade astrocytomas (**Figure S4**).

Analysis of specimens showed increased infiltration of mGAMs in high-grade gliomas (mIDH grade 3 astrocytomas and GBM), compared to mIDH grade 2 astrocytomas (**Figure 6A-B**). mGAM^+^ cells presented high co-expression of FOSL2 (**Figure 6A, C**) and were spatially colocalized in hypoxic niches enriched in HIF1α (**Figure 6A, D**). To compare potential metabolic differences between regions highly enriched in mGAM^-^ and mGAM^+^ cells, single-cell CyCIF data were co-registered into the MALDI-MSI coordinate space. A consensus analysis of metabolic differences between regions highly enriched in mGAM^-^ and mGAM^+^ cells (more than 50% of total cells per pixel) was performed (**Figure 6E-F**), followed by a Monte Carlo simulation to determine statistical significance (**Figure 6E,G**). The metabolic differences between mGAM^-^ and mGAM^+^ regions showed a z-score of 1.25, ranking in the top 11% quantile, and were statistically significant (p < 0.001) in a one-sample t-test. We therefore conclude that mGAM^+^ cells present a significantly dysregulated metabolic profile with respect to mGAM^-^ cells, further supporting the existence of mGAMs in spatially segregated hypoxic metabolic niches as a functionally distinct subpopulation.

**Figure 6:**
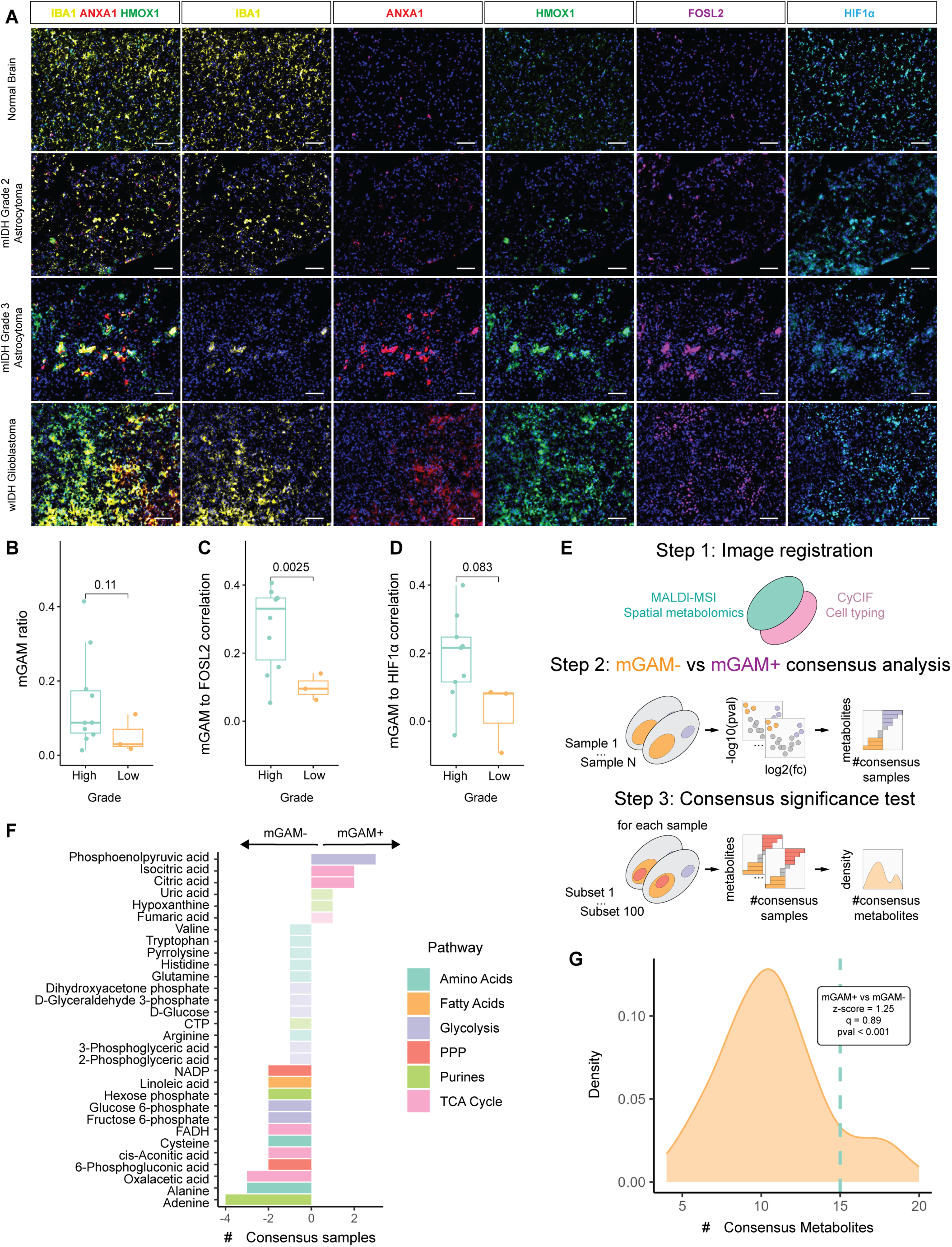
Cyclic Immunofluorescence (CyCIF) of representative regions within normal brain, low-grade glioma, grade 3 astrocytoma and glioblastoma fresh frozen human specimens. A) Highlighted proteins include IBA1, ANXA1, HMOX, FOSL2, and HIF1α. Normal brain and low-grade glioma samples show reduced mGAM^+^ (IBA1^+^, ANXA1^+^, HMOX^+^) infiltration, while astrocytoma and glioblastoma samples exhibit increased mGAM^+^ infiltration. mGAM^+^ cells co-express FOSL2 and colocalize with IBA1-enriched niches. Quantification in 10 high-grade (IV and III) and 3 low-grade (II) samples. B) mGAM^+^ ratio relative to the IBA1+ population and, spatial correlation (Pearson’s r) within the IBA1+ population of mGAM^+^ to C) FOSL2 and, D) HIF1α. E) Schematic of the computational method used to compare metabolism dysregulation between regions highly enriched in mGAM^-^ and mGAM^+^ cells. F) Consensus analysis of metabolism dysregulation between highly enriched mGAM^-^ and mGAM^+^ (more than 50% of total cells per pixel). G) Multiple randomly subsampled regions of the mGAM^-^ region are compared to the remaining mGAM^-^ pixels to estimate expected differences within the mGAM^-^ population in a Monte Carlo setting (orange density distribution). Metabolic differences between mGAM^-^ and mGAM^+^ regions (green dotted line) yield a z-score of 1.25, ranking in the top 11% quantile, and are statistically significant (p < 0.001) in a one-sample t-test.

### mGAMs induce mesenchymal transition in tumor cells and are linked to low grade glioma progression

To evaluate the impact of mGAMs on low grade glioma microenvironment, we injected mGAMs and non-mGAMs from a high-grade tumor into a low-grade glioma organotypic 3D culture (OTC) ^44^ and observed their growth characteristics and pattern (**Figure 7A**). OTCs are 4 mm^3^ morsels of tumor that retain cytoarchitecture and immune cells in ex-vivo culture conditions when grown under Matrigel encapsulation for several weeks. In the mGAM condition, distinct processes from the LGG OTC were seen to extend into the surrounding Matrigel after approximately 7 days in culture compared to the non-mGAM condition (mean area of extended processes = 115772 pixel^2^ (s.d. = 87886 pixel^2^) vs 43346 pixel^2^ (s.d. = 27164 pixel^2^, p<0.0001) mGAM^+^ vs mGAM^-^ conditions). Next we assessed the direct effect of mGAMs on IDH mutant tumor cells by performing co-culture experiments with the low-grade glioma cell line BT-142 ^45^ with mGAMs and non-mGAMs isolated from 4 patients with high grade IDH mutant or wildtype gliomas (MSK149, 153, 154, 157). In all patients we observed consistent increased spheroid formation in the mGAM co-culture condition (**Figure 7B**). CD44 expression has been associated with EMT, increased stemness ^46^ and mesenchymal transcriptional subtype ^47^, therefore we analyzed samples for CD44 and KI67 expression via flow cytometry and detected a consistent increase in the CD44^+^KI67^+^ population in the mGAM co-culture condition over the non-mGAM and control condition (**Figure 7D, G**), which appeared to coincide with increased neurosphere formation in the mGAM co-culture condition. To confirm that mGAM co-culture did indeed induce the expression of a mesenchymal profile in BT142 cells, bulk RNA sequencing was performed on cells from the mGAM co-culture experiment versus BT142 cells alone, which showed a marked upregulation of the mesenchymal-like (MES-like) transcriptional profile in the mGAM co-culture condition, which was completely absent in BT142 cells at baseline (**Figure 7F**). Flow analysis showed that nearly all cells were CD45-by Day 11 (**Figure S8G**), suggesting that this mesenchymal induction process occurred early on. Given the consistent upregulation of CD44 at the cell surface marker level (**Figure 7G**), we next sought to identify potential partner ligands which may be associated with this transition.

**Figure 7:**
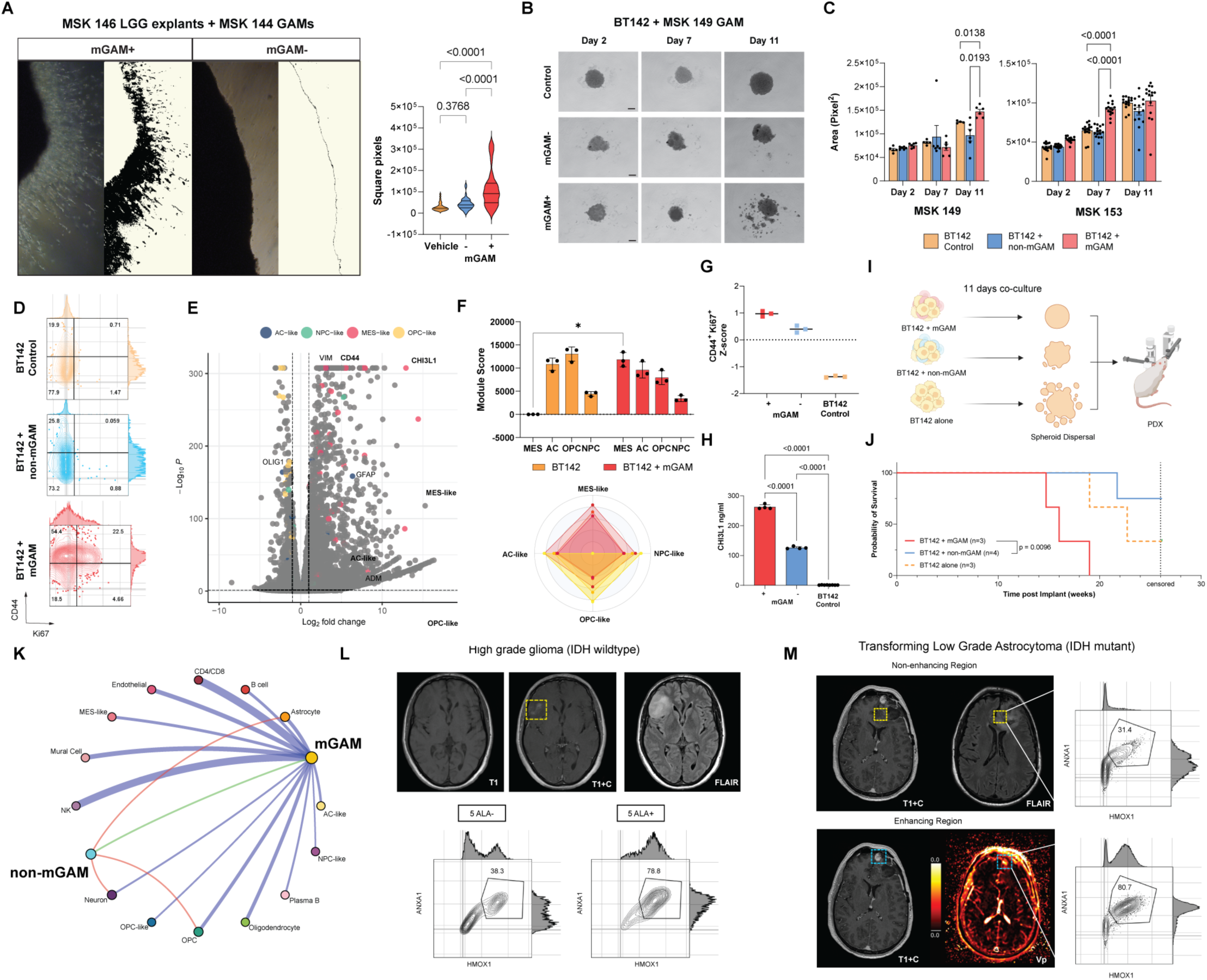
mGAMs are associated with low grade glioma progression. A) Low-grade glioma OTC invasion assay (scalebar = 200µm). Invasive processes are seen extending into the surrounding Matrigel in the mGAM injected condition with quantification of extended processes (right). B) BT-142 co-cultured with mGAMs and non-mGAMs from MSK149 at 11 days demonstrating increased spheroid formation in mGAM condition (scalebar = 200µm). C) Quantification of spheroid number and size based on image analysis over time showing significant difference between mGAM co-culture in terms of overall size and number of spheroids in 2 different patient derived GAMs. D) Representative flow cytometric analysis for KI67 and CD44 showing increased proliferation (KI67) and expression of CD44 in mGAM co-culture compared to non-mGAM condition. E) Volcano plot showing bulk RNA sequencing results comparing BT142 versus BT142 co-cultured with mGAMs, note high expression of CD44 and CHI3L1 in mGAM co-cultured condition. F) Upper: module score of Neftel glioma transcriptional subtype before and after co-culture with mGAMs. Lower: Radar plot of shift in transcriptional profile of BT142 cells after co-culture with mGAMs (p<0.05; n=3 replicates from one patient sample). G) Comparison of proportion of CD44^+^KI67^+^ cells between culture conditions (n=3, each dot represents a different patient sample). H) Measurement of CHI3L1 levels using ELISA showing high levels of CHI3L1 expression in the mGAM co-culture setting (n=4 replicates from one patient sample). I) Schematic diagram showing experimental plan for in-vivo implantation of co-culture conditioned cells. J) Kaplan Meier Curve showing survival of animals after injections of co-cultured cells (cells from one patient sample). K) Cellchat analysis of incoming and outgoing connections in mGAM and non-mGAM cells. (Blue = exclusive to mGAM, Red = exclusive to non-mGAM, Green = ligand-receptor interactions between mGAM and non-mGAMs. L) MRI in patient with IDH wildtype with radiological evidence of high grade transformation. High grade regions identified with 5-ALA and separately analyze with flow cytometry with mGAM markers (lower panel). M) MRI in a patient with a patient with IDH mutant transforming low-grade glioma undergoing surgical resection. Upper panel: T1 weighted contrast enhanced MRI highlighting targeted resection of non-enhancing component followed by flow cytometric analysis for mGAMs. Lower panel: T1 weighted MRI with contrast and perfusion imaging showing active enhancing component (transforming region) within a low-grade glioma. (T1 = T1 weighted MRI, T1+C = contrast enhanced T1 weighted imaging, FLAIR = Fluid Attenuated Inversion Recovery sequence, Vp = Plasma Volume measure on MR perfusion imaging). Targeted resection followed by flow cytometric analysis shows increased proportion of mGAMs.

We noted very high levels of expression of the CD44 ligand Chitinase-like ligand 1 (CHI3L1) in the mGAM co-culture condition from RNA sequencing (**Figure 7E**) and subsequently validated CHI3L1 protein levels through ELISA. High levels of CHI3L1 (>250ng/ml) were found in the supernatant of the mGAM co-culture condition compared to the non-mGAM co-culture (∼100ng/ml). In contrast BT142 cells did not produce any CHI3L1 at baseline (**Figure 7H**). To evaluate the effect of mGAM co-cultured glioma cells on tumor progression in vivo, BT142 cells which had undergone mGAM co-cultures were orthotopically implanted in immunodeficient mice, which demonstrated accelerated growth and reduced survival in the mGAM condition compared to the non-mGAM condition (p = 0.01) (**Figure 7I-J**). Taken together, these data suggest that mGAMs isolated from patient specimens consistently induce mesenchymal, proliferative changes in IDH mutant glioma cells, are associated with the increased protein expression of CD44 and CHI3L1, accelerated tumor growth and reduced survival. We also examined other potential ligand receptor cell-cell interactions between mGAMs and other cell types by running the *Cellchat* algorithm on the GBMap dataset (**Figure 7K**). Compared to non-mGAMs, mGAMs have a much higher number of unique interactions compared to non-mGAMs, pointing towards a potential hub role in the TME. Finally, to better understand the relationship between mGAMs and high grade transformation, we prospectively identified clinical cases where distinct, actively transforming region of tumor could be identified pre-operatively on MR imaging. In this setting, nascent contrast enhancement captured on surveillance MR imaging not only served as a marker of high grade transformation, but also provided spatiotemporal information, with the newly transformed region reflecting more recent changes in the disease course compared to the pre-existing non-enhancing regions. Simultaneous sampling of both enhancing and non-enhancing regions would therefore not only mitigate patient-to-patient variability as a confounder, but also provide an opportunity to measure mGAM abundancy from pathologically distinct regions – these regions were surgically targeted using a combination of neuronavigation and/or fluorescent tumor signal after the administration of 5-aminolaevulinic acid, a surgical adjunct which highlights regions of active tumor ^48^. Each region was analyzed for mGAM markers which showed a higher proportion of mGAMs in 5-ALA fluorescent areas (78.8% versus 38.3% respectively) in an IDH wildtype patient (**Figure 7L**), and importantly higher abundance in areas of a transforming IDH mutant low grade glioma (80.7% versus 31.4%) versus the non-transformed area (31.4%) (**Figure 7M**). mGAM infiltration was also observed in the non-contrast enhancing regions in both cases, albeit at a lower level, thereby strongly implying that the presence of mGAM infiltration is not simply a byproduct of blood brain barrier breakdown but precedes the transformation process.

## Discussion

Malignant gliomas are diffuse primary brain cancers characterized by unrelenting progression ^1,14^. They characteristically harbor a large population of heterogeneous myeloid cells or GAMs and are thought to be critical to tumor pathogenesis. How pathogenic functions are distributed among heterogenous human GAM subpopulations remains unclear and is reflected in the lack of molecular definition and identifying marker sets. Genomic driver mutations such as IDH1/2 mutations and 1p/19q co-deletion correlate with prognosis but are poor markers for disease progression, particularly in low-grade gliomas where the timing and mechanism for transformation is unknown and poorly correlated to mutation burden ^49^. The immune microenvironment is strongly implicated in progression: GAM infiltration correlates with tumor grade ^15^ with many ascribed pro-tumorigenic functions (summarized in ^12^). Single-cell RNA sequencing studies have demonstrated GAM heterogeneity and the spectrum of gene expression within the GAM population; frequent co-expression of classical M1 and M2 genes support the view that context-specific GAM activation states exist and extend beyond the M1/M2 dichotomy, yet there remains no identifiable correlation between in vitro defined macrophage phenotype markers and macrophage cluster signature genes identified in GBM patient tumors ^19–24,33^. We therefore proposed to derive the imprint of the glioma environment on GAM transcription and demonstrate the functional significance of pathological GRNs in human GAMs: We show that cluster-based identification of GAM subsets is inconsistent and reference dependent (**Figure 2A**) and GAM subpopulations can be better resolved through GRN inference. A gene regulatory circuit consists of a transcription factor, *cis*-acting elements and chromatin changes that coordinate the expression of genes in a given cell type under specific conditions ^4^. Using this approach we identify a pathogenic subset within the GAM population which we term mGAMs. Our computational strategy was a multi-step process aimed at improving the signal to noise ratio prior to GRN inference, enabling us to identify the FOSL2 transcription factor network. FOSL2 (also known as fra-2) is a member of the Activator Protein 1 (AP-1) family of transcription factors associated with the acute stress response. FOSL2 has been implicated in the transcriptional network in mesenchymal GBM previously ^50,51^, and has been implicated as a potential master regulator in GAMs ^52^. Dynamic re-wiring of chromatin architecture is known to be mediated by AP-1 transcription factors to bring about co-expression of macrophage regulatory genes ^53^ and FOSL2 and its frequent heterodimer JUN play a role in Epithelial Mesenchymal Transition (EMT) ^54^. The resulting FOSL2 regulon encompasses many of the genes associated with known GAM pro-tumorigenic functions, such as TGFB1, CD163, HMOX1, VEGFA, METRNL matrix metalloproteinases, including mesenchymal genes such as VIM and CD44, and MARCO ^55,56^. Of note, the cytokine Meteorin-L (METRNL) has recently found to be associated with the inhibition of CD8+ T-cell responses ^57^. Thus, the FOSL2 regulon serves as a biologically plausible definition for seemingly heterogenous gene programs expressed by GAMs. By applying network sensitivity analysis, we identify the cell surface markers ANXA1 and HMOX1 as optimal surrogate markers for this underlying network and validated our markers using low-input ChIPseq ^58^ (**Figure 3F**), and ATAC accessibility profiling of mGAMs (**Figure 5F-G**). Annexin A1 is a 37kDa phospholipid binding protein widely expressed on immune, epithelial and endothelial cells, is present both intracellularly and at the cell surface, and mediates anti-inflammatory effects of glucocorticoids ^59^. HMOX1 is a stress induced enzyme involved in the catabolism of heme, with potent anti-inflammatory and cytoprotective effect against hypoxia and its presence on cell surface lipid rafts has been linked to hypoxia ^60^ and its expression is linked to a worse prognosis in glioma ^61,62^. Cell surface mGAM markers allowed immediate isolation of mGAM populations from surgical samples for direct functional analysis. Our data identified marked differences in angiogenic capacity in mGAMs compared to non-mGAM counterparts; secretion of ECM proteins such as vimentin and versican corroborated computational predictions: all pro-tumorigenic functions critical to tumor progression. From an immunological standpoint we show that mGAMs and non-mGAMs are both capable of eliciting polyfunctional T-cell responses to recall antigen ^63^, a feature which has not been previously demonstrated in human glioma samples. The most striking finding however, was the ability for mGAMs to induce CD4^+^FOXP3^+^ T-cells upon co-culture ^64^. T-regulatory cells are classically described as CD4^+^FOXP3^+^CD25^+^ cells, however a subset of T-regs, CD4^+^FOXP3^+^CD25^-^ are considered a precursor to regulatory T-cells ^64^. The mechanism for this induction is unknown but could be related to the pleiotropic effects of TGF beta 1, a constituent target gene within the FOSL2 regulon. These functional findings, along with the spatial co-localization of mGAMs along hypoxic niches, elaborate on recent studies which have shown how the hypoxic TME organizes cell types and layers ^65,66^. The overlay of spatial metabolomic data provides further evidence of dysregulated metabolism in mGAM associated regions, lending credence to the evolving idea of tumor hypoxic niches and metabolites specifying a GAMs pathogenic phenotype. Further spatial transcriptomic analysis at single-cell resolution will help unravel cellular neighborhoods specifically associated with mGAMs.

Transcriptionally mGAMs closely resemble hypoxic monocytes, and the mGAM phenotype can be partially modelled using primary human monocytes (**Figure S6B-E**). Hypoxia is known to activate FOSL2 ^67^, but the binding of FOSL2 to its constituent downstream mGAM target genes appears to require tumor specific input, pointing towards the necessity of both tumor and hypoxic elements for mGAM specification. We attempt to resolve the ontogeny of mGAMs through mitochondrial DNA lineage tracing and show that a significant proportion of mGAMs have a common bone marrow origin to monocytes. The high concordance with the FOSL2 regulon accessibility profile in monocytes strongly suggests that monocytes may be poised for activation towards an mGAM state upon exposure to the tumor hypoxic niche. Given the putative bone marrow derived origin of mGAMs and associated immunosuppressive genes, it may be reasonable to speculate that mGAMs are simply Myeloid Derived Suppressor Cells (MDSC), immature bone marrow cells derived from myeloid progenitors which have been shown to have immunosuppressive effects ^68^. However the lack of consensus on unique human markers for MDSCs has rendered reliable identification of human MDSCs in gliomas challenging, and scRNA-seq studies to date have found considerable transcriptional diversity within MDSC gates ^69^. Flow analysis for standard MDSC markers (defined as CD14^+^CD33^+^HLA-DR^low^ (M-MDSC), and CD16^+^CD33^+^HLA-DR^low^ (P-MDSC)) in our experiments did not identify mGAMs (**Figure S3E**), although we acknowledge that this was a limited sample set (n=3). mGAMs therefore do not fit neatly into existing MDSC nomenclature and instead, we define mGAMs based on activity of the FOSL2 driven gene program and correspondent accessibility profile, irrespective of ontogeny. This molecular definition encompasses and may explain the myriad of GAM subtypes proposed in the literature – CD163, HMOX1, MARCO, CD73, KDM6B, HIF1A, as potential markers of distinct GAM subsets (**Figure S5**). The represents a shift away from markers that define computationally defined clusters to ones that define a functional subgroup through robust GRN inference and functional validation.

To better understand the causal relationships between mGAMs and low grade glioma progression, we directly implanted mGAMs isolated from high-grade tumors into low-grade glioma tumor OTCs and observed invasive outgrowth into surrounding matrix. Co-culture of mGAMs with low grade glioma cells also caused increased glioma cell proliferation and concomitant expression of CD44, an adhesion and cancer stem cell marker ^43^. Bulk RNA sequencing showed a clear shift in mesenchymal subtype upon co-culture with mGAMs, and BT142 cells co-cultured with mGAMs express high levels of CHI3L1, a major structural polymer part of the glycoside hydrolase family 18 and is not only secreted by macrophages but can also polarize macrophages through down regulation of injury associated inflammatory responses ^70^. CHI3L1 is also highly expressed in GBM and their overexpression is associated with a worsened prognosis ^71^, however its downstream targets and functions are unclear. Prior studies have noted a strong association between tumour and myeloid cells in the mesenchymal subtype of GBM ^34^, with oncostatin-M (OSM) proposed as a mediator between these two cell types. OSM is not significantly expressed by mGAMs, and unlike CHI3L1, OSM expression does not correlate with prognosis in either at the bulk TCGA level in GBM or LGG (**Figure S8A-D**). It is therefore tempting to speculate that a CHI3L1-CD44 axis may play a role in mesenchymal transition in gliomas, however this will need to be further verified experimentally. Finally we prospectively identified human samples of tumors undergoing active transformation and show that newly transforming regions are associated with higher proportions of mGAMs, further lending support to the idea that mGAMs are critically involved in malignant glioma progression.

### Limitations of Study

Our experimental work is primarily based on human surgical samples with inherent patient variability and sampling heterogeneity. Although mGAMs do not express typical MDSC markers, it is possible that they may share some overlap with other markers of MDSCs. In addition, a significant minority of mGAMs do not share monocytic mitochondrial mutations, therefore we cannot exclude that resident microglia may also be capable of transition to the mGAM phenotype. Further lineage tracing studies will be required to confirm or refute this possibility.

## Conclusion

The presence of a pathogenic subpopulation within GAMs responsible for pro-tumorigenic function has long been suspected however the molecular definition and cell markers have so far been elusive. Here we demonstrate the functional significance of a pathological GRN driven by FOSL2 in GAMs conserved across human high grade gliomas. mGAMs not only provides a unifying explanation for the myriad of GAM phenotypes seen and their ascribed functions, but also links this phenotype to disease progression, forming a potential mechanistic basis for low-grade glioma transformation into high-grade, where the timing of progression is of clinical importance and the risk to benefit ratio of aggressive early intervention remains ill-defined. Temporal precedence of mGAMs in disease progression points towards their potential role as both a prognostic indicator but also a targetable subset within the wider population of GAMs. Future development of spontaneously transforming LGG *in vivo* models will help us design strategies to intercept these cells early in the disease process and to fundamentally alter their natural history.

## Acknowledgements

We would like to acknowledge Alexandra Giantini-Larsen, Alexandra Miner, Shkurte Ademi Donohue, Nidia Claros and Greta Ghita for their technical assistance and in preparing and staining of specimens, and Dr Enrico Moiso for assistance with image analysis.

Research reported in this publication was supported by the National Institute of Neurological Disorders and Stroke of the National Institutes of Health (R21 NS121812) and the National Cancer Institute (R01 CA3865730-01) and Cancer Center Support Grant (P30 CA008748) and Break Through Cancer. Evan D. Bander received funding from the Leon Levy Neuroscience Research Fellowship, Feil Family Brain and Mind Research Institute.

This research was partially funded through the Swim Across America, Ludwig Institute for Cancer Research, and Parker Institute for Cancer Immunotherapy.

We acknowledge the use of the Flow Cytometry Core Facility, and the Proteomics Core Facility.

We acknowledge the help of Ms Morgan Lallo at the Center of Epigenetics Research at Memorial Sloan Kettering Cancer Center for assistance with experimental design and data analysis.

We acknowledge the help and support of Drs Ronan Chaligne and Meril Takazawa at the Single-cell Analytics Innovation Lab (SAIL) at Memorial Sloan Kettering Cancer Center for assistance with experimental design, data generation and analysis.

Some figures were created in Biorender.com

## Author contributions

Conceptualization (KY, VT)

Data curation (KY, ZAM, KH, KT, UT, NSM, CB, VT)

Formal Analysis (KY, ZAM, KT, KH, JG, GB, RK, NYRA, CL, VT)

Funding acquisition (KY, EDB, VT)

Investigation (KY, ZAM, KT, KH, EDB, IS, JG, GB, UT, VT)

Methodology (KY, ZAM, KH, IS, P-JH, JG, GB, CL)

Project administration (KY, VT)

Resources (TM, NYRA, VT)

Software (ZAM, KT, RK, CL)

Supervision (KY, NYRA, TM, VT)

Validation (KY, ZAM, PJH, RK, CL)

Visualization (KY, ZAM, IS, JG, GB, RK)

Writing – original draft (KY, ZAM, GB, CL)

Writing – review & editing (KY, CL, P-JH, VT)

## Data and Code Availability

All datasets are available at the National Centre for Biotechnology Information Gene Expression Omnibus (NCBI GEO). Single-cell RNA sequencing data can be found at GSE275225, and single-cell ATAC sequencing data is available at GSE275224. ChIP-seq data is available at GSE272216. Bulk RNA sequencing data is available at GSE275484. Mitochondrial scATAC data is available at GSE276283.

Published datasets were obtained from GEO repository (identifiers: GSE182109, and GSE166420) for GAMs, and healthy human microglia ^35^ (SynapseID: syn21438358).

Supplemental data is also available at: https://data.mendeley.com/preview/6fb6888f9m?a=09da1332-a11f-4dd0-8d96-9b3064013b9f

Original code can be found at: www.github.com/tabarlab/Glioma-mGAM

## Methods

### Tumor processing and dissociation

Tumor tissue from glioma operations was collected immediately after resection and transported in MACS storage solution (Miltenyi Biotec, Germany) within 15 min to the laboratory. Resected tissue was morselized into small pieces under sterile conditions using sterile scalpels, triturated using pipettes, then transferred to a 50 mL falcon tube (Corning, USA) for incubation with papain (37°C, 25 min) (Worthington, USA) according to manufacturer instructions. After incubation HBSS without magnesium or calcium was then added to make up to 50 ml and the sample was centrifuged (300×g, 5 min, 4°C) and supernatant aspirated before being passed through a 70um cell strainer (BD Biosciences, USA). Erythrocytes were removed by resuspending and incubating the obtained pellet in 5 ml of 1X RBC Lysis Buffer (eBioscience, USA) for 10 min, followed by centrifugation (300×g, 5 min,4°C). Residual myelin and extracellular debris were eliminated using the Debris Removal Kit (Miltenyi Biotec, Germany). Cell counts were determined using an automated cell counter (Nexcelom Bioscience, USA) after resuspending the pellet in PBS. A portion of the single-cell suspensions were then enriched for CD45+ cells by CD45-MACS enrichment (Miltenyi Biotec, Germany). The final cell suspensions were centrifuged again (350×g, 10 min, RT) and isolated unsorted, CD45+, and CD45-cells were stored in Bambanker (Wako, Japan). Cell suspensions were immediately placed in a storage container (Mr. FrostyTM, ThermoFisher Scientific, USA) and stored in LN2. Informed consent was obtained for all patients and human specimens were collected under approved institutional review board number #09-156.

### Parallel single-cell RNA and ATAC sequencing

Single cells from each sample were split into two populations to undergo single-cell RNA- and ATAC-sequencing, respectively. For scRNA-seq, cells were loaded for GEM generation using the 10x Genomics Chromium platform. 3’ RNA gene expression libraries were prepared using the Chromium Single-cell 3′ Reagent Kit v3 (10x Genomics, USA) according to the manufacturer’s guidelines. For the scATAC experiment, we followed the 10x single-cell ATAC–seq protocol from 10x Genomics Chromium (Single-cell ATAC Reagent Kits User Guide). Single-cell ATAC–seq libraries were generated using the Chromium Chip E Single-cell ATAC kit (10x Genomics, USA) and indexes (Chromium i7 Multiplex Kit N, Set A, 10x Genomics, USA) following the manufacturer’s instructions. Final libraries were quantified using a Qubit fluorimeter (Life Technologies, USA) and the nucleosomal pattern was verified using a BioAnalyzer (Agilent Technologies, USA). scRNA and scATAC libraries were sequenced on a NovaSeq6000 (Illumina, USA).

### Flow Cytometry and Fluorescence Activated Cell Sorting (FACS)

For analysis, cells were thawed at 37°C then centrifuged (300×g; 4°C; 5 min); supernatant was aspirated then resuspended with FACS buffer, cell count was performed using automated cell counter (Nexcelom Bioscience, USA), then incubated with Human Fc block (Miltenyi Biotec, Germany) according to the manufactureŕs instructions. Cells were then stained with antibody panel for 30 minutes, re-centrifuged at 300×g; 4°C for 5 min, then re-suspended in FACS buffer. Analysis was performed on the 5-laser Aurora (Cytek Biosciences, USA). Sorting was performed using FACS ARIA (BD Biosciences, USA). Gating was performed using FlowJo (FlowJo LLC, USA). For fluorescence activated cell sorting (FACS), cells were prepared in the same way and diluted to a concentration of between 5-20 x 10^6^ cells/ml, and were sorted on the FACSAria (BD Biosciences, USA) with the assistance of the Flow Cytometry Core Facility at Memorial Sloan Kettering Cancer Center.

### Single-cell RNA-sequencing pre-processing and analysis

Raw sequencing reads were aligned to the GRCh38 reference genome using *Cell Ranger* (v7.1). Following alignment, cells were integrated and analyzed using R (v4.1) and Seurat (v4.3.0) ^36^. Prior to downstream analysis, cells with < 200 or > 7500 expressed genes, or > 25% mitochondrial transcripts were deemed low-quality and removed. Additionally, genes expressed in < 3 cells were removed. The remaining 52,708 cells’ gene expression matrix was normalized and scaled before applying principal component analysis (PCA) for dimensionality reduction, using the top 2000 most variable genes in the single-cell object. Subsequently, the computed PCs were batch corrected for patient-to-patient variability using the *Harmony* package (v1.2.0).

These batch-corrected PCs were then used as UMAP (non-linear dimensionality reduction method) inputs for visualization and downstream dataset exploration. Graph-based cell clustering was performed using K-nearest neighbor (KNN) graphs with the Louvain algorithm, setting the resolution parameter to 0.3. The top genes expressed in each cluster were obtained through *Seurat*’s *FindMarkers* function and used to identify cell types. For visualization, plots were generated with Seurat, ggplot2, and scCustomize v2.1.2 (Marsh SE (2021). scCustomize: Custom Visualizations & Functions for Streamlined Analyses of Single Cell Sequencing. https://doi.org/10.5281/zenodo.5706430.RRID:SCR_024675.).

### Reference Mapping of cells to harmonized glioblastoma reference map

We mapped our wildtype, mutant and control cells to the core GBmap reference ^32^ via *Azimuth*. Briefly, this anchoring method for reference mapping uses the low dimensional structure of the reference to find “anchor genes” in the query dataset. This in turn allows for the projection and transfer of cell annotations between the reference and the query data. Assignment of labels for individual cells in the query dataset is based on the maximum prediction score of a given cell type or state. For our data we performed unsupervised anchoring based on the first 50 principal components and the SCT normalized matrices of the reference and query data. We finally transferred level 3 and level 4 annotations into our single-cell data object metadata.

### Single-cell ATAC-sequencing pre-processing and analysis

Downstream analysis was done in R (v4.1) using *Seurat* (v4.3), *Signac* (v1.5.0), and *chromVAR* (v.1.8). Prior to downstream analysis, single cells were filtered based on the following criteria: peak region fragments > 200 and < 20,000, > 15% fragments in peaks, < 5% reads in blacklisted regions, < 4 nucleosome signal ratio, and > 2 TSS enrichment score. Latent semantic indexing (LSI) was applied to each sample independently (*RunTFIDF*, *FindTopFeatures*, and *RunSVD* functions). Samples were combined, integrated, and batch corrected using *Harmony* on the LSI reduction. The gene activity matrix was generated using *Signac*’s *GeneActivity* function. Transcription factor activities were calculated using *Signac*’s *chromVAR* implementation (*RunChromVAR* function).

### Gene Enrichment Analysis

Gene enrichment analyses were performed with the web-based tool, Enrichr (https://maayanlab.cloud/Enrichr/), using the Gene Ontology (GO) Biological Process 2023 database. The significant GO terms were selected with the threshold of adjusted p-value < 0.05. Enrichr calculates p-values using the Fisher exact test and corrects for multiple testing using the Benjamini–Hochberg method for multiple hypotheses testing. A combined score for each GO term is calculated using the log of the p-value from the Fisher exact test multiplied by the Z-score of the deviation from the expected rank.

### Gene Marker-Pair Identification

The method for marker pair identification was independently derived, however, it is similar in principle to the method described in. 30 surface marker genes from the FOSL2 gene module enriched in mGAMs were identified through the Human Protein Atlas (proteinatlas.org). The expression of these 30 genes was binarized using 1 read as a cutoff for whether the gene was expressed or not. For each of the 435 possible gene pairs, cells with both genes expressed were classified as mGAMs. We compared the performance of this simple classifier with the starting known states and calculated the true positive (TP), false positive (FP), true negative (TN), and false negative (FN) values. The true positive rate (TPR) and false positive rates (FPR) were then calculated for each gene pair as follows:

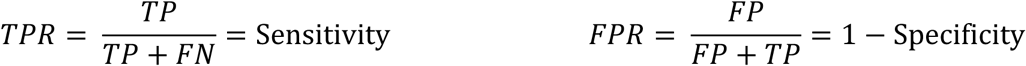

Additionally, to find the optimal gene marker-pair for classifying cells as mGAMs, we calculated the Geometric Mean (G-Mean) metric for each pair, as follows:

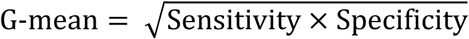

A heatmap of all gene pairs’ G-mean was plotted. The optimal gene pairs were identified for having the highest G-mean among all possible gene pairs.

### Integration of GAM scATAC-seq and scRNA-seq data

After filtering for cells with microglial and GAM signatures we proceeded to integrate data across patients and batches. By using the 2000 most variable RNA features across the 15 sequenced patients we integrated our single-cell ATAC-seq data using CCA-based co-anchoring of the two modalities on the RNA and gene activities ^49^. We confirmed successful integration by visualizing RNA expression and gene activities at canonical genes across patients.

### Integration with external datasets

We obtained publicly available datasets containing wildtype, mutant ^24,33^ (GEO accession numbers: <u>GSE182109</u>, and <u>GSE166420</u>) TAMs, and healthy human microglia ^35^ (<u>SynapseID: syn21438358</u>). As a healthy control we selected cells from individuals with mild cognitive impairment and temporal lobe epilepsy. Similarly, we selected patients with non-recurrent tumors in the diseased samples. We re-processed these samples using the same pipeline we used for our in-lab RNA data. Finally, we integrated cells from our 15 patients with these external data. The final integrated Seurat object was then re-scaled, PCs were re-computed and batch-corrected using *Harmony* (v1.2.0). We generated the UMAP embedding using the batch-corrected PCs from *Harmony*. Consensus clusters were then generated using *FindClusters* with a 0.3 resolution.

### Filtering for canonical wildtype, mutant and control cell populations

We aimed to identify the populations with “purest” wild type, mutant and healthy control signatures, while controlling for potential biases introduced by discretizing cells into clusters. For this, we applied a recently published algorithm for cluster free differential abundance testing ^37^. Briefly, *MiloR* assigns cells to partially overlapping neighborhoods on a k-nearest neighbor graph, which in turn can be tested for differential abundance by using a Negative Binomial GLM framework. We used our external-internal harmonized PCAs as input to the algorithm and then tested for the purest healthy control, wild type and IDH mutant abundance neighborhoods. To identify these cells (n = 4843), we used the cutoffs of logFC > 2 and Spatial FDR<0.001.

### Building gene regulatory networks

We filtered our integrated ATAC and RNA object to cells with the canonical signatures and applied *FigR* ^38^ to infer gene regulatory networks (GRNs) and identify candidate TF regulators. This algorithm does this by first finding the best matching RNA and ATAC cells (*Optmatch*) by optimizing the geodesic distance among all combinations of possible ATAC-RNA cells within the derived kNN graph. We used a search range of 0.1 and the integrated UMAP coordinates next to the CCA corrected PCAs as input. This creates a computationally paired “multiomic” dataset, which can be used as an input to a previously published framework designed to identify significant distal peak-to-gene expression interactions in 100 kb windows around each TSS of a gene. With this we identified a total of 14202 unique chromatin accessibility peaks genome-wide showing a significant association to gene expression (permutation P ≤ 0.05) of 7944 genes. Of these 365 genes showed n ≥ 7 significant peak-gene associations across (DORCs) We finally applied the TF relevance testing across these genes via *FigR*, which simultaneously computes a z-score for TF motif enrichment and a Spearman correlation between the TF RNA expression levels and the DORC accessibility score.

### Defining the FOSL2 regulon signature

Following the building of the FOSL2 regulon via *FigR* as described above, we identified 248 target genes for FOSL2 with a regulation score above 0.5. As our analysis has shown that the FOSL2 regulon is specific to the archetypal WT GAMs, we filtered the target genes by whether they are differentially expressed in these archetypal cells or not, leading to a shortened list of 97 genes. In order to further refine this list, we ranked genes based on their expression level in archetypal WT cells and filtered the list for genes expressed in ≥ 75% of these cells leading to a more targeted gene set of 54 genes representing our FOSL2 regulon gene signature. Gene signature scores were calculated using the *AddModuleScore* function in the *Seurat* package. Cells with a score greater or equal to 0.25 were labeled as FOSL2 regulon high cells.

### TCGA Survival Analysis

Survival analysis was performed using the cSurvival v1.0.1 web portal (https://tau.cmmt.ubc.ca/cSurvival/). Adjusted p-values <0.05 were considered statistically significant. Survival analysis was performed on the TCGA-GBM and TCGA-LGG datasets, using the FOSL2 regulon as our gene set and looking at overall survival (OS).

### Low input ChIP-Seq

Freshly harvested cells were sorted, crosslinked in PBS with 1% formaldehyde for 10 min at room temperature, quenched with 0.125M glycine for 5 min and washed in PBS with protease inhibitors (Millipore Sigma, USA) three times before being snap frozen in cryotubes and sent to MSKCC’s Epigenetics Research Innovation Lab for processing. Low input ChIP-seq was performed using the Low-Cell ChIP-Seq kit (Active Motif, USA) following the manufacturer’s instructions, with the subsequent modifications. After thawing, the fixed cells were resuspended in 130uL of ChIP buffer, transferred into a microTUBE-130 (Covaris, USA) and sonicated using a Covaris E220 ultra-sonicator, with the following parameters: PIP=140, Duty Factor=5, CBP/Burst per sec=200, Time = 900s. After sonication, samples were transferred to siliconized micro-centrifuge tubes and centrifuged at 18,000g for 2 minutes at 4°C. After centrifugation, the supernatants were transferred to new labeled siliconized micro-centrifuge tubes. One percent of each sample was used as input. All samples and inputs were snap frozen and stored at -80°C. After the chromatin being processed following the manufacturer’s recommendations, all samples and inputs were de-crosslinked overnight at 65°C. DNA was finally isolated using the ChIP DNA Clean & Concentrator kit (Zymo Research, USA) following the manufacturer’s instructions. After quantification of the recovered DNA fragments, libraries were prepared using the ThruPLEX®DNA-Seq kit (Takara Bio Inc., Japan) following the manufacturer’s instructions, purified with SPRIselect magnetic beads (Beckman Coulter), quantified using a Qubit Flex fluorometer (ThermoFisher Scientific, USA) and profiled using a TapeStation 2200 (Agilent Technologies, USA). The libraries were sent to the MSKCC Integrated Genomics Operation core facility for sequencing on an Illumina NovaSeq 6000 (aiming for 30-40 million 100bp paired-end reads per library).

### Epigenome Analysis

ChIP sequencing reads were trimmed and filtered for quality and adapter content using version 0.4.5 of *TrimGalore* (https://www.bioinformatics.babraham.ac.uk/projects/trim_galore), with a quality setting of 15 and running version 1.15 of *cutadapt*. Reads were aligned to human assembly hg38 with version 2.3.4.1 of *bowtie2* (http://bowtie-bio.sourceforge.net/bowtie2/index.shtml) and were deduplicated using *MarkDuplicates* in *Picard Tools* (v2.16.10). To ascertain regions of target enrichment, *MACS2* (https://github.com/taoliu/MACS) was used with a p-value setting of 0.001 and scored against matched input as background. The *BEDTools* suite (http://bedtools.readthedocs.io) was used to create normalized read density profiles. A global peak atlas was created by first removing blacklisted regions (http://mitra.stanford.edu/kundaje/akundaje/release/blacklists/hg38-human/hg38.blacklist.bed.gz) then merging all peaks within 500 bp and counting reads with version 1.6.1 of *featureCounts* (http://subread.sourceforge.net). Data were normalized by sequencing depth (to ten million unique alignments). Peak-gene associations were created by assigning all intragenic peaks to that gene, and otherwise using linear genomic distance to transcription start site. Peak intersections were calculated using *bedtools* v2.29.1 and *intersectBed* with 1 bp overlap. Motif signatures were obtained using Homer v4.5 (http://homer.ucsd.edu). Composite plots were created using *deepTools* v3.3.0 by running *computeMatrix* and *plotHeatmap* on normalized bigwigs with average signal sampled in 25 bp windows and flanking region defined by the surrounding 2 kb.

### Culture of GAMs and supernatant collection

FACS-sorted GAMs were plated on 24-well tissue culture-treated plastic plates and cultured in RPMI-1640 media (Gibco, USA) containing 10% FBS (HyClone, Cytiva, USA) or serum-free macrophage medium (Gibco, USA). Culture medium was determined based on downstream application: RPMI-1640 with 10% FBS for phagocytosis assessment, serum-free macrophage medium for ELISA. After 24 hours, supernatant was harvested and stored at -80°C until further analysis.

### Phagocytosis assay

Approximately 24 hours after initial plating, GAM medium was changed to serum-free DMEM/F12 (Gibco, USA) with 1x N2 supplement (Invitrogen, USA). After medium change, pHrodo Red E.coli BioParticles (ThermoFisher Scientific, USA) were added at a dilution of 1:100 to each well. Cells were incubated with bioparticles for 30 minutes at 37°C. Following incubation, cells were washed 1x with DPBS (Corning, USA) to remove excess bioparticles and then dissociated from the plate using Accutase (Stem Cell Technologies, USA). Cells were centrifuged and transferred to flow tubes for flow cytometric quantification of RFP on the 5-laser Aurora (Cytek Biosciences, USA). Gating, calculation of median/mean fluorescence, and generation of flow plots was performed in FlowJo (FlowJo LLC, USA).

### ELISA of cultured GAM supernatant

Vimentin ELISA was performed using the FastScan™ Total Vimentin ELISA kit (Cell Signaling Technology, USA) according to the manufacturer’s instructions. Versican ELISA was performed using Human Versican ELISA Kit (Novus Biological, USA) according to manufacturer’s instruction. CHI3L1 ELISA was performed using Human CHI3L1 ELISA kit (Biotechne, Germany)) according to manufacturer’s instructions. Supernatant samples were diluted 1:50 prior to analysis. Colorimetric measurement was performed using Biotek Synergy HTX microplate reader (Agilent Technologies, USA).

### Sample preparation for t-CyCIF and MALDI-MSI

Fresh frozen human specimens were sectioned at 10μm thickness using a cryostat (Micro hm 550, Thermo Scientific, USA) and thaw-mounted onto optical slides for t-CyCIF. Consecutive sections were mounted onto indium tin oxide (ITO) coated slides (Bruker Daltonics, USA) for MALDI-MSI.

### Cyclic Immunofluorescence

Tissue Cyclic Immunofluorescence (t-CyCIF) was performed on fresh frozen human specimens. Specimens mounted on slides were fixed with 4% PFA for 20 minutes at room temperature, then permeabilized for 5 minutes with 0.1% detergent solution (Thermo Scientific, USA). Primary antibodies diluted in Hoechst 33342 (Thermo Scientific, USA) were incubated overnight at 4oC in the dark. Specimens were photobleached in a solution of 4.5% H2O2, 24mM NaOH in PBS for 1 hour under an LED light source to inactivate fluorophores. T-CyCIF protocol was adapted from .doi.org/10.17504/protocols.io.bjiukkew using the antibodies listed in Table SX. Images were acquired on a CyteFinder II slide scanning fluorescence microscope (RareCyte Inc., USA) with CyteFinder software (v3.11.024). Stitching and registration of tiles and cycles were done in MCMICRO using ASHLAR (v1.10.2) module. Single-cell segmentation was performed using the cell detection feature in QuPath. Single-cell data were exported and thresholds for identifying positive cells for each marker were visually determined for each sample, using prior knowledge of the expected staining patterns. All quantitative statistics and plots were generated in R. Additional details and code can be at found at www.cycif.org.

### Matrix-assisted laser desorption/ionization mass spectrometry imaging

Matrix-assisted laser desorption/ionization mass spectrometry imaging (MALDI-MSI) was performed on consecutive tissue sections. A matrix solution of 1,5-diaminonaphthalene hydrochloride was prepared at a concentration of 4.3 mg/mL in a mixture of 4.5 parts HPLC-grade water, 5 parts ethanol, and 0.5 parts 1 M HCl (v/v/v). A TM-sprayer (HTX imaging, USA) was used to apply a uniform layer of MALDI matrix, with the following four-pass spray method: a flow rate of 0.09 mL/min, spray nozzle velocity of 1200 mm/min, spray nozzle temperature of 75°C, nitrogen gas pressure at 10 psi, a track spacing of 2 mm, and Mass spectrometry imaging experiments were conducted in negative ion mode using a timsTOF fleX mass spectrometer (Bruker Daltonics, USA). The instrument was calibrated with a tune mix solution (Agilent Technologies, USA) via the electrospray source. Instrumental parameters included a pixel step size of 50 µm, covering the m/z range of 52– 1000. Each pixel was acquired with 500 shots at a laser power of 47% (arbitrary scale) and a frequency of 10000 Hz.

Data was visualized and exported to imzml format using SCiLS Lab software (version 2023a core, Bruker Daltonics, USA).

### MALDI-MSI and CyCIF registration

CyCIF single-cell data was used to create reference cell density images for each sample at the same resolution as the MALDI-MSI data. Correspondingly, MALDI-MSI data was used to generate reference TIC images for each sample. An in-house MATLAB script allowed the registration of the single-cell CyCIF data into the MALDI-MSI coordinate space. Using the MATLAB function *cpselect*, fiducial points were manually set on distinctive features common to both images. The function *fitgeotrans* computed a projective similarity transformation, and the centroids of all cells were transformed using the *transformPointsForward* function. An in-house R package then integrated the transformed cell data into the MALDI-MSI object, performed cell typing, and computed per-pixel cell-type-specific counts.

### Consensus metabolism comparison mGAM+ vs mGAM-

Highly enriched regions for mGAM+ and mGAM-were identified by retaining only those pixels where 50% of all cells belonged to each respective group. Differential abundance comparisons between the groups were conducted on a per-sample basis. For each metabolite and sample, fold change and FDR-adjusted p-values (using a t-test) were calculated. The consensus analysis indicated the number of samples agreeing on the dysregulation of each metabolite.

To test the statistical significance of the consensus metabolic differences between mGAM+ and mGAM-regions, a Monte Carlo simulation strategy was employed. For each sample, 100 randomly sampled subsets of the mGAM-group, matching the size of the mGAM+ population, were compared to the remaining mGAM-pixels. The number of metabolites dysregulated in at least two samples (consensus metabolites) was recorded for each iteration. This distribution estimated the inherent metabolic heterogeneity of the mGAM-enriched areas. The statistical significance of the number of consensus metabolites obtained in the mGAM+ vs. mGAM-comparison was assessed by computing the corresponding z-score, quantile, and one-sample t-test.

### Angiogenesis Tube Forming Assay

Human umbilical vein cells (HUVEC) were first cultured in a 6-well tissue culture treated plate (Corning, USA) in EBM-2 Basal Medium (Lonza, USA) supplemented with EGM-2 SingleQuots Supplement Pack (Lonza, USA) containing 2% FBS, 0.04% hydrocortisone, 0.4% hFGF-B, 0.1% VEGF, 0.1% R3-IGF-1, 0.5% ascorbic acid, 0.1% hEGF, 0.1% GA-1000, and 0.1% heparin. Cells are used at low passage (between 2-6). One day prior to assay, HUVEC cells are starved of VEGF and grown for an additional 24 hr. 50µl of reduced growth factor basement membrane (Geltrex, Gibco) is placed in each 96 well. HUVEC cells are washed in DPBS (Corning, USA) and dissociated by adding Accutase (Stem Cell Technologies, USA). Cells are counted and 15,000 cells are plated in each 96 well. Tubes were visualized at multiple timepoints. Peak tube formation was seen at 18 hours. Calcein AM (Thermofisher, USA) was added to the endothelial cells in media at a final concentration of 2 µg/ml and applied at the final timepoint for visualization. Pictures were taken with the upright microscope and images stitched together using ImageJ. Quantification of Tube Network was performed manually, with number of tubes counted noted for each well.

### Glioma and Patient-derived monocyte co-culture and bulk RNA-seq

Patient-derived PBMCs from non-glioma patients are sorted for CD14+ monocytes through magnetic-activated cell sorting (MACS) using CD14 MicroBeads (Miltenyi Biotech, Germany). These monocytes are then co-cultured alongside a previously established patient-derived GBM cell line, MSK-103. Cells are maintained for three days in glioma culture media, consisting of DMEM/F12 basal media (Gibco, USA) supplemented with 1x N2 supplement (Invitrogen, USA), B27 without Vitamin A (Gibco, USA), EGF (20 ng/mL; R&D Systems), FGF2 (20 ng/mL; R&D Systems, USA), 1x Penicillin-Streptomycin (Gibco, USA), and 5% FBS (Gibco, USA). Following the initial three-day co-culture, hypoxia was induced by culturing the cells under 0.5% O2 for 4 hours. Cells from monocultures and co-cultures at normoxic and hypoxic conditions are then collected, stained for CD45 (BioLegend, USA) and sorted through FACS. The CD45+ sorted cells are then lysed for RNA collection and purification using RNeasy Mini kit (Qiagen, USA) as per the manufacturer’s protocol. RNA library was constructed, and RNA sequencing was performed on the Illumina Novaseq 6000. FASTQ were checked for read quality and the clean reads were aligned to the human reference genome (hg38) using the Illumina Dragen v3.10 pipeline. Gene expression count matrices were obtained through *featureCounts* v2.0.3. RNA-seq analysis was performed using *DESeq2* (v.1.40.2). Genes with log2 fold change ≥ 1 and adjusted p-value < 0.05 were considered as DEGs. Enrichment analysis was performed using the *GSEApy* v1.1.2 package. PCA plots, Venn diagrams and gene expression heatmaps were generated using *Matplotlib* v3.9.0.

### **Low-grade** glioma spheroid co-culture assay

BT142, a grade III IDH1-mutant anaplastic astrocytoma cell line ^45^, was purchased from ATCC. BT142 cells and previously sorted GAMs were centrifuged at 300g for 5 mins, washed with PBS and centrifuged again. The supernatants were discarded, and the pellets were resuspended at a concentration of 100 cells/uL using BT142 growth media consisting of DMEM/F12 basal media (Gibco, USA) supplemented with B27 without Vitamin A (Gibco), EGF (20 ng/mL; R&D Systems, USA), FGF2 (20 ng/mL; R&D Systems, USA), heparan sulfate (2 mg/mL; R&D Systems), and PDGF-AA (100 ng/mL; Peprotech, USA). BT142 cells were then mixed with mGAMs or non-mGAMs at a 1:1 ratio. Mixed cells were seeded into wells of an ultra-low attachment 96-well U-bottom plate (PrimeSurface®; S-Bio, Japan) at a concentration of 10,000 cells per well in a final volume of 100uL. Wells seeded with only 5,000 cells of BT142 were also prepared. The plate was then centrifuged at 90g for 2 min. Cells were cultured in a humidified 37°C, 5% CO2 incubator, with partial media changes every 2 days. Cells were observed daily under an inverted bright-field microscope for spheroid formation. At day 11, spheroids were dissociated using the Papain dissociation kit (Worthington Biochemical, USA), stained for CD45 (BioLegend, USA) and CD44 (BioLegend, USA), fixed with Cytofix/Cytoperm solution (BD Biosciences, USA), followed by incubation with Perm/Wash buffer, and stained for intracellular Ki67 (BD Biosciences, USA). Data was acquired on a 5 laser Aurora full spectrum cytometer (UV-V-B-YG-R, Cytek, USA) and analyzed using FlowJo software v. 10.10.0 (FlowJo LLC, USA).

### Organotypic 3D culture Invasion Assay

Organotypic 3D cultures assays (OTC) were processed from fresh tumor samples, on ice, immediately post-resection. The tumor tissue is dissected into smaller pieces (∼6mm in length). Multiple small sphere-like OTCs are then punched out of the tumor pieces using a sterile 2mm biopsy punch (Integra Miltex, USA). Small spherical depressions are made in a sterile parafilm square. We add 10 μL of growth factor reduced Matrigel® (Corning, USA) to each depression before transferring one 4mm^3^ OTC into each well followed quickly by another 20 μL of Matrigel®. The Matrigel-embedded OTCs are allowed to fully polymerize in a 37°C incubator for 30 minutes before being transferred into a 6-well plate at 4-5 OTCs per well with 3 mL of OTC culture media: DMEM/F-12 media (Gibco, USA), N-2 media supplement (Gibco, USA), and 1x Penicillin-Streptomycin (Gibco, USA). Culture media is changed every other day. FACS-sorted GAM populations are injected into each OTC at a concentration of 50,000 cells/OTC in 1 μL under a dissecting microscope with a 1 μL Hamilton Syringe (Model 7001) and an AS point style 25-gauge needle. OTCs were followed daily for changes and imaged on an inverted bright-field microscope, for a week following treatment.

### Isolation of primary cells

Human blood samples were collected in heparinized CPT tubes from patients undergoing tumor resection. PBMCs were obtained through centrifugation of CPT tubes for 15 minutes at 2500 rpm. PBMCs were collected from the interphase, washed three times with PBS for 5 minutes at 1500 rpm and either cryopreserved for future use or subjected to T cell isolation. Pan T cells were isolated through magnetic bead selection according to manufacturer’s instructions using EasySep™ Human T Cell Isolation Kit (Stemcell Technologies, USA). FACS-sorted glioblastoma-associated macrophages (GAMs) were resuspended in RPMI containing, 10% FBS, NEAA, 2 mM L-glutamine, penicillin/streptomycin, 1 mM sodium pyruvate, and 50 uM β-mercaptoethanol (full culture media) at a concentration of 0.3-0.5 x 10^6^/mL and cultured overnight under normoxic (20% O2) or hypoxic (3% O2) conditions.

### Peptide stimulation of T cells

Isolated autologous T cells were added to conditioned GAMs at a 1:1 ratio and co-cultured with DMSO, 1 ug/mL CEF peptide pool (JPT Peptide Technologies, Germany), or 1 ug/mL SARS-CoV2 spike protein peptide pool (Miltenyi Biotec, Germany) overnight under normoxic or hypoxic conditions. After 24 hours, cells were transferred to normoxia and cultured for an additional 9 days in the presence of 10 IU/mL hIL-2 and 10 ng/mL hIL-15. Media containing cytokines was replenished every two to three days. On day 10, T cells were re-stimulated by addition of peptides and subjected to intracellular cytokine staining.

### Intracellular cytokine staining and T cell phenotyping

On day 10 after culture setup, T cells were restimulated as described above. Monensin was added to the culture after 1 hour of stimulation at a final concentration of 1-2 uM. After an additional 4-6 hours, cells were washed and stained with Fixable Viability Dye – Zombie NIR (BioLegend, USA) for 15 minutes on ice in PBS. Cells were washed twice and incubated with human Fc blocking reagent (Miltenyi Biotech, Germany) for 15 minutes in PBS/2% FBS on ice. The antibody cocktail for surface staining containing the following antibodies was added in 2x concentration to cells and Fc block: anti-human CD3 – BUV395 (BD Biosciences, USA), anti-human CD4 – BV650 (BD Biosciences, USA), anti-human CD8 – BUV615 (BD Biosciences, USA), anti-human CD45RA – BUV737 (BD Biosciences, USA), and anti-human CD62L – BUV563 (BD Biosciences, USA). Cells were incubated for 30 minutes on ice and washed twice in PBS/2% FBS for 3 minutes at 2000 rpm. Cells were prepared for intracellular staining using the FoxP3/Transcription Factor Staining Buffer Set (ThermoFisher Scientific, USA). Intracellular staining was performed in permeabilization buffer for 45 minutes on ice with the following antibodies: anti-human IFNγ – PE/Dazzle594 (BioLegend, USA), anti-human TNFα, and anti-human FoxP3 – eFluor450 (Invitrogen, USA). Cells were washed in permeabilization buffer and resuspended in PBS. Data was acquired on a 5 laser Aurora full spectrum cytometer (UV-V-B-YG-R, Cytek, USA) and analyzed using FlowJo software v. 10.10.0 (FlowJo LLC., USA).

### Mitochondrial lineage tracing sample preparation

Immune cells were first isolated from dissociated glioma samples via CD45^+^ MACS followed by FACS purification for mGAM and non-mGAM populations. Peripheral blood from the same patient was collected and PBMCs isolated using gradient separation (Ficoll-Paque PLUS, Cytiva, USA) followed by CD14 MACS sorting to obtain monocytes. Purified viable cells were used as input to the mtscATAC-seq protocol, and libraries were prepared using the Chromium Next GEM single-cell ATAC reagent kit v2.0 following the manufacturer’s instructions with the modifications described in the mtscATAC-seq protocol. Library size and quality were analyzed before fragmentation and after final purification. Libraries were sequenced on an Illumina NovaSeq X (Illumina, USA).

### Mitochondrial lineage tracing analyses

Mitochondrial genotyping was performed with *mgatk* ^42^ in ‘tenx‘ mode using the barcodes identified as cells by *CellRanger*-ATAC. Cells were further quality-controlled by requiring a minimum of 10x coverage for brain-derived myeloid cells and 15x coverage for monocytes from peripheral blood. Additional chromatin quality control, including a minimum 25% reads in peaks and at least 1,000 nuclear fragments were further applied, resulting in the final cell numbers shown in **Figure 6f**. Variants were called per-library using default mgatk cutoffs ^42^, and a total of 650 variants were identified using the union of the variant calls for all three populations. Population pseudobulks were estimated from the rowMeans of the heteroplasmy matrix. To estimate the proportion of cells derived from peripheral blood, we determined an extended set of 1,017 high-quality monocyte-derived heteroplasmic variants (at least 1 confident cell detected; strand correlation > 0.65; mean heteroplasmy < 0.5) as conducted in a previous expanded analysis ^42^. The percent of cells (**Figure 6j**) were the fraction of cells having at least one of these variants at a 10% heteroplasmy threshold.

### Image Analysis

Images captured by light microscopy at x20 were loaded into Fiji. For neurosphere quantification and for invasion quantification images were first converted into 8-bit black and white images, followed by automated image thresholding with manual adjustment to ensure correct thresholding. The function Analyze Particles was then applied with parameters set to count particles above 50 square pixels. For OTC invasion assay, thresholding was performed to capture the invasive edges and measured in pixels. Statistical analysis was performed using Prism software v.10.2.3 (GraphPad Software, USA).

## Supplemental information

List of Antibodies and Reagents Supplementary Figures (1-8)

FigR results table and list of genes for FOSL2 regulon

**Supplementary Figure 1:**
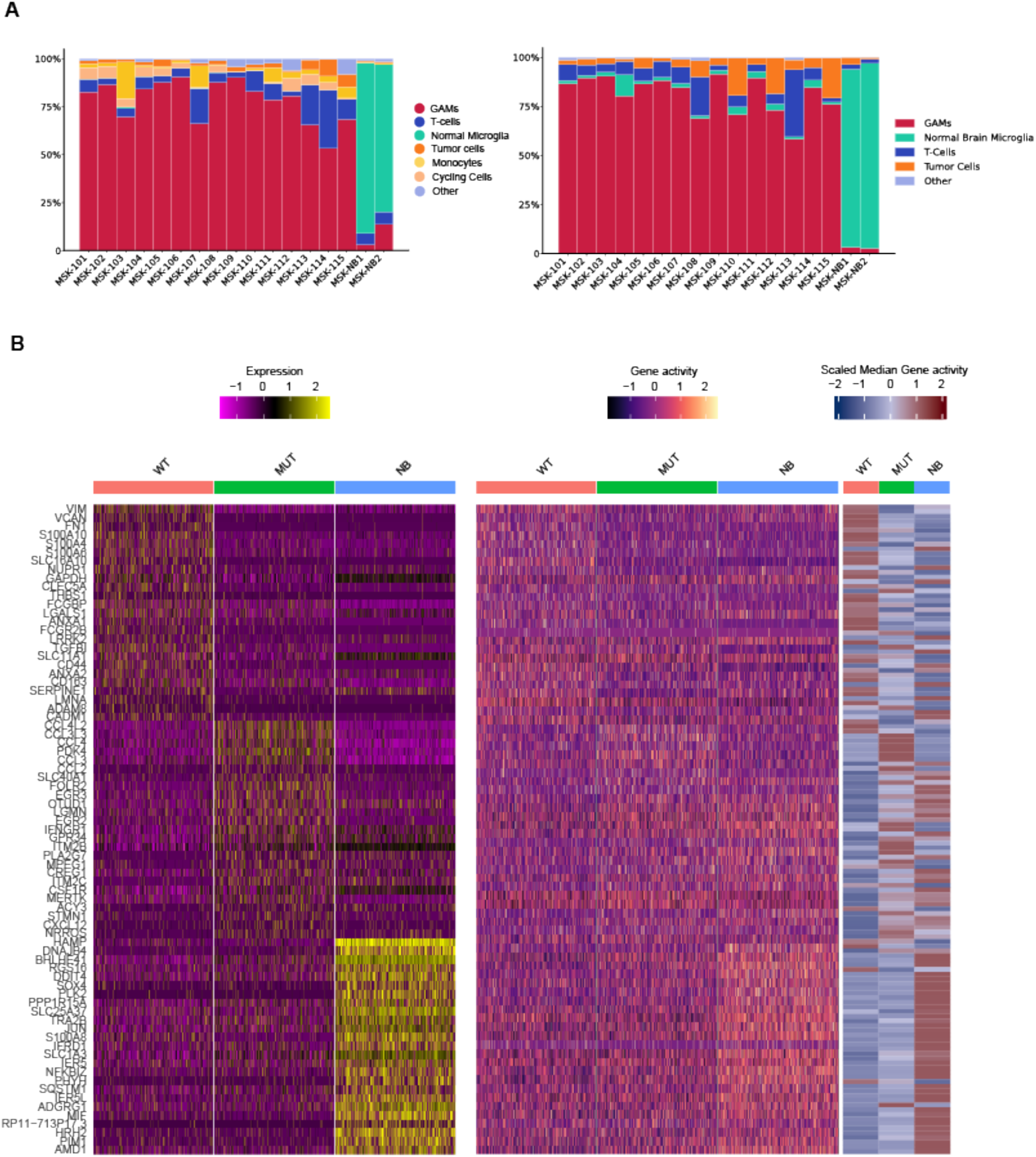
Cell proportions and gene clustering. A) Cell type proportions in parallel scRNA and scATAC data. B) Gene expression clustering for scRNA and Gene activity clustering for scATAC data.

**Supplementary Figure 2:**
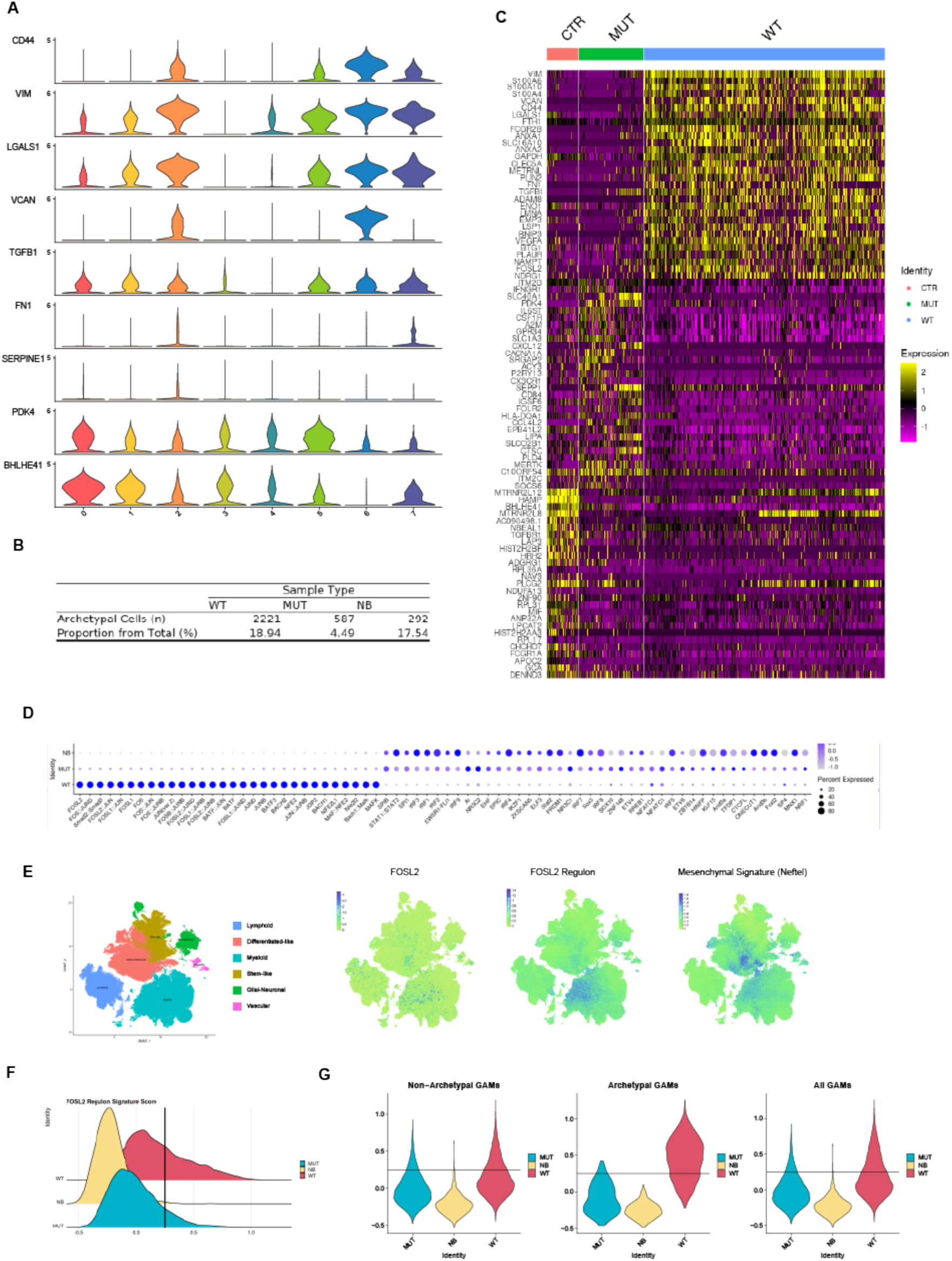
Validation of FOSL2 regulon in public datasets. A) Representative violin plots of differentially expressed genes between wtIDH and mIDH GAMs showing distribution of expression across different clusters. No single cluster demonstrates enrichment for either wtIDH or mIDH associated genes. B) % archetypal cells by condition, based on scRNA data. C) Gene expression re-clustering based on archetypal cells only showing clear separation between conditions. D) ChromVAR analysis on scATAC data to identify accessible motifs in archetypal cells between conditions. Since samples from all conditions were processed in the same manner, any differences in accessibility profile in AP-1 related genes are unlikely to be related to dissociation artefact. E) FOSL2 expression and FOSL2 regulon expression overlaid onto GBMap data ^39^. FOSL2 expression is found in tumor and GAMs, however FOSL2 regulon expression is highly expressed in the myeloid/GAM population. F) Distribution of FOSL2 regulon signature score across IDH wildtype, mutant and control samples. G) Expression of FOSL2 module in non-archetypal cells, archetypal cells and all GAMs.

**Supplementary Figure 3:**
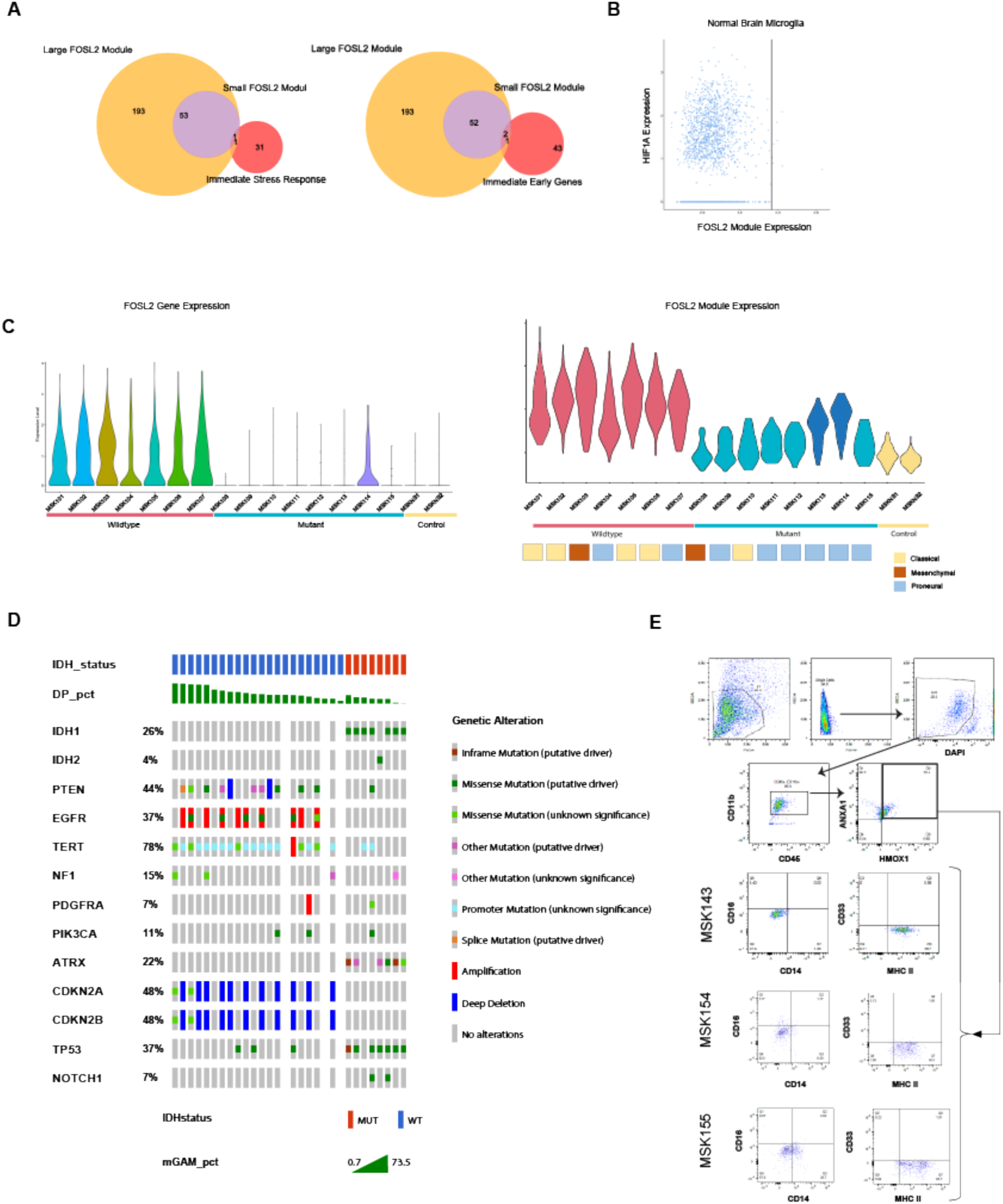
Sample validation. A) Venn diagram showing overlap between Integrated Stress Response genes ^72,73^, and Immediate Early genes ^74^, and the small and large FOSL2 regulon respectively. B) Scatterplot stratifying all microglial cells from Normal Brain samples by HIF1A (y-axis) as a measure of hypoxia, versus FOSL2 regulon expression (x-axis). The cut-off threshold where we call cells as FOSL2 regulon high in the rest of the dataset is indicated. Collectively, this shows that hypoxic microglial cells do not express the FOSL2 regulon, suggesting this is not a purely hypoxia driven phenomenon. C) Left: Violin plots showing FOSL2 expression by patient samples and condition. Right: Violin plots showing FOSL2 regulon expression by patient samples, condition, grade and mutational subtype. D) Oncoprint of samples analysed via flow cytometry using mGAM markers. All wtIDH are high-grade by definition and hence have a higher proportion of mGAMs, however in the mIDH samples, there are no specific mutations associated with low mGAM proportions, with the majority of patients harboring canonical IDH1 and TP53 mutations and ATRX loss and one patient with IDH2 mutation. E) Flow validation experiment re-analyzing flow sorted cells for CD11b and CD45 expression, ANXA1 and HMOX1 expression and expression of MDSC markers. Note the low expression of both CD33 and HLA-DR in mGAM+ population.

**Supplementary Figure 4:**
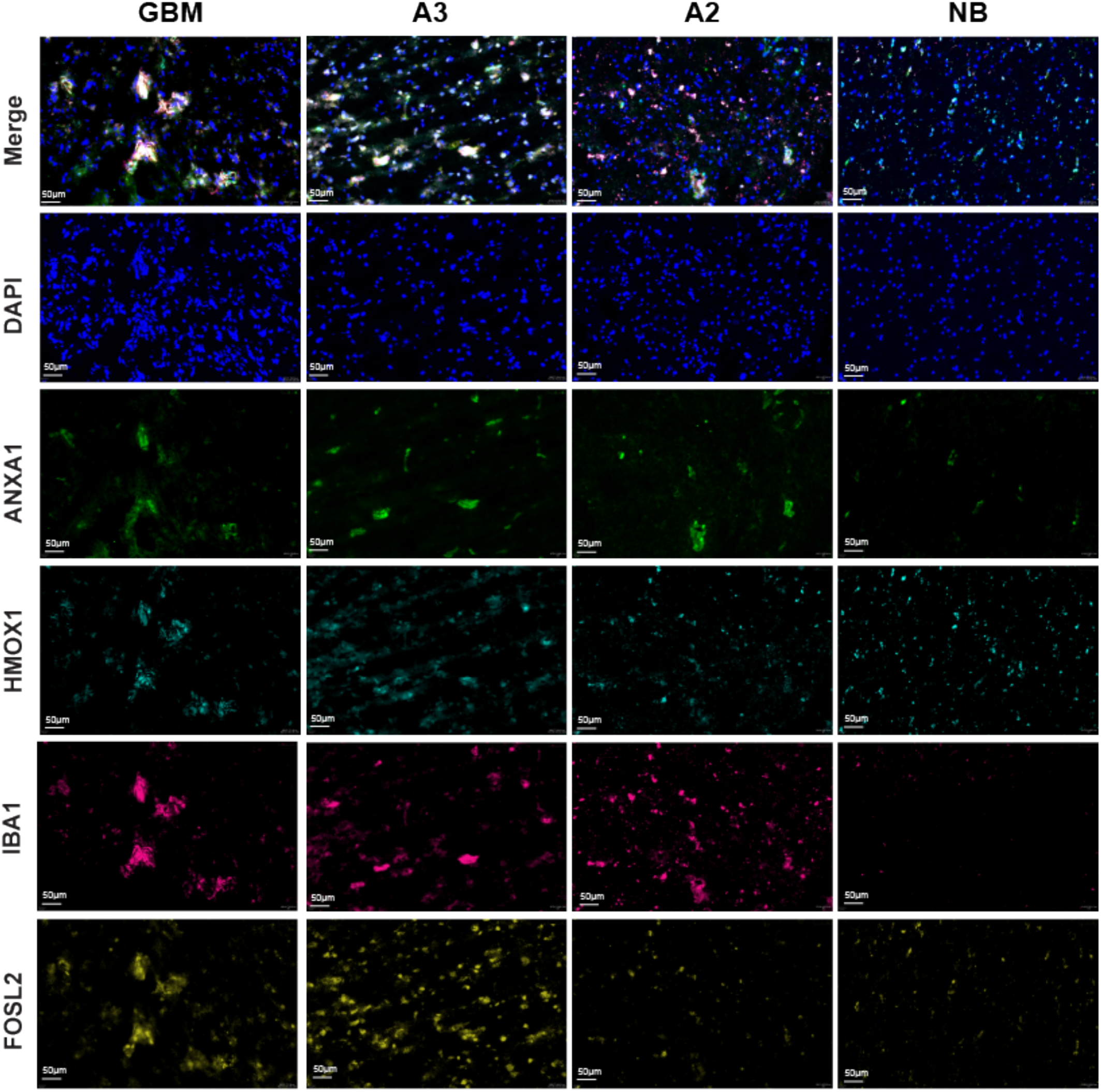
Cyclic IF analysis of different regions in Glioblastoma. (GBM), Astrocytoma grade 3 (A3), Astrocytoma grade 2 (A2) and Normal Brain samples showing individual staining for FOSL2, IBA1, HMOX1, ANXA1, DAPI and merged images. These supplementary images show clustering of co-localized FOSL2, ANXA1, HMOX1 and IBA1 cells in clusters particularly in high grade tumors (GBM and A3).

**Supplementary Figure 5:**
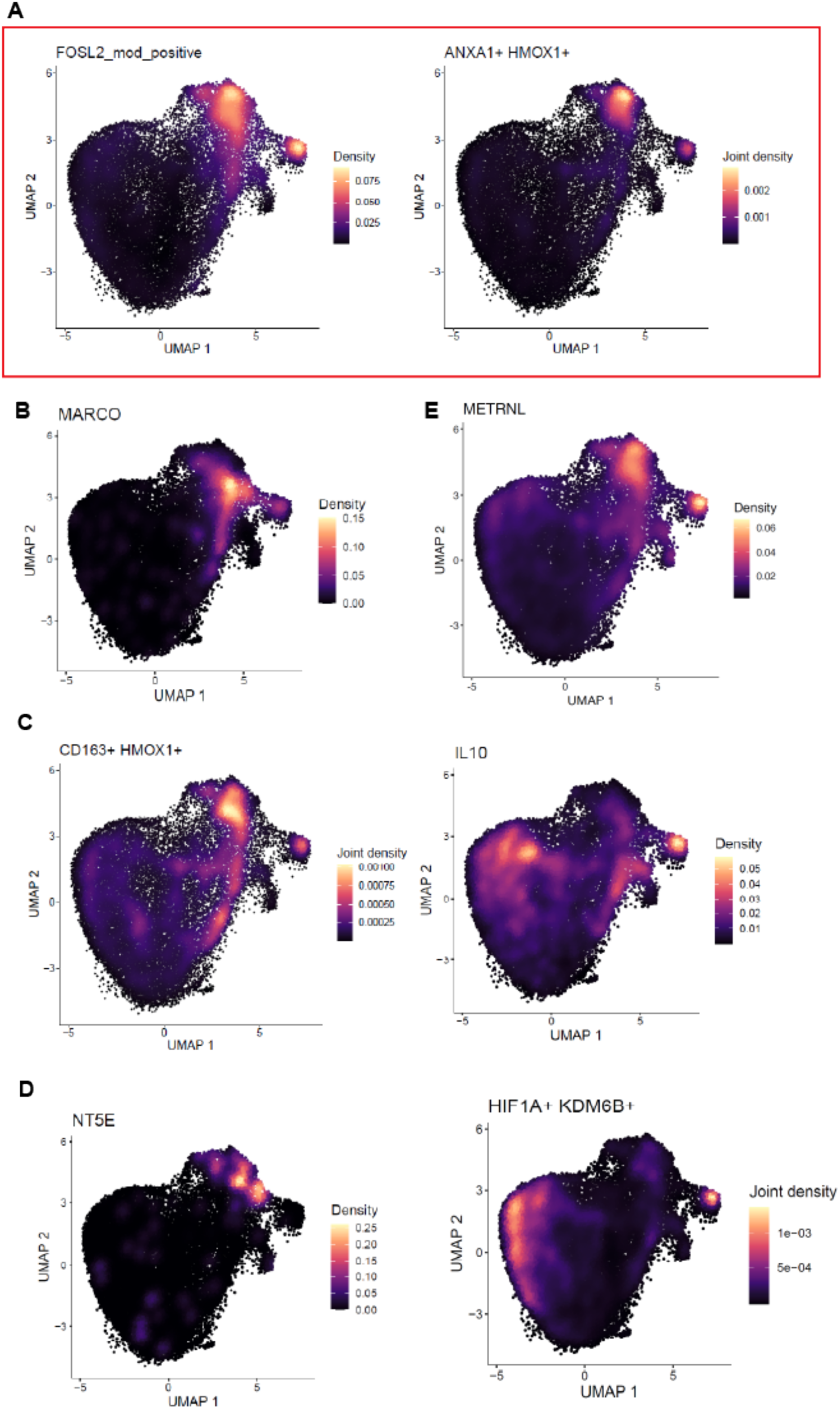
Comparison of proposed GAM subpopulations in the literature versus mGAMs overlaid on consensus dataset described in main text. A) FOSL2 regulon expression and ANXA1/HMOX1 co-expression. B) MARCO expression ^57,58^. C) CD163, HMOX1 co-expression and IL10 expression ^77^. D) CD73 (NT5E) expression ^78^ and KDM6B/HIF1A co-expression ^79^.

**Supplementary Figure 6:**
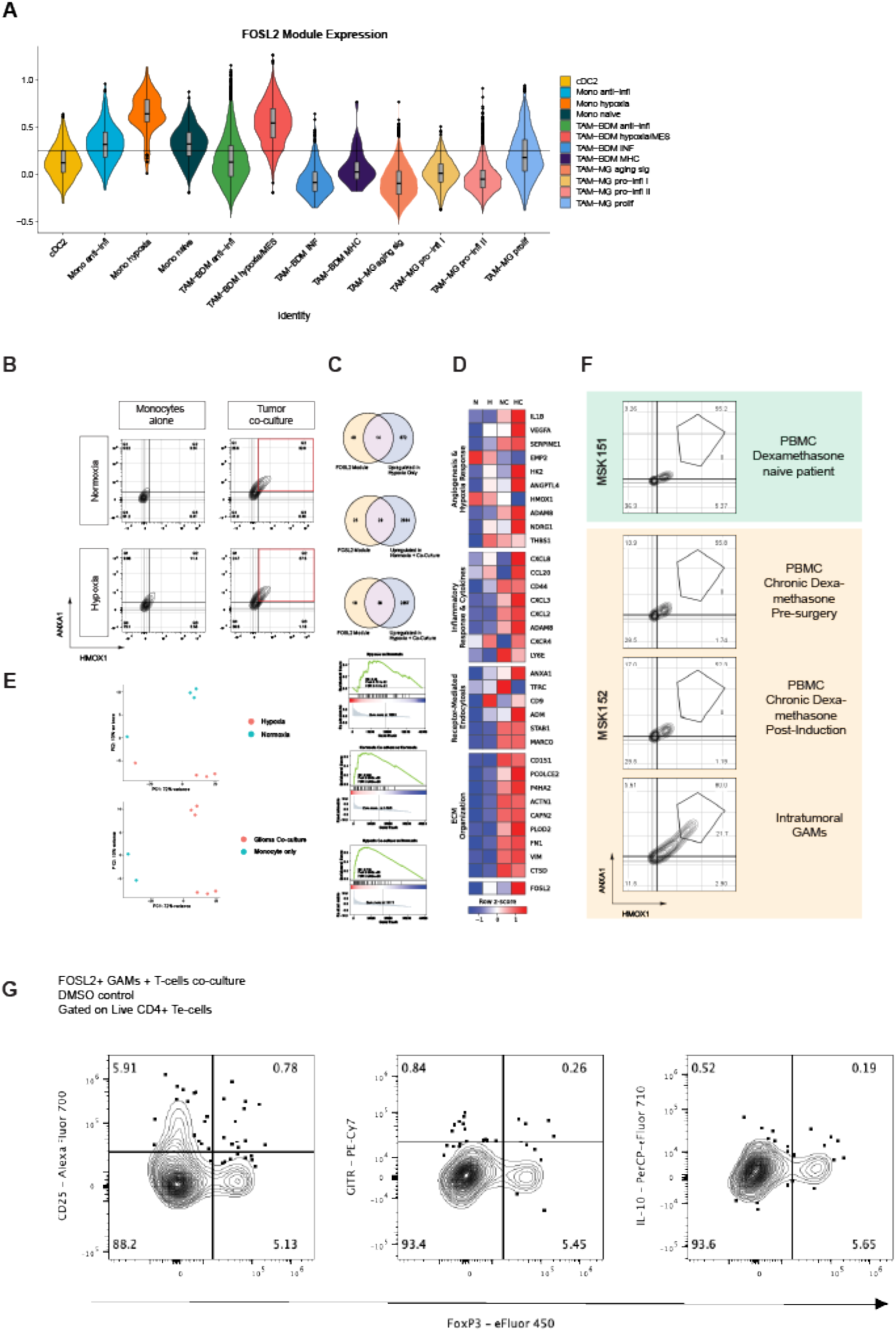
Analysis of primary monocytes and T-cells. A) Violin plot showing the FOSL2 module expression in all human immune cell types defined in GBMap ^32^. B) Flow cytometry plots of ANXA1 and HMOX1in primary monocytes with or without tumor co-culture under hypoxic (0.5%) and normoxic conditions (n=3). C) upper: Overlap between FOSL2 small regulon and monocyte conditions (H = hypoxia, HC = hypoxia co-culture, N = Normoxia, NC = normoxia co-culture) lower: Gene Set Enrichment Analysis of highly differentially expressed genes in bulk RNAseq analysis of conditioned monocytes. D) Heatmap showing expression of FOSL2 regulon associated genes divided by functional subclasses. E) PCA of bulk RNA sequencing samples of different monocyte co-culture conditions. Upper: Hypoxia versus normoxia, lower: Glioma co-culture versus monocyte only. F) Analysis of PBMCs from patients undergoing glioma surgery in dexamethasone naïve and chronic dexamethasone patients. G) T-cell immunophenotyping of T-cells co-cultured with mGAMs showing increased FOXP3 signal.

**Supplementary Figure 7:**
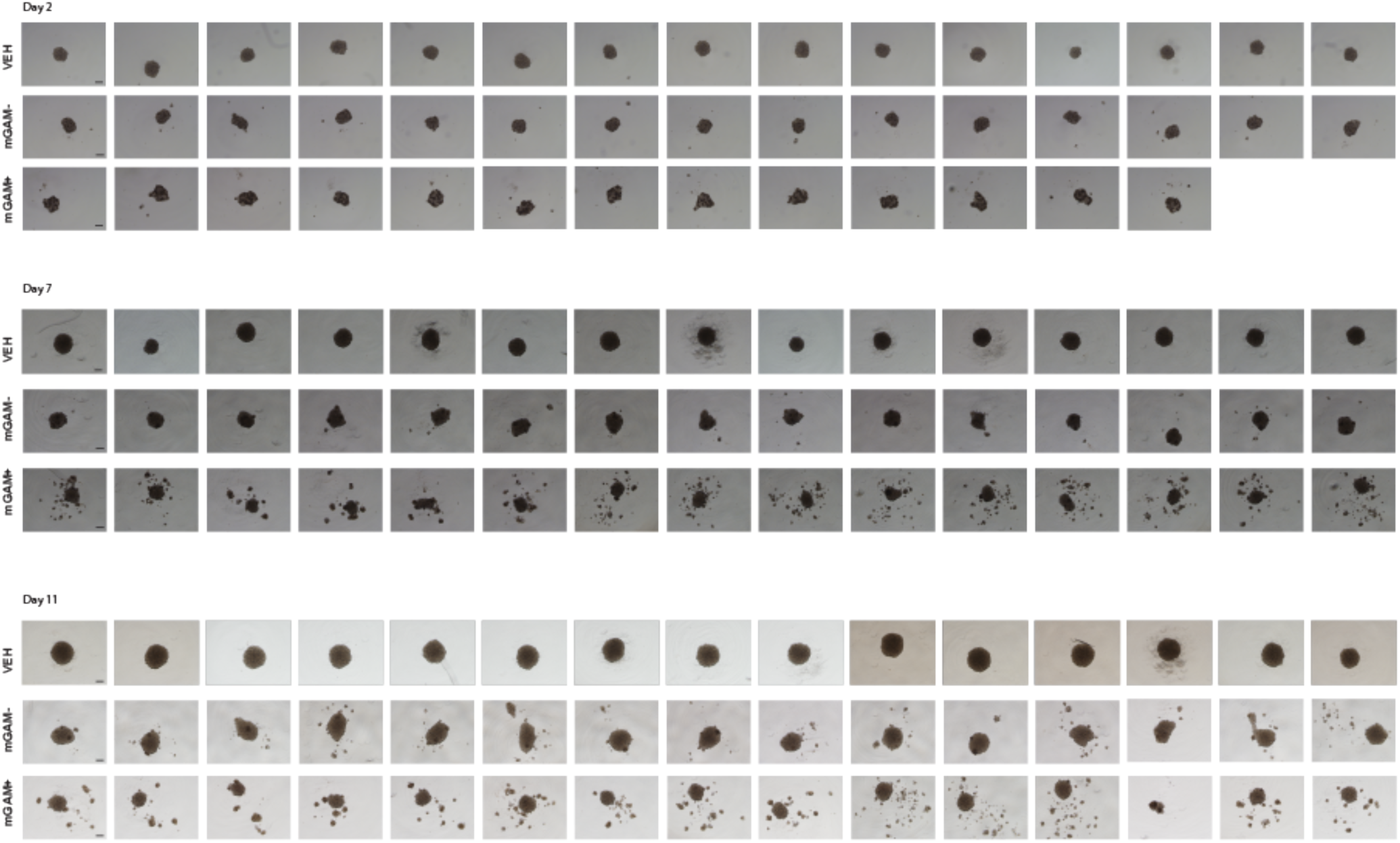
Co-culture of mGAMs from MSK153 and BT142 cells showing consistency of effect across replicates over time. (n=48 per condition, 13-15 wells randomly selected for imaging at each timepoint). An increased number of spheroid particles are seen in the mGAM+ condition, starting from Day 7, quantified in Figure 4d (scalebar = 200µm).

**Supplementary Figure 8:**
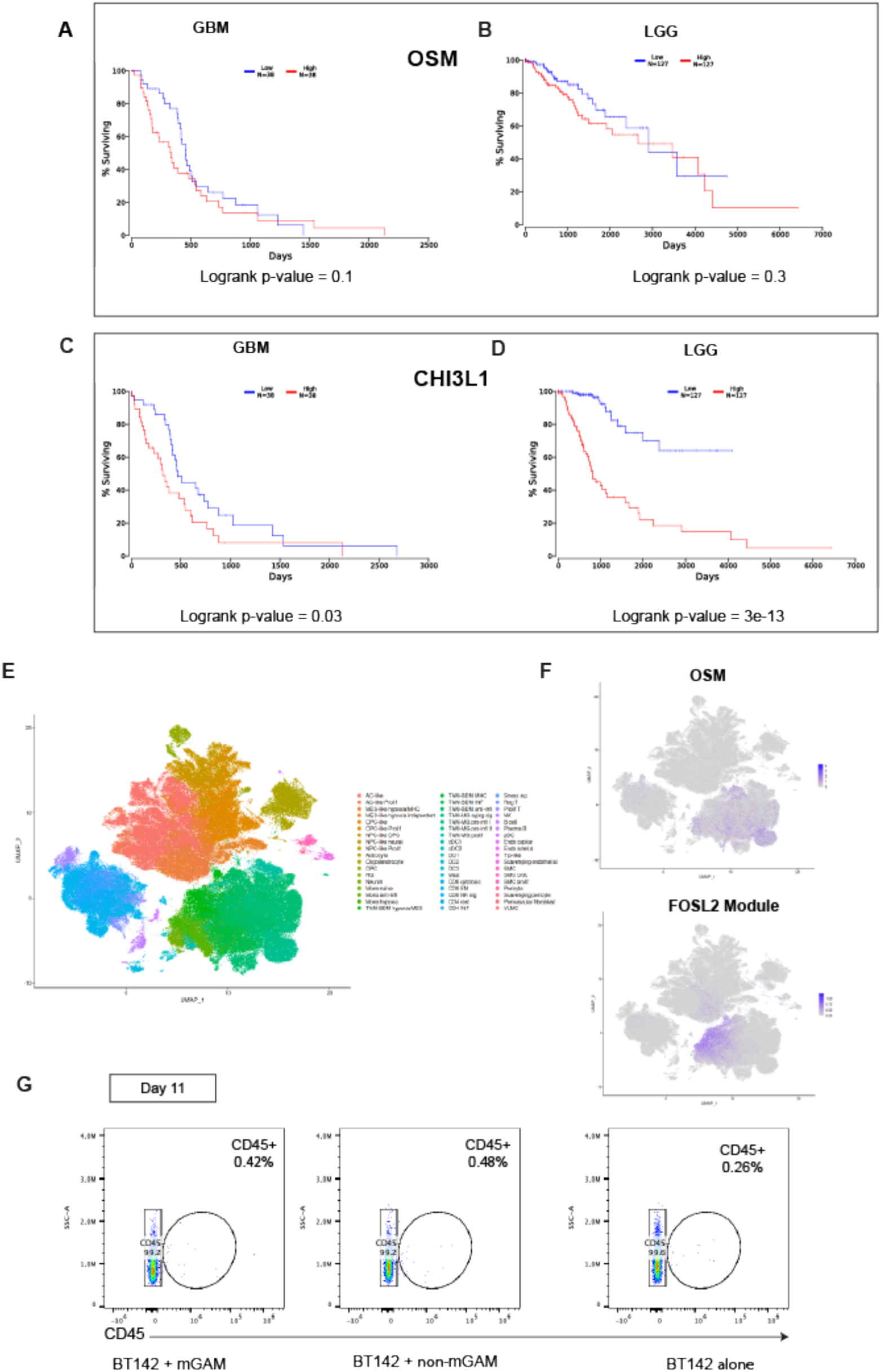
A-D) TCGA survival data in GBM and LGG stratified by expression of Oncostatin-M (OSM) and Chitinase-3-like protein 1(CHI3L1). E-F) GBMap^32^ cell type annotations and overlay of expression of OSM expressing cells versus FOSL2 module. G) Flow cytometry plots of BT142 and mGAM/non-mGAM/control co-cultures at Day 11 showing minimal residual CD45+ immune cells within the co-culture by Day 11.

## References

1. Weller, M. et al. Glioma. Nat. Rev. Dis. Primer 1, 15017 (2015).

2. Jaeckle, K. A. et al. Transformation of low grade glioma and correlation with outcome: an NCCTG database analysis. J. Neurooncol. 104, 253–259 (2011).

3. Verhaak, R. G. et al. Integrated genomic analysis identifies clinically relevant subtypes of glioblastoma characterized by abnormalities in PDGFRA, IDH1, EGFR, and NF1. Cancer Cell vol. 17 98–110 (2010).

4. Brennan, C. W. et al. The somatic genomic landscape of glioblastoma. Cell vol. 155 462–77 (2013).

5. Louis, D. N. et al. The 2021 WHO Classification of Tumors of the Central Nervous System: a summary. Neuro-Oncol. 23, 1231–1251 (2021).

6. Cancer Genome Atlas Research, N. Comprehensive genomic characterization defines human glioblastoma genes and core pathways. Nature vol. 455 1061–8 (2008).

7. Bai, H. et al. Integrated genomic characterization of IDH1-mutant glioma malignant progression. Nat. Genet. 48, 59–66 (2016).

8. Aoki, K. et al. Prognostic relevance of genetic alterations in diffuse lower-grade gliomas. Neuro-Oncol. 20, 66–77 (2018).

9. Alghamri, M. S. et al. Tumor mutational burden predicts survival in patients with low-grade gliomas expressing mutated IDH1. Neuro-Oncol. Adv. 2, vdaa042 (2020).

10. Barthel, F. P. et al. Longitudinal molecular trajectories of diffuse glioma in adults. Nature 576, 112–120 (2019).

11. Suzuki, H. et al. Mutational landscape and clonal architecture in grade II and III gliomas. Nat. Genet. 47, 458–468 (2015).

12. Hambardzumyan, D., Gutmann, D. H. & Kettenmann, H. The role of microglia and macrophages in glioma maintenance and progression. Nat. Neurosci. 19, 20–27 (2016).

13. Ceccarelli, M. et al. Molecular Profiling Reveals Biologically Discrete Subsets and Pathways of Progression in Diffuse Glioma. Cell 164, 550–563 (2016).

14. Varn, F. S. et al. Glioma progression is shaped by genetic evolution and microenvironment interactions. Cell 185, 2184–2199.e16 (2022).

15. Komohara, Y., Ohnishi, K., Kuratsu, J. & Takeya, M. Possible involvement of the M2 anti-inflammatory macrophage phenotype in growth of human gliomas. J. Pathol. 216, 15–24 (2008).

16. Zhai, H., Heppner, F. L. & Tsirka, S. E. Microglia/macrophages promote glioma progression. Glia 59, 472– 485 (2011).

17. Pyonteck, S. M. et al. CSF-1R inhibition alters macrophage polarization and blocks glioma progression. Nat Med vol. 19 1264–72 (2013).

18. Butowski, N. et al. Orally administered colony stimulating factor 1 receptor inhibitor PLX3397 in recurrent glioblastoma: an Ivy Foundation Early Phase Clinical Trials Consortium phase II study. Neuro Oncol vol. 18 557–64 (2016).

19. Muller, A., Brandenburg, S., Turkowski, K., Muller, S. & Vajkoczy, P. Resident microglia, and not peripheral macrophages, are the main source of brain tumor mononuclear cells. Int J Cancer vol. 137 278– 88 (2015).

20. Venteicher, A. S. et al. Decoupling genetics, lineages, and microenvironment in IDH-mutant gliomas by single-cell RNA-seq. Science 355, eaai8478 (2017).

21. Friebel, E. et al. Single-Cell Mapping of Human Brain Cancer Reveals Tumor-Specific Instruction of Tissue-Invading Leukocytes. Cell (2020) doi:10.1016/j.cell.2020.04.055.

22. Klemm, F. et al. Interrogation of the Microenvironmental Landscape in Brain Tumors Reveals Disease-Specific Alterations of Immune Cells. Cell (2020) doi:10.1016/j.cell.2020.05.007.

23. Pombo Antunes, A. R., et al. Single-cell profiling of myeloid cells in glioblastoma across species and disease stage reveals macrophage competition and specialization. Nat. Neurosci. 24, 595–610 (2021).

24. Abdelfattah, N. et al. Single-cell analysis of human glioma and immune cells identifies S100A4 as an immunotherapy target. Nat. Commun. 13, 767 (2022).

25. Martinez-Lage, M. et al. Immune landscapes associated with different glioblastoma molecular subtypes. Acta Neuropathol. Commun. 7, 203 (2019).

26. Glass, C. K. & Natoli, G. Molecular control of activation and priming in macrophages. Nat. Immunol. 17, 26–33 (2016).

27. Lavin, Y. et al. Tissue-resident macrophage enhancer landscapes are shaped by the local microenvironment. Cell 159, 1312–1326 (2014).

28. Gosselin, D. et al. Environment drives selection and function of enhancers controlling tissue-specific macrophage identities. Cell 159, 1327–1340 (2014).

29. Cheng, D. T. et al. Memorial Sloan Kettering-Integrated Mutation Profiling of Actionable Cancer Targets (MSK-IMPACT): A Hybridization Capture-Based Next-Generation Sequencing Clinical Assay for Solid Tumor Molecular Oncology. J. Mol. Diagn. JMD 17, 251–264 (2015).

30. Schep, A. N., Wu, B., Buenrostro, J. D. & Greenleaf, W. J. chromVAR: Inferring transcription factor-associated accessibility from single-cell epigenomic data. Nat. Methods 14, 975–978 (2017).

31. Kiselev, V. Y., Andrews, T. S. & Hemberg, M. Challenges in unsupervised clustering of single-cell RNA-seq data. Nat. Rev. Genet. 20, 273–282 (2019).

32. Ruiz-Moreno, C. et al. Harmonized single-cell landscape, intercellular crosstalk and tumor architecture of glioblastoma. 2022.08.27.505439 Preprint at 10.1101/2022.08.27.505439 (2022).

33. Friedrich, M. et al. Tryptophan metabolism drives dynamic immunosuppressive myeloid states in IDH-mutant gliomas. *Nat*. Cancer 2, 723–740 (2021).

34. Hara, T. et al. Interactions between cancer cells and immune cells drive transitions to mesenchymal-like states in glioblastoma. Cancer Cell 39, 779–792.e11 (2021).

35. Olah, M. et al. Single cell RNA sequencing of human microglia uncovers a subset associated with Alzheimer’s disease. Nat. Commun. 11, 6129 (2020).

36. Hao, Y. et al. Integrated analysis of multimodal single-cell data. Cell 184, 3573–3587.e29 (2021).

37. Dann, E., Henderson, N. C., Teichmann, S. A., Morgan, M. D. & Marioni, J. C. Differential abundance testing on single-cell data using k-nearest neighbor graphs. Nat. Biotechnol. 40, 245–253 (2022).

38. Kartha, V. K. et al. Functional inference of gene regulation using single-cell multi-omics. Cell Genomics 2, 100166 (2022).

39. Sorensen, M. D., Dahlrot, R. H., Boldt, H. B., Hansen, S. & Kristensen, B. W. Tumour-associated microglia/macrophages predict poor prognosis in high-grade gliomas and correlate with an aggressive tumour subtype. Neuropathol Appl Neurobiol (2017) doi:10.1111/nan.12428.

40. Wang, Q. et al. Tumor Evolution of Glioma-Intrinsic Gene Expression Subtypes Associates with Immunological Changes in the Microenvironment. Cancer Cell 33, 152 (2018).

41. Xiao, Y. et al. CD44-Mediated Poor Prognosis in Glioma Is Associated With M2-Polarization of Tumor-Associated Macrophages and Immunosuppression. Front. Surg. 8, 775194 (2021).

42. Lareau, C. A. et al. Mitochondrial single-cell ATAC-seq for high-throughput multi-omic detection of mitochondrial genotypes and chromatin accessibility. Nat. Protoc. 18, 1416–1440 (2023).

43. Ludwig, L. S. et al. Lineage Tracing in Humans Enabled by Mitochondrial Mutations and Single-Cell Genomics. Cell 176, 1325–1339.e22 (2019).

44. Shimizu, F., Hovinga, K. E., Metzner, M., Soulet, D. & Tabar, V. Organotypic explant culture of glioblastoma multiforme and subsequent single-cell suspension. Curr. Protoc. Stem Cell Biol. Chapter 3, Unit3.5 (2011).

45. Luchman, H. A. et al. An in vivo patient-derived model of endogenous IDH1-mutant glioma. Neuro-Oncol. 14, 184–191 (2012).

46. Anido, J. et al. TGF-beta Receptor Inhibitors Target the CD44(high)/Id1(high) Glioma-Initiating Cell Population in Human Glioblastoma. Cancer Cell vol. 18 655–68 (2010).

47. Neftel, C. et al. An Integrative Model of Cellular States, Plasticity, and Genetics for Glioblastoma. Cell 178, 835–849.e21 (2019).

48. Stummer, W. et al. Extent of resection and survival in glioblastoma multiforme: identification of and adjustment for bias. Neurosurgery vol. 62 564–76; discussion 564-76 (2008).

49. Johnson, B. E. et al. Mutational Analysis Reveals the Origin and Therapy-Driven Evolution of Recurrent Glioma. Science 343, 189–193 (2014).

50. Carro, M. S. et al. The transcriptional network for mesenchymal transformation of brain tumours. Nature 463, 318–325 (2010).

51. Wu, L. et al. Natural Coevolution of Tumor and Immunoenvironment in Glioblastoma. Cancer Discov. 12, 2820–2837 (2022).

52. Chen, Y. et al. Tumor-associated monocytes promote mesenchymal transformation through EGFR signaling in glioma. Cell Rep. Med. 4, 101177 (2023).

53. Phanstiel, D. H. et al. Static and Dynamic DNA Loops form AP-1-Bound Activation Hubs during Macrophage Development. Mol. Cell 67, 1037–1048.e6 (2017).

54. Yin, J. et al. HGF/MET Regulated Epithelial-Mesenchymal Transitions And Metastasis By FOSL2 In Non-Small Cell Lung Cancer. OncoTargets Ther. 12, 9227–9237 (2019).

55. Sa, J. K. et al. Transcriptional regulatory networks of tumor-associated macrophages that drive malignancy in mesenchymal glioblastoma. Genome Biol. 21, 1–17 (2020).

56. Chen, A. X. et al. Single-cell characterization of macrophages in glioblastoma reveals MARCO as a mesenchymal pro-tumor marker. Genome Med. 13, 88 (2021).

57. Jackson, C. M. et al. The cytokine Meteorin-like inhibits anti-tumor CD8+ T cell responses by disrupting mitochondrial function. Immunity 57, 1864–1877.e9 (2024).

58. Brind’Amour, J. et al. An ultra-low-input native ChIP-seq protocol for genome-wide profiling of rare cell populations. Nat. Commun. 6, 6033 (2015).

59. Yang, Y. H., Aeberli, D., Dacumos, A., Xue, J. R. & Morand, E. F. Annexin-1 Regulates Macrophage IL-6 and TNF via Glucocorticoid-Induced Leucine Zipper1. J. Immunol. 183, 1435–1445 (2009).

60. Dunn, L. L. et al. New Insights into Intracellular Locations and Functions of Heme Oxygenase-1. Antioxid. Redox Signal. 20, 1723–1742 (2014).

61. Sferrazzo, G. et al. Heme Oxygenase-1 in Central Nervous System Malignancies. J. Clin. Med. 9, 1562 (2020).

62. Nishie, A. et al. Macrophage infiltration and heme oxygenase-1 expression correlate with angiogenesis in human gliomas. Clin. Cancer Res. Off. J. Am. Assoc. Cancer Res. 5, 1107–1113 (1999).

63. Ticha, O., Klemm, D., Moos, L. & Bekeredjian-Ding, I. A cell-based in vitro assay for testing of immunological integrity of Tetanus toxoid vaccine antigen. Npj Vaccines 6, 1–11 (2021).

64. Zelenay, S. et al. Foxp3+ CD25– CD4 T cells constitute a reservoir of committed regulatory cells that regain CD25 expression upon homeostatic expansion. Proc. Natl. Acad. Sci. U. S. A. 102, 4091–4096 (2005).

65. Ravi, V. M. et al. Spatially resolved multi-omics deciphers bidirectional tumor-host interdependence in glioblastoma. Cancer Cell 40, 639–655.e13 (2022).

66. Greenwald, A. C. et al. Integrative spatial analysis reveals a multi-layered organization of glioblastoma. Cell 0, (2024).

67. Zhang, Y., Wang, S., Hu, H. & Li, X. A systematic study of HIF1A cofactors in hypoxic cancer cells. Sci. Rep. 12, 18962 (2022).

68. Gabrilovich, D. I. & Nagaraj, S. Myeloid-derived suppressor cells as regulators of the immune system. Nat Rev Immunol vol. 9 162–74 (2009).

69. Hegde, S., Leader, A. M. & Merad, M. MDSC: Markers, development, states, and unaddressed complexity. Immunity 54, 875–884 (2021).

70. Zhao, T., Su, Z., Li, Y., Zhang, X. & You, Q. Chitinase-3 like-protein-1 function and its role in diseases. Signal Transduct. Target. Ther. 5, 1–20 (2020).

71. Li, F. et al. CHI3L1 predicted in malignant entities is associated with glioblastoma immune microenvironment. Clin. Immunol. 245, 109158 (2022).

72. Verginadis, I. I. et al. A stromal Integrated Stress Response activates perivascular cancer-associated fibroblasts to drive angiogenesis and tumour progression. Nat. Cell Biol. 24, 940–953 (2022).

73. Pakos-Zebrucka, K. et al. The integrated stress response. EMBO Rep. 17, 1374–1395 (2016).

74. Tullai, J. W. et al. Immediate-Early and Delayed Primary Response Genes are Distinct in Function and Genomic Architecture. J. Biol. Chem. 282, 23981–23995 (2007).

