## Supplementary Figures for "A pathogenic subpopulation of human glioma associated macrophages linked to glioma progression"

**Supplementary Figure 1**

**A**

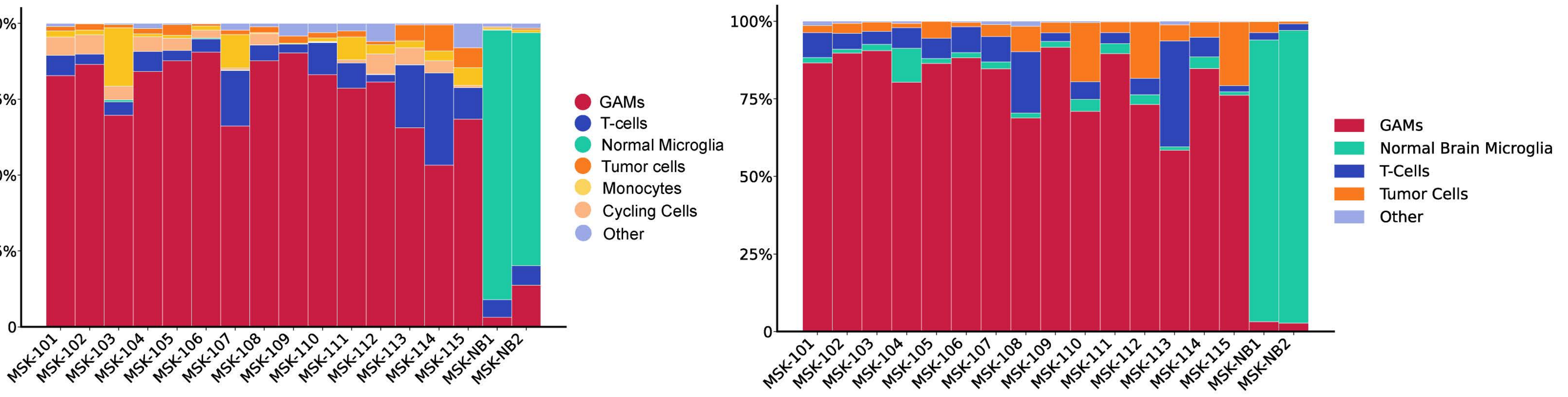


**B**

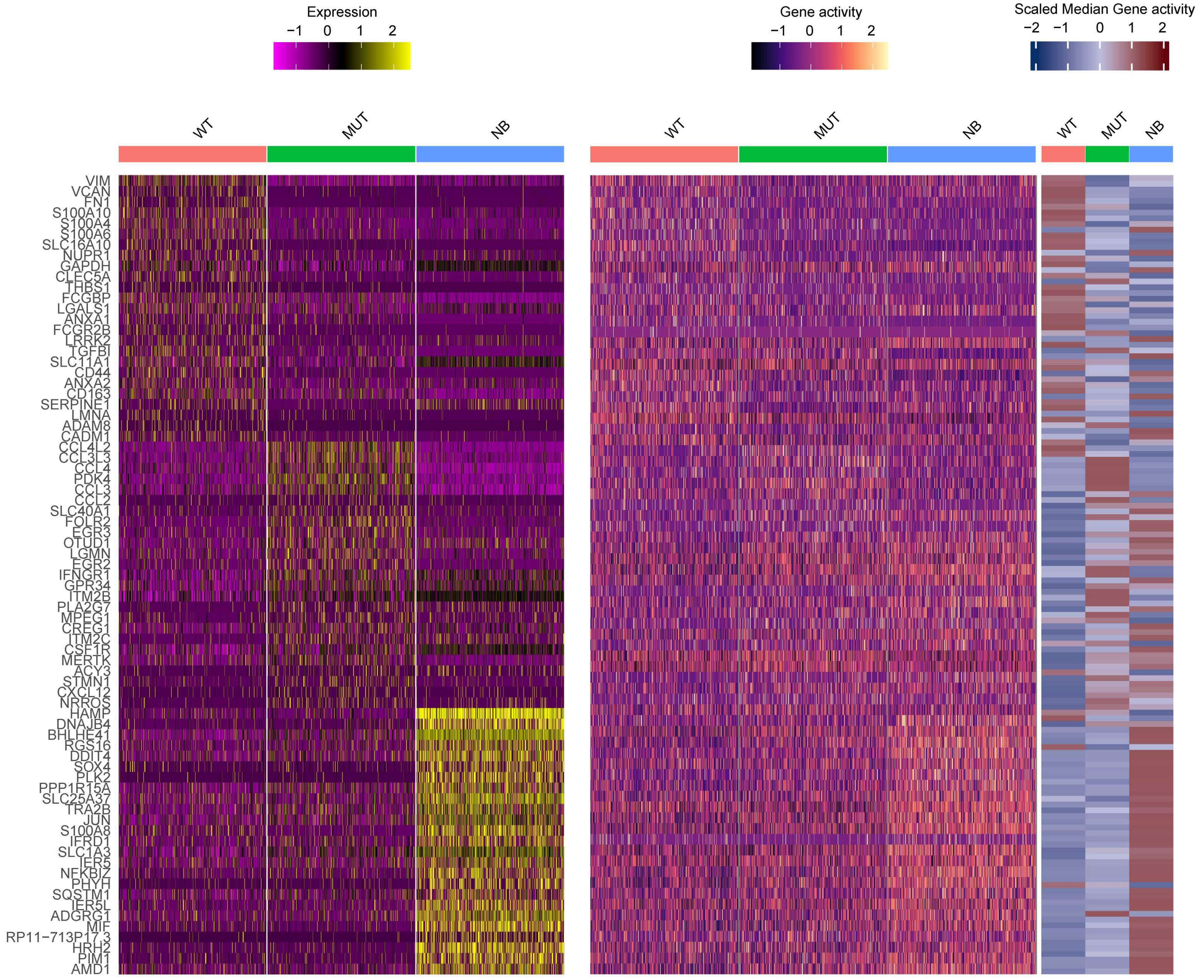

### Supplementary Figure 2

**A**

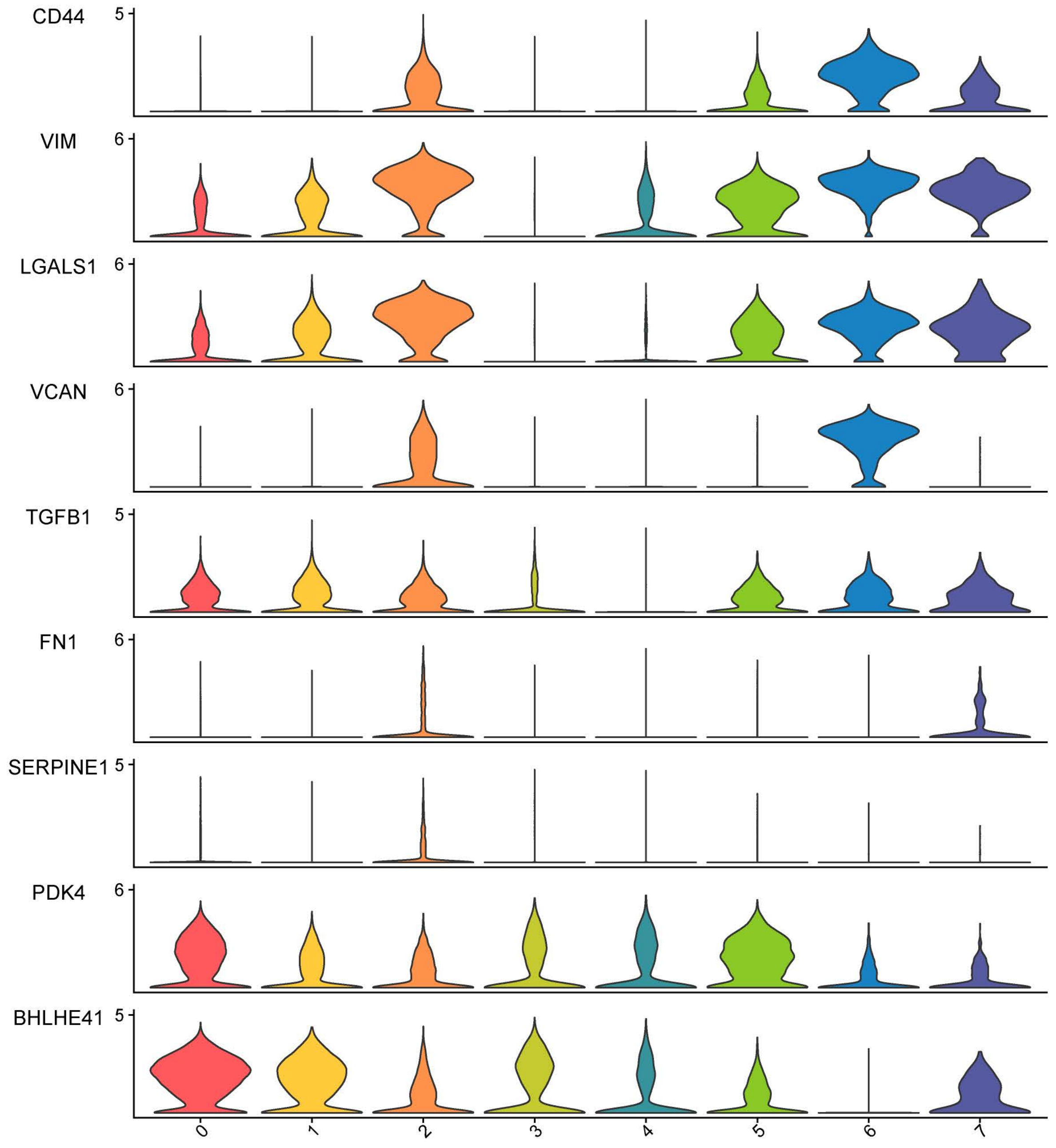


**B**

| | Sample Type | | |
|---|---|---|---|
| | WT | MUT | NB |
| Archetypal Cells (n) | 2221 | 587 | 292 |
| Proportion from Total (%) | 18.94 | 4.49 | 17.54 |

**C**

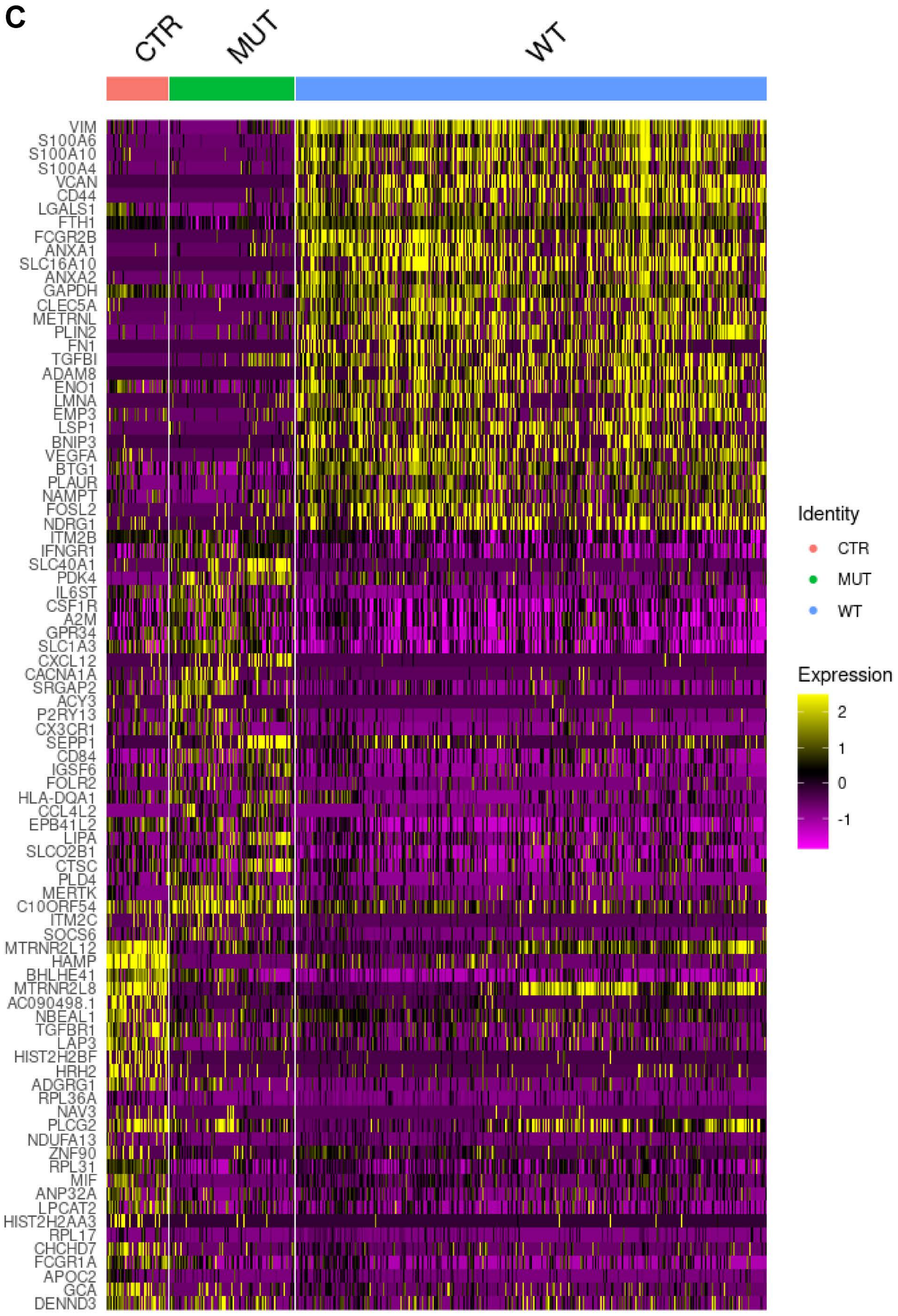


**D**

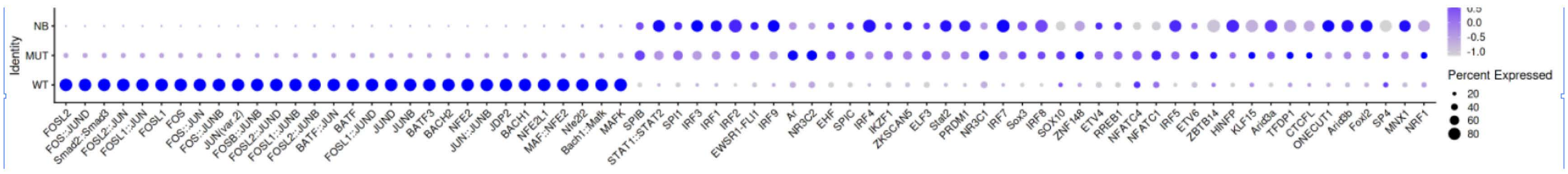


**E**

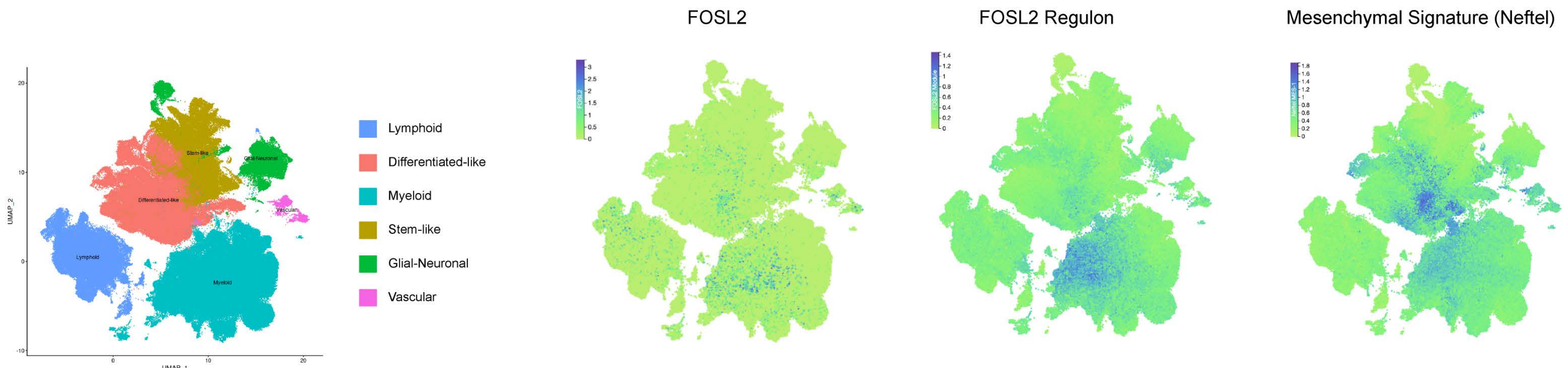


**F**

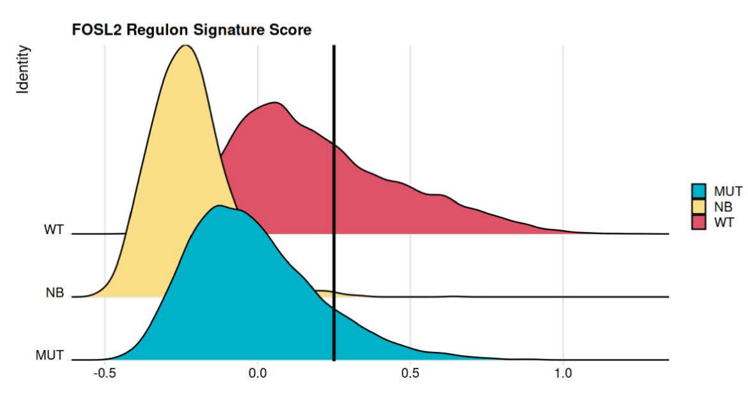


**G**

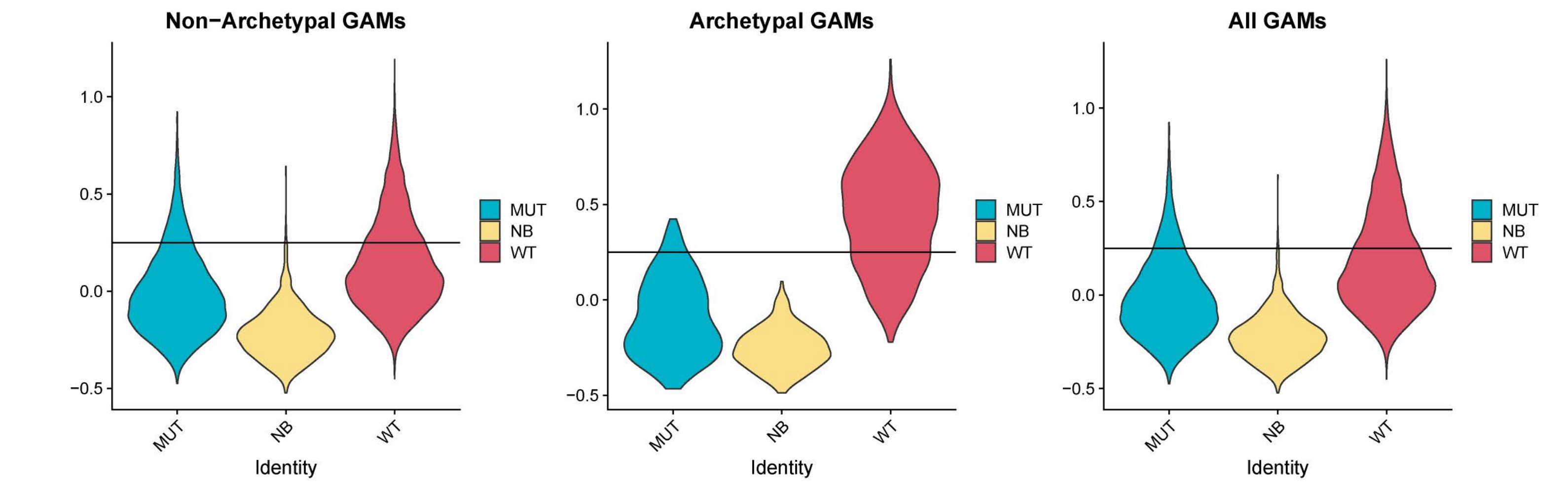

**Supplementary Figure 3**

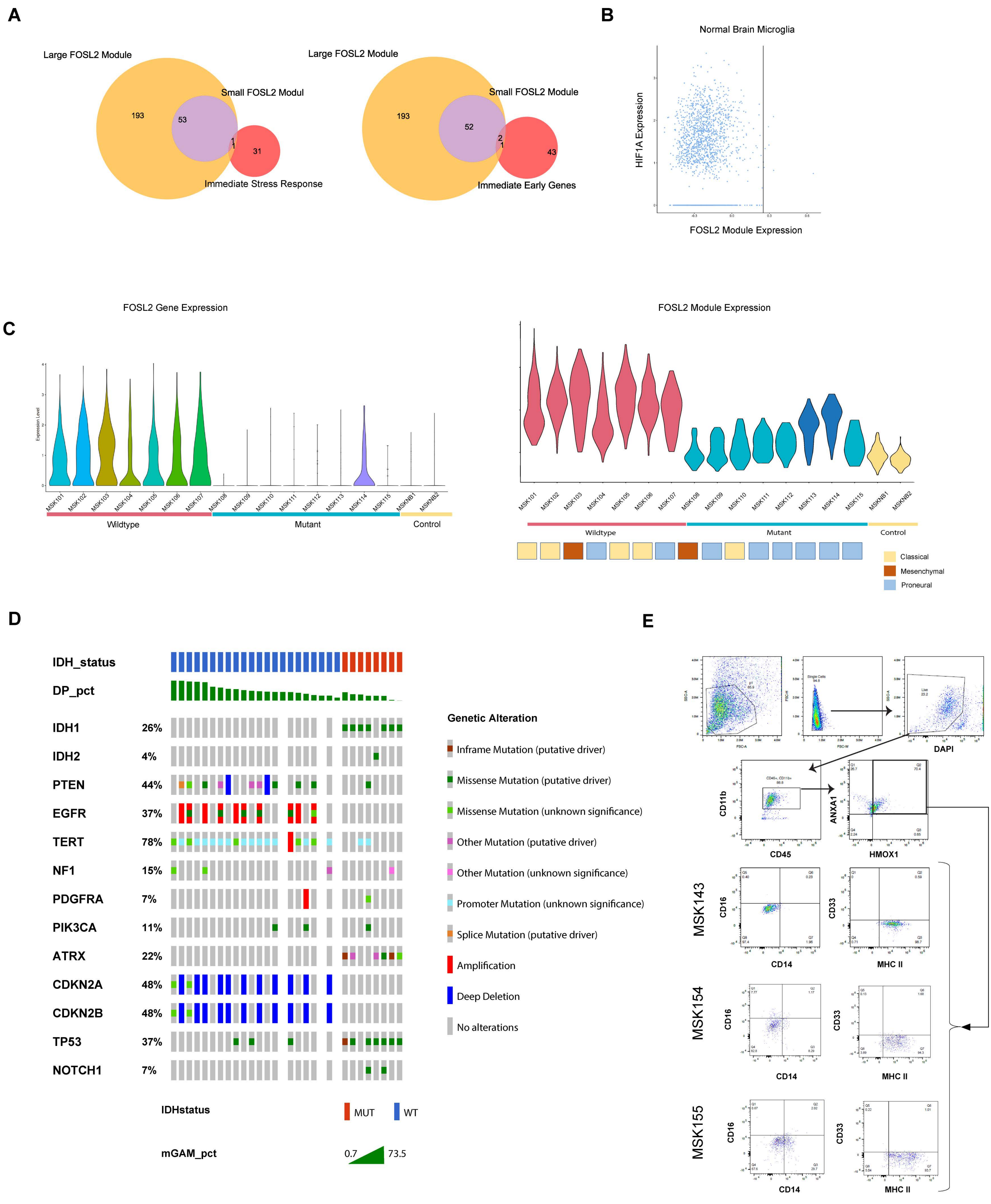

Supplementary Figure 4

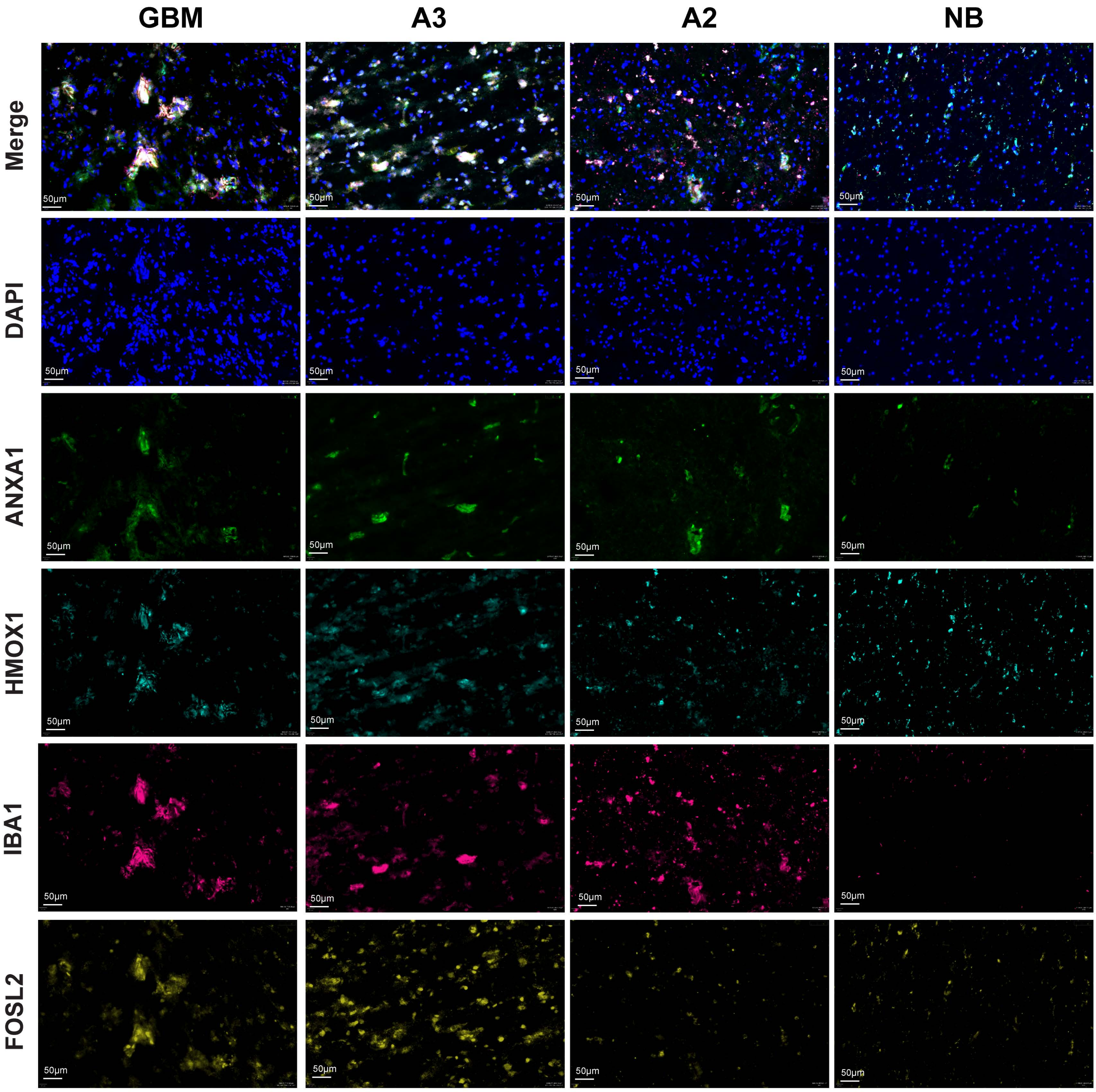

**Supplementary Figure 5**

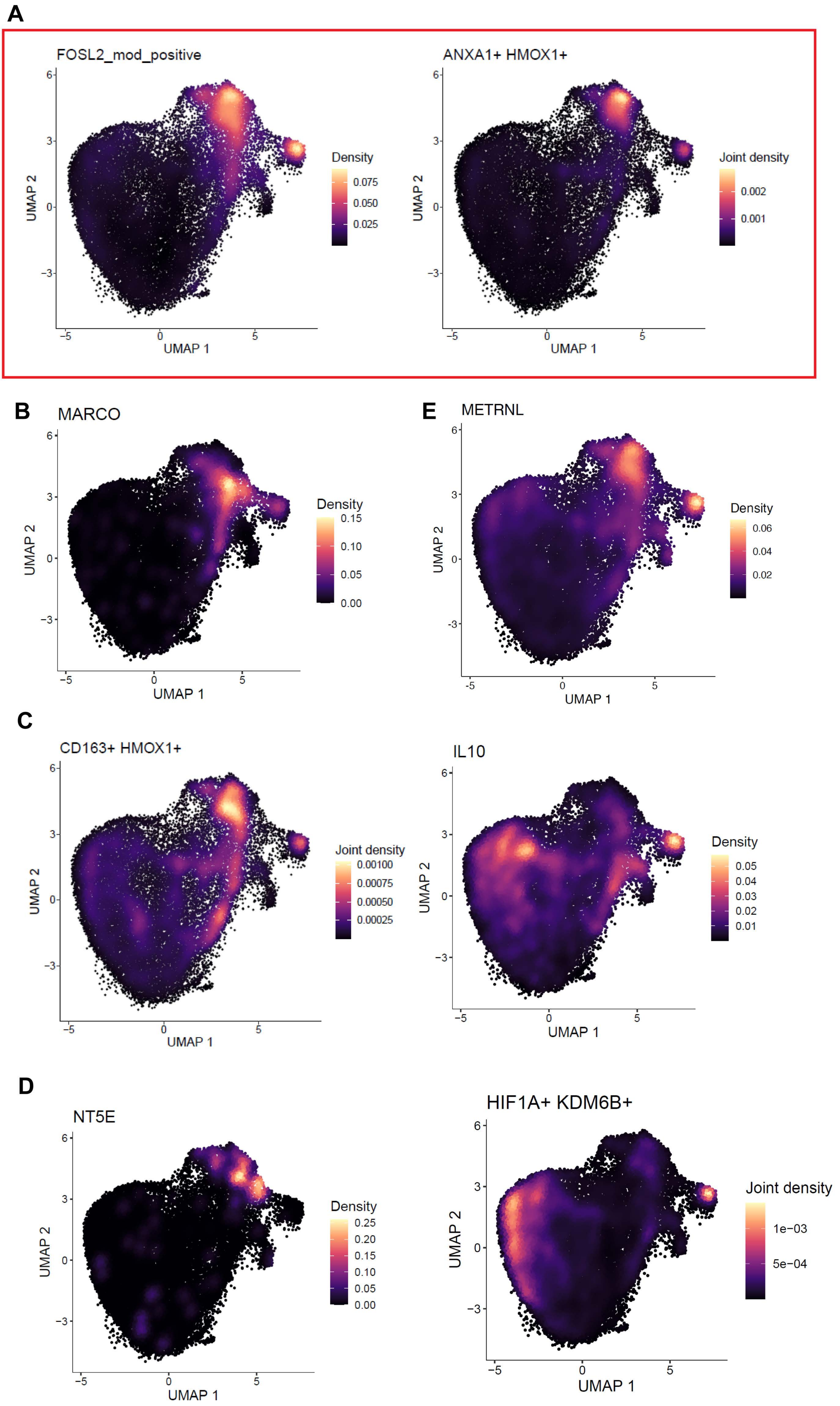

### Supplementary Figure 6

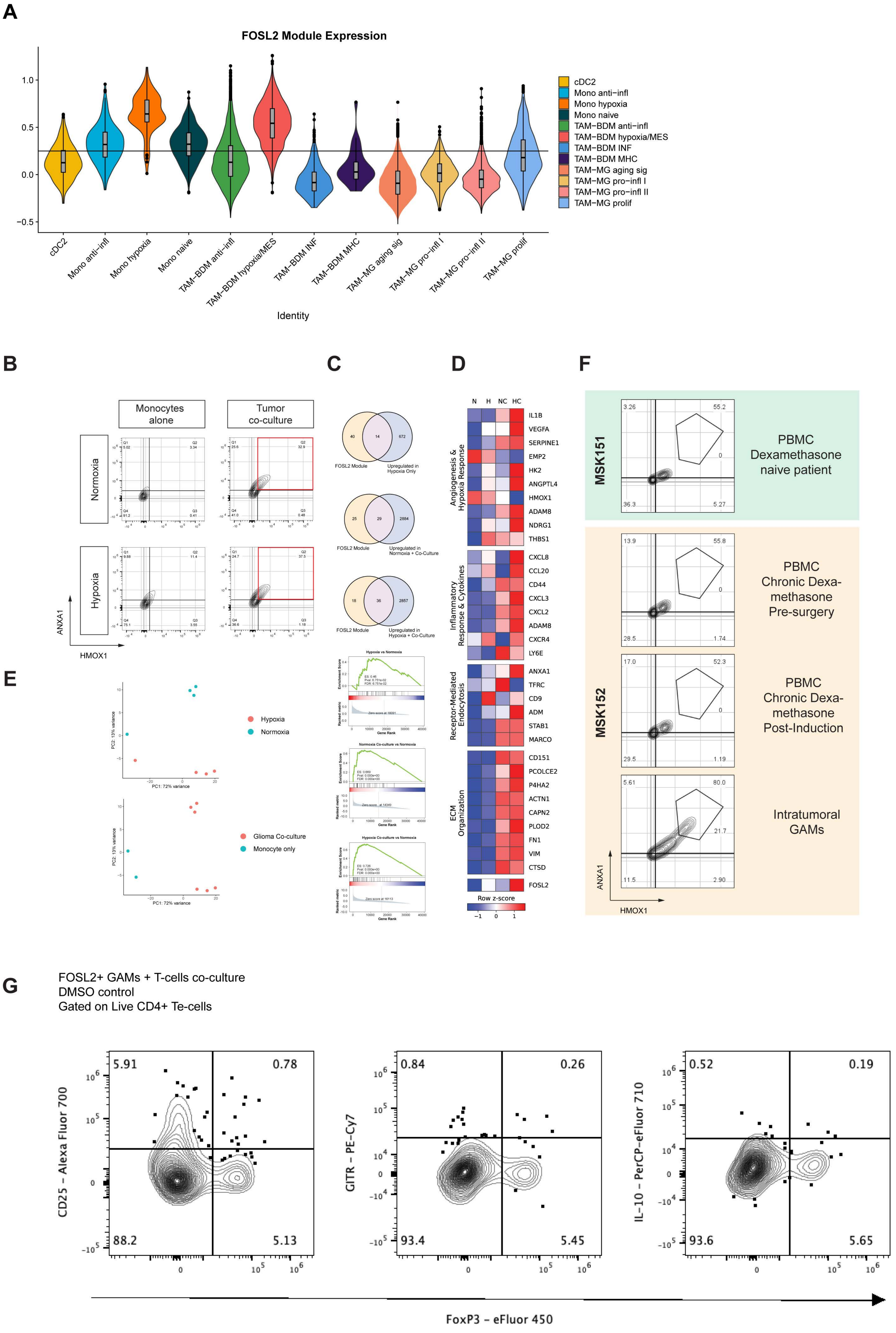

### Supplementary Figure 7

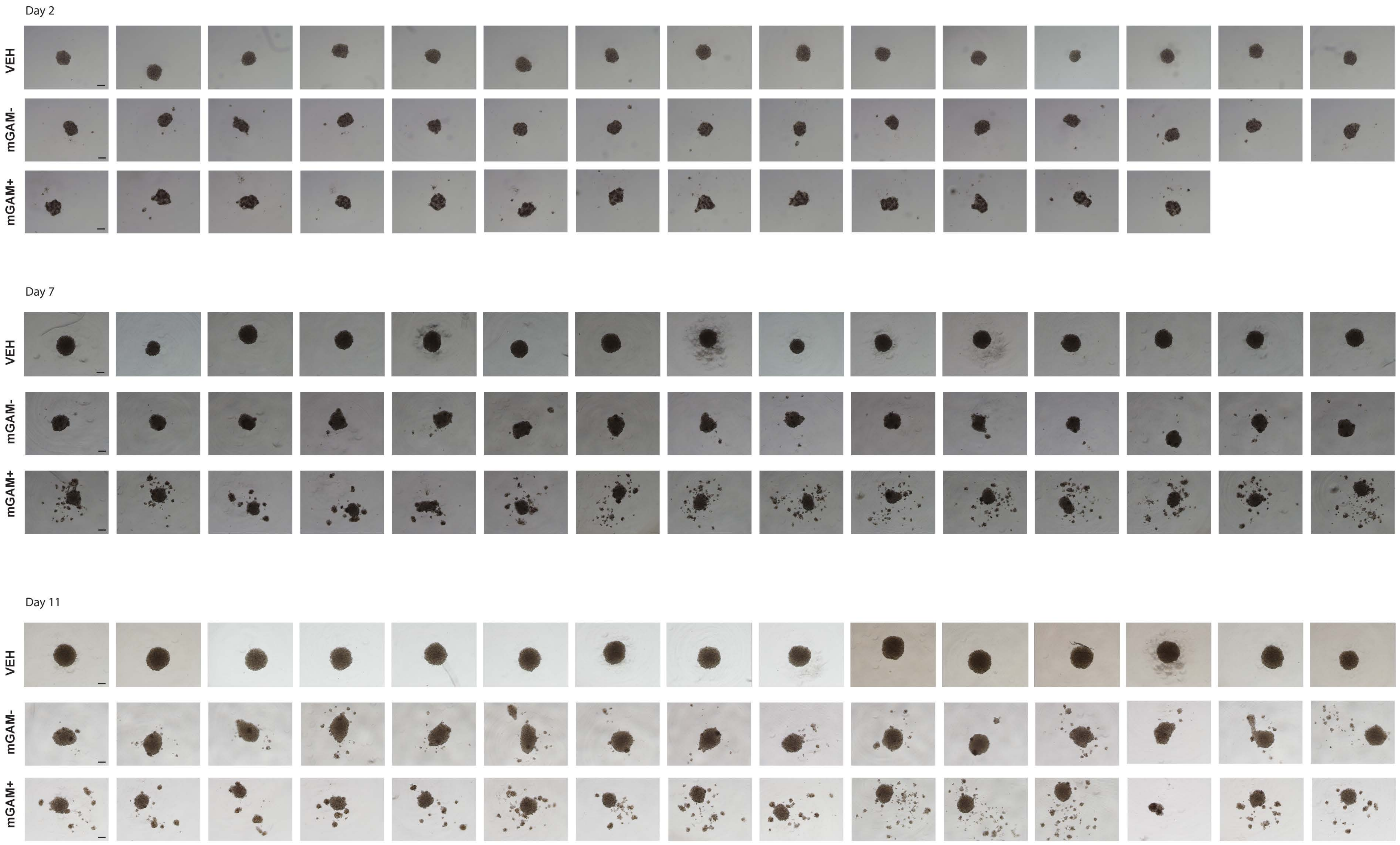

#### Supplementary Figure 8

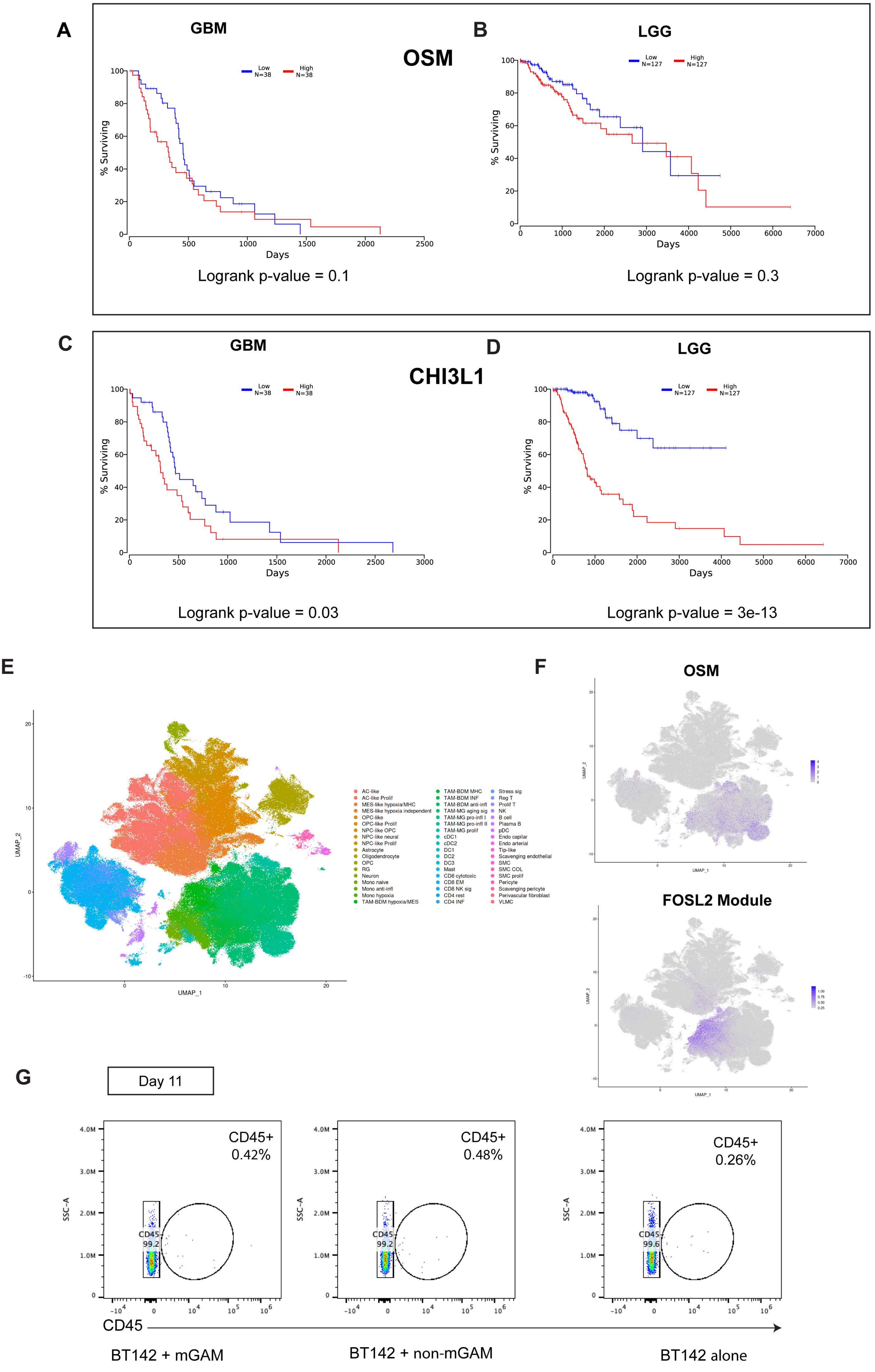
